# Desegregation of neuronal predictive processing

**DOI:** 10.1101/2024.08.05.606684

**Authors:** Bin Wang, Nicholas J Audette, David M Schneider, Johnatan Aljadeff

**Author notes:** To whom correspondence should be addressed; (J.A.).

## Abstract

Neural circuits construct internal ‘world-models’ to guide behavior. The predictive processing framework posits that neural activity signaling sensory predictions and concurrently computing prediction-errors is a signature of those internal models. Here, to understand how the brain generates predictions for complex sensorimotor signals, we investigate the emergence of high-dimensional, multi-modal predictive representations in recurrent networks. We find that robust predictive processing arises in a network with loose excitatory/inhibitory balance. Contrary to previous proposals of functionally specialized cell-types, the network exhibits desegregation of stimulus and prediction-error representations. We confirmed these model predictions by experimentally probing predictive-coding circuits using a rich stimulus-set to violate learned expectations. When constrained by data, our model further reveals and makes concrete testable experimental predictions for the distinct functional roles of excitatory and inhibitory neurons, and of neurons in different layers along a laminar hierarchy, in computing multi-modal predictions. These results together imply that in natural conditions, neural representations of internal models are highly distributed, yet structured to allow flexible readout of behaviorally-relevant information. The generality of our model advances the understanding of computation of internal models across species, by incorporating different types of predictive computations into a unified framework.

## Introduction

Predictive coding, the process of computing the expected values of sensory, motor, and other task-related quantities, is thought to be a fundamental operation of the brain [1, 2]. Violation of internally-generated expectations, known as *prediction-errors*, is an important neural signal that can be used to guide learning and synaptic plasticity [3, 4]. Signatures of predictive coding, including neural correlates of prediction-errors, were identified in multiple brain circuits, and across animal species [2, 5–7]. Two well-studied examples are motor-auditory [8–13] and visual-auditory predictions [14–16] in the mouse cortex. Previous work has proposed that a *canonical cortical microcircuit* underlies the computation of predictions and prediction-errors [2, 8, 9, 17–19]. While some predictions of this proposed microcircuit were confirmed in restricted scenarios, the hypothesis that the circuit-motif within the mouse cortex is a general mechanism for predictive processing faces a number of challenges.

First, typical experimental paradigms study predictive coding in animals trained to make a single association [12, 16, 20], while natural sensorimotor associations are typically high-dimensional (e.g., speech production [21]), as well as context-dependent [22, 23]. Little is known about how specific neural architectures in the brain learn to implement such high-dimensional computations. Second, multiple brain circuits outside of the mammalian cortex exhibit predictive coding, including subcortical circuits mediating placebo analgesia (*prediction-based* suppression of pain [24]); and motor-visual circuits in cephalopods that predict the animal’s appearance to an external observer, and use it to generate high-dimensional camouflage patterns [25]. It is not known whether these neural circuits use similar or altogether different strategies for predictive processing as the mammalian cortex. Third, predictive neural representations emerge on timescales ranging from ∼1 minute [26,27], ∼1 hour [28,29], to days [16,30]. This suggests that predictive processing is supported by plasticity mechanisms operating on a range of timescales (including short-term plasticity [31]), and that circuit reorganization may not always be required for implementing predictive computations.

The evidence that computing predictions is an integral part of sensory processing has garnered significant attention from the theoretical neuroscience community. Several studies have proposed recurrent network models that may perform these computations [32–39]. These studies typically focus on predicting a small number of stimuli within a single sensory modality. Moreover, in most cases, these models have been compared with coarse-grained neuroimaging data [7, 40]. Therefore, we lack cellular-level and circuit-level understanding of neural mechanisms underlying multi-modal predictive computations, which limits our ability to test hypotheses related to the circuit computation of predictive coding based on modern large-scale neural recordings.

Another major current gap from both experimental and modeling perspectives is predictive processing in high-dimensions: (*i*) What are the neural representations of predictable and unpredictable sensory variables in natural conditions with rich stimulus ensembles and complex inter-dependencies between stimuli [7, 41, 42]? (*ii*) What are the circuit mechanisms underlying the computation of those representations, and how are they learned? Specifically, it remains unknown whether circuits that implement predictive coding of high-dimensional stimulus ensembles are functionally segregated [2, 17, 18], and if so, whether this segregation emerges during learning or depends on molecularly distinct cell-types.

We address these questions by developing a mathematical framework to examine the predictive representations in recurrent networks processing naturalistic inputs during and after learning, and by relating this model to cellular- and population-level neural recordings. From a mechanistic perspective, we provide novel predictions into the expected degree of excitation/inhibition balance in the high-dimensional case, and shed light on the role that E/I balance plays in canceling interference between multiple learned stimuli. Moreover, since E/I balance is enforced by mechanisms operating on heterogeneous timescales [43], our model may allow incorporating seemingly unrelated phenomena into a unified framework, e.g., predictive responses that change as a result of short- or long-term plasticity. From a functional perspective, the model suggests that predictive processing of high-dimensional stimuli is robust when the representations of stimuli and of prediction-errors are *desegregated* at the cellular-level, and distributed across excitatory and inhibitory neurons. Finally, we applied our theory to examine the distinct roles played by excitatory and inhibitory neurons in generating internal predictions, and to assess the layer-specific predictive representations.

Our modeling and analysis overcomes key limitations of previous studies of predictive processing, and generates novel predictions that we confirmed here based on experimental data. Therefore, we believe that our work reveals principles of predictive processing across species and brain-regions and provides a quantitative framework for design and analysis of future experiments to decipher neural circuits underlying those computations.

## Results

### Recurrent networks that learn to generate high-dimensional predictions

We studied the neural representations formed in recurrent neural networks that perform predictive processing of multi-modal sensory and motor inputs. We focused on a typical associative training scenario where animals are presented with pairs of sensory stimuli simultaneously [9, 11, 12] or after a short delay [16]. The stimuli comprising each pair are typically of different sensory modalities (e.g., auditory-visual [16]), or involve a sensory-motor association (e.g., locomotion-auditory [12]). In this scenario, predictive computations are thought to be learned over time through synaptic-weight updates [9, 11, 12, 16, 20, 44]. Our network model consists of *N* recurrently connected neurons whose firing-rates depend nonlinearly on the input current driving their responses (Fig. 1a). The presentation of stimuli to the network is determined by the variables *x* and *y*. The strength of the input to each neuron corresponds to the components of the stimulus-specific feedforward synaptic weight vectors ***w*** and ***v***. There are *P* stimulus-pairs, and when *P* is of the same order as the number of neurons *N*, the network is said to perform *high-dimensional* predictive processing.

**Fig. 1.**
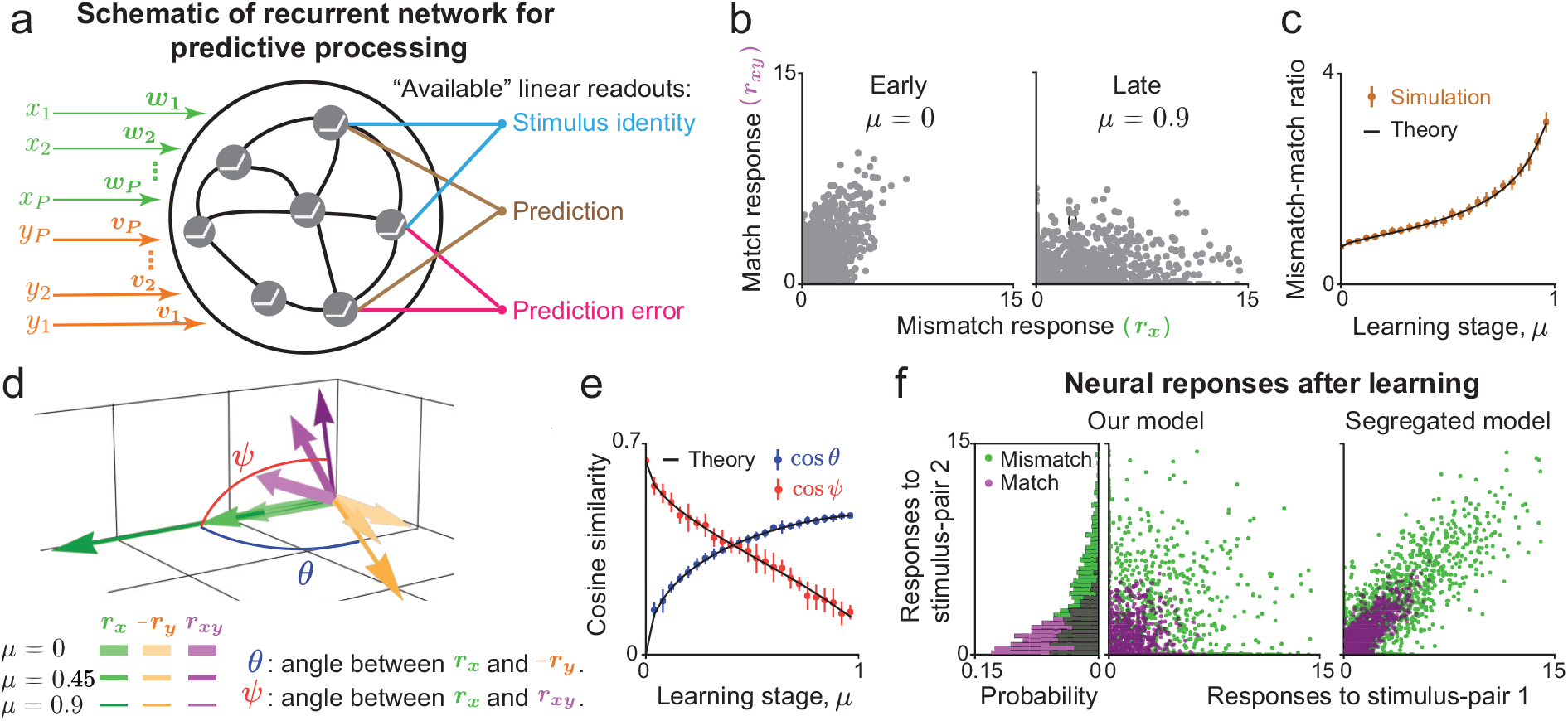
Emergence of predictive stimulus representations in a recurrent network model during learning. (a) Schematic of a recurrent network model driven by *P* pairs of stimuli (*x* and *y*). Associative training increases the correlations between the feedforward weights carrying the input signals (***w*** and ***v***). The recurrent weights jointly optimize prediction-errors and overall encoding efficiency. The neural representation formed under such optimal recurrent connectivity allows reading-out the identity of the presented stimulus; predicting a ‘missing’ stimulus; and evaluating the prediction-error. (b) Firing-rate responses of individual neurons in the match and mismatch conditions. Initially match and mismatch responses are correlated. After learning, responses are less correlated, and match responses are suppressed while the mismatch responses are amplified. (c) The ratio between average firing-rates in the mismatch and match conditions increases during learning. (d) Reduced three-dimensional neural activity space. Each vector represents the mean-subtracted firing-rate vector of neurons in the network at different conditions and stages of learning. (e) Learning leads to anti-correlation between neural responses to the stimuli *x* and *y* when presented separately (blue), and decorrelates the neural responses in the match and mismatch conditions (red), quantified by the angle between the population vectors. (f) Firing-rate responses of individual neurons to two stimulus-pairs in the match and mismatch conditions. In our model (left) there are no correlations between the responses to the two stimuli. Those responses are expected to be strongly correlated in a model in which predictive coding is functionally segregated (right). Error bars indicate standard deviations over 10 instances of the network. See Methods for additional details.

Before training, the feedforward weight vectors corresponding to each stimulus-pair are random and uncorrelated within the pair (i.e., ***w*** *·* ***v*** = 0). During training, those weights become correlated (***w*** *·* ***v*** = *µ*, with *µ* > 0), consistent with measurements of learning-induced functional reorganization of excitatory synaptic connections [45–47]. Weights of recurrent connections are chosen to minimize errors between internally generated predictions and the actual stimuli, while maximizing the overall encoding efficiency (Methods). Under these assumptions, we obtained key statistics of neural activity in the network for different stimulus inputs, at different stages of learning (Methods). The resulting neural activity allows flexibly reading-out the stimulus identity, predicting the ‘missing’ stimulus (i.e., predicting *y* based on *x*), and evaluating the prediction-error (Fig. 1a). We applied the modeling framework developed here (SI §1-2) to investigate the structure of multi-modal predictive neural representations and the circuit mechanisms supporting it.

We first examined neural responses during learning in the *match* (*x* = *y*), and *mismatch* (*x* ≠ *y*) conditions. We set *x* and *y* to be binary variables corresponding to the presence (*x, y* = 1) or absence (*x, y* = 0) of visual-auditory, visual-motor, or auditory-motor pairings [12, 13, 16, 20]. Our mathematical formalism extends to scenarios where more than two stimuli are predictive of each other, and where the inputs to the network vary continuously (e.g., running- or visual-flow-speed [20, 44]; Methods). Before associative training (*µ* = 0), most of the neurons in the network have comparable match (***r***_*xy*_) and mismatch (***r***_*x*_, ***r***_*y*_) responses (Fig. 1b). After training (*µ* = 0.9), match responses are suppressed while mismatch responses are amplified (Fig. 1b). Correspondingly, the ratio of average mismatch and match firing-rates increases (Fig. 1c), consistent with associative learning experiments [12, 16, 20]. Thus, the presence of stimulus *y* suppresses the response evoked by stimulus *x*, and generates a *prediction* (or *expectation*) of *x*. Amplified mismatch responses are interpreted as *prediction-errors* [2, 7].

During learning, the mismatch responses (***r***_*x*_, ***r***_*y*_) become *anti*-correlated (Fig. 1d,e), i.e., the presence of stimulus *y* more effectively suppresses responses to *x* alone. This anti-correlation does not appear between ***r***_*x*_ and ***r***_*y*_ of *another stimulus-pair* (Fig. S1a), suggesting that the predictive signal triggered by stimulus *y*, is *specific* to its paired stimulus *x*, consistent with Refs. [12, 48]. The *specific* suppression of responses to predictable stimuli is accompanied by a weaker, *global* gain that depends on the overall magnitude of sensory input (SI §2). Furthermore, match and mismatch neural responses *decorrelate* during learning (Fig. 1d,e), consistent with Ref. [16], suggesting that neural responses can be used to distinguish between presentation of stimulus *x* in the match or mismatch condition. Notably, Owing to the neural response nonlinearity, the match response is not a sum of the two mismatch responses (***r***_*xy*_ ≠ ***r***_*x*_ + ***r***_*y*_, Fig. 1c).

Next we examined neural responses when the network is trained with *two* stimulus-pairs (*P* = 2, Fig. 1f), making a step towards the high-dimensional scenario. [2, 17, 18, 49, 50] proposed that neurons involved in predictive processing are functionally segregated, i.e., neurons that signal prediction-error for one stimulus association tend to signal prediction-error for other associations, and similarly for ‘representation’ neurons that encode the stimulus itself. This proposal would predict a high degree of correlation between neural responses to two stimulus-pairs (Fig. 1f, right). However, we found no such correlation in our model (Fig. 1f, left). This implies, for example, that a neuron that signals prediction-error for stimulus-pair 1, may have a selective response to stimulus *x* ‘itself’ for pair 2, and raises the question of what circuit mechanisms may support this cellular-level desegregation of response types.

### Learning and stimulus dimensionality determine the properties of effective predictive processing circuits

We then investigated circuit mechanisms underlying multi-modal high-dimensional predictive processing. We decomposed the input to each neuron into feedforward and recurrent components, which respectively correspond to the actual stimulus signal and to internally generated predictions (Fig. 2a), similarly to analyses of previous experiments [2, 12, 17, 20]. To quantify the relative contribution of each component, we follow the excitatory/inhibitory (E/I) balance literature [33, 51], and define the *balance* level *B* as the ratio between the total feedforward input and the net input to each neuron, in each condition (Fig. 2a).

**Fig. 2.**
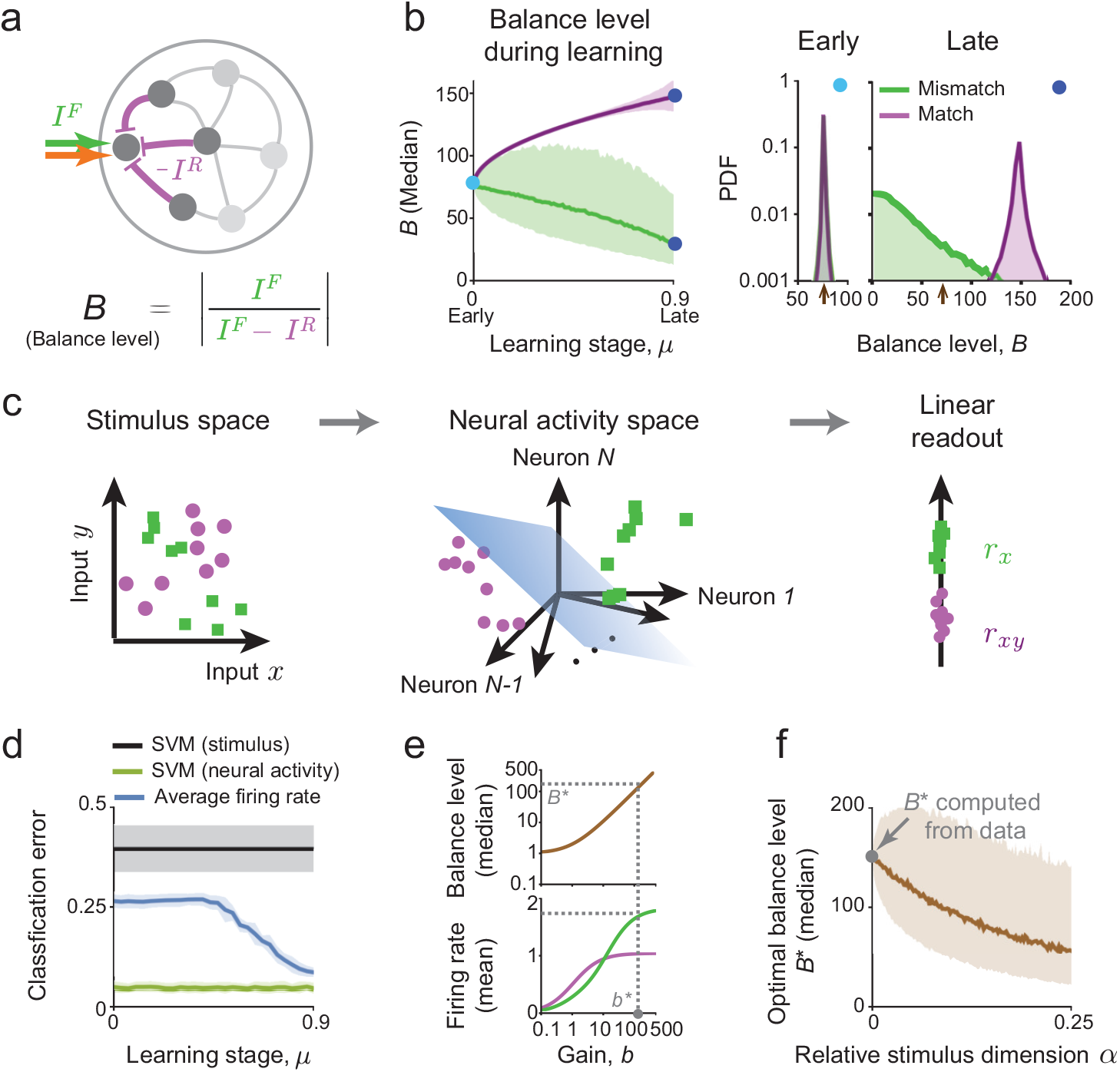
Balance between feedforward and recurrent inputs is an important mechanism supporting predictive processing. (a) The input to each neuron is decomposed into feedforward and recurrent components, which respectively correspond to the actual stimulus signal and internally generated predictions. Each neuron’s balance level *B* is the ratio between the total feedforward input and the net input (Methods). (b) The median of *B* in the match and mismatch conditions during learning (left, shaded area indicates inter-quartile range). ‘Snapshots’ of the distributions of *B* early and late in learning show that the distributions become separable in match and mismatch conditions (right). The arrows on the x-axis indicate the distribution mode early in learning. (c) Schematic showing the nonlinear transformation from the stimulus space (left) to neural activity space (center), which facilities a linear readout of relevant stimulus features (here, decoding if *x* is presented in the match/mismatch condition).(d) Error of a support vector machine classifier trained to identify the match/mismatch condition based on the input stimuli (black) and on neural responses (green). After learning, a linear classifier based on the average firing-rate (blue) performs almost as well as the optimal classifier, suggesting that functionally relevant features from all stimulus-pairs can be extracted without re-learning. (e) Illustration of the procedure to determine the optimal *b*^⋆^. The balance level *B* increases mono-tonically with the gain parameter *b* (top). Increasing *b* leads to a larger margin between match and mismatch responses (improved separability) at the cost of higher firing-rates (bottom). The optimal balance level *B*^⋆^ is determined by constraining the average firing-rate in the mismatch condition and minimizing it in the match condition. (f) Increasing the stimulus dimension leads to decrease in *B*^⋆^, i.e., a more loose balance (shaded area indicates inter-quartile range). At *α* = 0, we fit *B*^⋆^ to experimental data (Methods, [20]).

During associative learning, internally generated predictions become more accurate, facilitating more robust cancellation of the feedforward stimulus input by recurrent feedback conveying prediction signals. Thus, the overall balance level increases in the match condition but decreases in the mismatch condition (Fig. 2b, left). Notice that the balance level distributions (over neurons and stimuli) are initially similar in the match and mismatch conditions, but become significantly different in late stages of learning (Fig. 2b, right). Indeed, after learning, the mode of the balance level distribution is at *B* ≈ 0 in the mismatch condition, which explains the strong prediction-error responses.

To understand the role of balance in predictive processing, we examined its effect on the nonlinear transformation the network performs, from input stimuli to neural activity (Fig. 2c). In our model, the geometry of neural responses facilitates robust readout of prediction-errors. Specifically, while prediction-errors cannot be read-out by a linear decoder from the stimulus input, such a readout is feasible once the input is transformed into the network’s high-dimensional response (Fig. 2d). Moreover, while the prediction-error itself is stimulus-specific, the decoder that performs this computation is stimulus-independent after learning—it is simply the average firing-rate (Fig. 2d). In other words, the learned structure of neural responses enables applying the same decoder to all stimulus-pairs without ‘re-learning’.

Given the essential role of the nonlinear transformation for predictive processing, we next focused on the effect of the overall nonlinear gain parameter *b* (Methods, [34]). We found that increasing *b* leads to increases of the average match and mismatch firing-rate responses, together with a wider margin between them (Fig. 2e, top). Therefore, large *b* facilitates decoding prediction-errors, at the cost of increased overall neural activity. Motivated by this observation, and since *b* is an intrinsic network quantity that can potentially be adjusted dynamically, we sought to find an optimal value (denoted *b*^⋆^). Specifically, we constrained the average network response in the mismatch condition to be larger than a certain threshold, while requiring a minimal but nonzero average response in the match condition (Fig. 2e), consistent with reports of weak neural responses to predictable stimuli [12, 20]. The resulting *b*^⋆^ corresponds to an optimal balance level *B*^⋆^ supporting efficient encoding *and* robust decoding (Fig. 2e, bottom).

We carried out this optimization procedure for networks trained to perform predictive processing of stimulus ensembles with increasing dimensionality (i.e., increasing *α* = *P/N*), with the same firing-rate constraints chosen such that the value of *B*^⋆^ at *α* = 0 matches experimental data. We additionally assumed that an ‘over-trained’ animal learns a single stimulus-pair (i.e., *α* = 1*/N* ≈ 0). Surprisingly, we found that the optimal balance level *decreases* with *α* (Fig. 2f), independently of the stimulus statistics (Fig. S1b,c). This is because as the number of stimulus-pairs learned by the network increases, so does the interference between internally generated predictions corresponding to different stimulus-pairs (Methods). We therefore expect networks performing predictive processing in natural conditions (large *α*) to exhibit ‘loose’ balance, which minimizes the overall effect of interference arising from learning to generate a large number of internal predictions.

We used neural activity recorded from animals trained on visual-motor (V-M) [20] and auditory-motor (A-M) associations [12] to constrain our network model. Specifically, we estimated the balance levels in mouse sensory cortex by assuming that after training the neural network *in vivo* reaches the optimal balance level. In the V-M experiment [20], mice were trained to associate their running speed with the speed of visual-flow in virtual reality (Fig. 3a). The voltage of primary visual cortex neurons was intracellurlary recorded in the match and mismatch conditions. Fitting the average voltage change in the two conditions to our model gives the estimated balance level 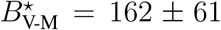. A consistent result was obtained in the A-M experiment [12], where mice were trained to press a lever and received closed-loop auditory feedback (Fig. 3b,c). Here the recording was extracellular, so fitting *B*^⋆^ relied on a slightly modified procedure (Methods).

**Fig. 3.**
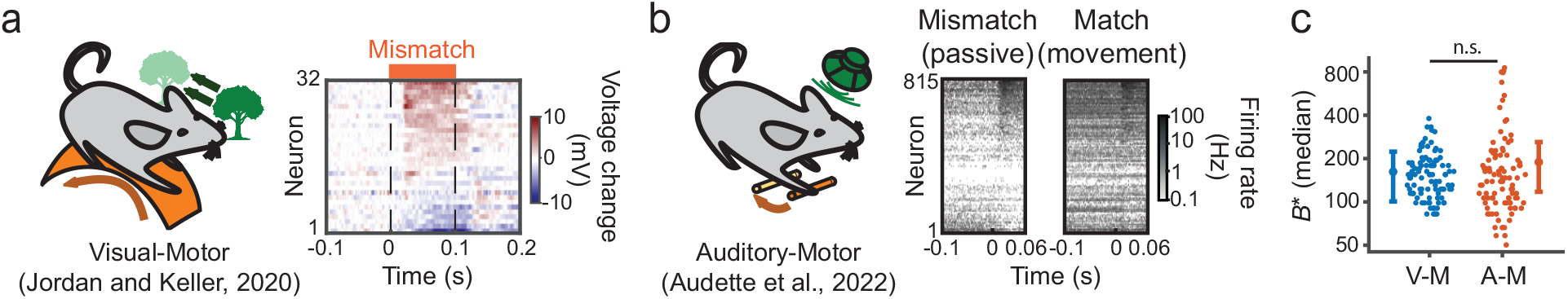
Estimating the balance level from predictive coding experiments. (a) Schematic of a learned visual-motor association between running and virtual reality visual flow [20]. Voltage levels of different neurons in primary visual cortex reveal tuning to mismatch between running speed and visual flow (prediction-errors). (b) Schematic of a learned audio-motor association between a lever press and a sound [12]. Neurons’ firing-rates reveal tuning to auditory stimuli presented without (passive, prediction-errors) and with a lever press (movement). (c) Estimating the median optimal balance level for V-M (blue) and A-M (red) experiments gives similar values. We assume that *α* = 0 based on the fact that the animals underwent extensive training on a single pair of stimuli in both experiments. Error bars are based on repeated subsampling (Methods).

It is notable that balance level estimates were consistent across animals (Fig. S2); and laboratories (Fig. 3), despite the fact that the experiments studied different brain regions and sensory modalities, using different methods. While these factors may affect the balance level to some degree, our model predicts that the balance level can decrease by up to one order of magnitude when the stimulus dimension increases (Fig. 2e, Fig. S1b,c). This prediction could be confirmed if future experiments reveal a more loose balance in animals habituated to rich sensory environments.

### Stimulus and prediction-error representations are desegregated in the model

We next investigated how different functional responses are organized within the network. Previous work postulated that two distinct neural populations exist in predictive processing circuits: (*i*) *internal representation* (*R*) neurons that ‘faithfully’ represent external sensory stimuli and encode internal predictions, and (*ii*) *prediction-error* (*PE*) neurons, which signal the difference between the actual stimulus inputs and internal predictions. Given that neurons selective to these signals also exist in our network model, we wondered whether they form functionally segregated populations. We adopted classification criteria used in experimental work (Methods, [2, 48]): *R* neurons are those which respond strongly and similarly in match and mismatch conditions, while *PE* neurons are those which respond strongly in the mismatch condition but weakly in the match condition (Fig. 4a).

**Fig. 4.**
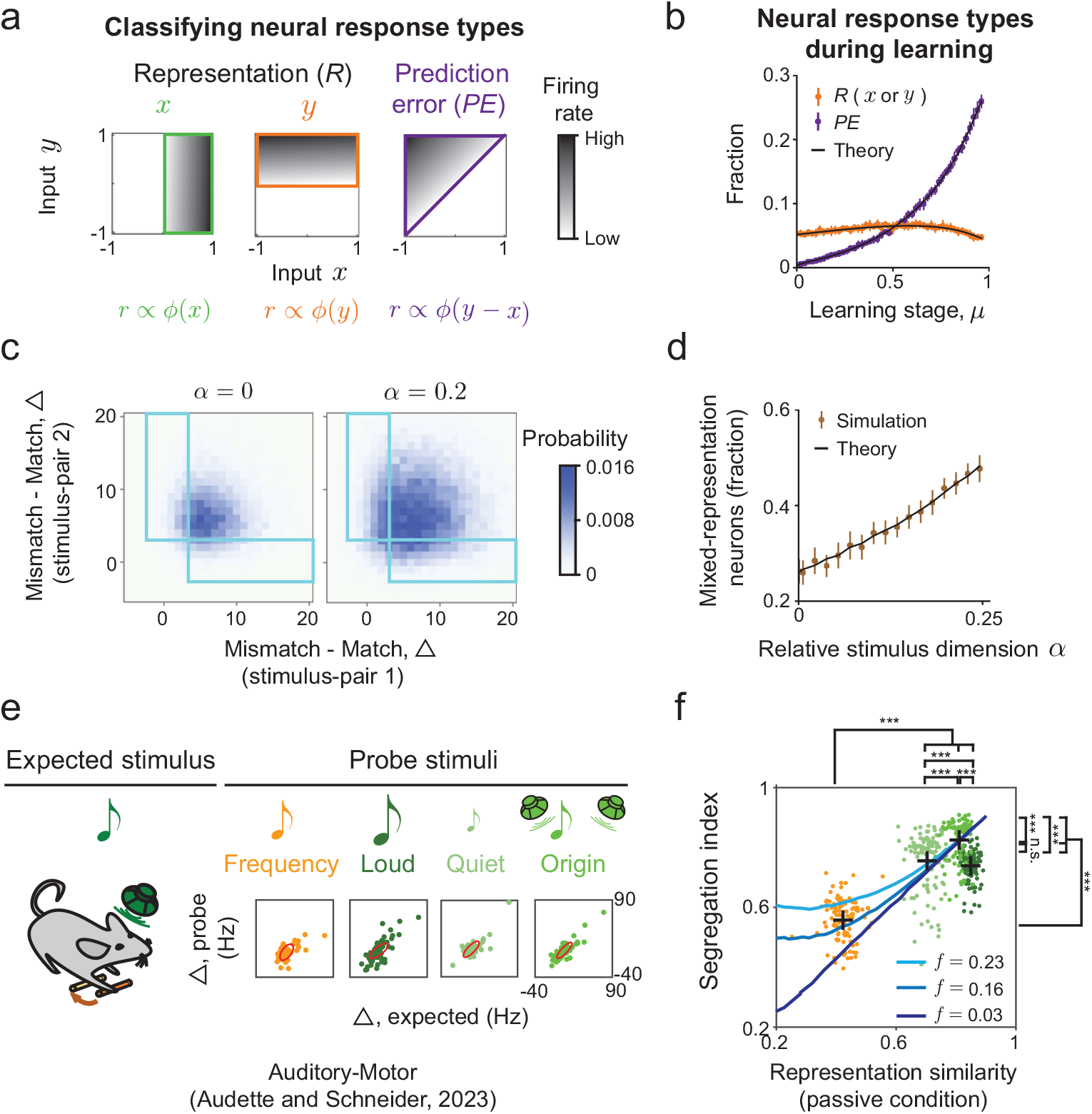
Desegregated stimulus and error representations in networks performing high-dimensional predictive processing. (a) Schematic of typical tuning profiles of different functional cell-types to the stimuli *x* and *y*. (b) Fraction of representation (*R*) and prediction-error (*PE*) neurons in the model at different learning stages. Error bars indicate standard deviation over 10 instances of the network. (c) Joint distribution of individual neurons’ Δ values, the difference between mismatch and match responses to two specific stimulus-pairs. Only neurons responsive to both stimulus-pairs are included in the distribution (Methods). Mixed representation neurons have significantly different Δ values for the two stimulus-pairs, i.e., they are in the blue rectangular regions. As the stimulus dimension (*α*) increases, more neurons have a mixed representation of stimuli and prediction-errors. (d) The fraction of mixed representation neurons increases as stimulus dimension increases. Error bars indicate standard deviations over 200 instances of the network. The threshold for defining response types was based on neural activity statistics at *α* = 0 and was used for all values of *α* (SI §5). (e) Evaluating the segregation of stimulus and prediction-error representations based on neural recordings during a learned auditory-motor association. Shown are the Δ values of stimulus-responsive neurons for the expected sound and each probe type (colors). Red ellipses indicate the spread of data. The length and direction of major and minor axes correspond to the amplitude and direction of the two leading principal components. (f) Segregation index as a function of representation similarity for different pairs of expected and probe sounds. Colored points correspond to subsamples of the data, and crosses correspond to the average for each probe type (Methods). Experimental data is compared with equivalent quantities from the model, obtained by varying the sparsity of responses in the model (*f*, see SI §3).

Based on these criteria, we first computed the fractions of *R* and *PE* neurons when the network learns a single stimulus association (*P* = 1, Fig. 4b). As training progresses, the fraction of *PE* neurons increases significantly, consistent with experiments [16, 52], and with the notion that the network learns to ‘recognize’ the stimulus pairing. This result is independent of the classification criterion (Fig. S3). The fraction of *R* neurons remains unchanged (Fig. 4b), though we note that the trend does depend on the criterion (Fig. S3).

We next asked how neurons responded to more complex stimulus ensembles, specifically for two learned pairs of stimuli. The hypothesis that predictive processing is segregated [2, 18] asserts that if a neuron is a *PE* neuron for stimulus-pair 1, and if it is active during presentation of stimuli from pair 2, it will likely be categorized as a *PE* neuron with respect to those stimuli too. To test this hypothesis, we computed the joint distribution of neural responses in the four relevant conditions (mismatch/match, stimulus-pair 1/2) and categorized each neuron as *R* or *PE*, separately for each stimulus-pair (Methods). We started with the low-dimensional scenario, where the two stimulus-pairs in question are the only stimuli learned by the network (*P* = 2, *α* = *P/N* ≈ 0). Surprisingly, under the data-constrained parameters, although many neurons belong to the same functional type with respect to the two stimulus-pairs, approximately 25% of neurons are in fact *‘mixed’*: they are classified as having different functional types (Fig. 4c, left).

Furthermore, increasing the dimension of the stimulus the network learns, leads to a twofold increase in the fraction of mixed neurons (Fig. 4c,d). Intuitively, loose balance between high-dimensional feedforward and recurrent inputs leads to a broad balance level distribution across the network (Fig. S4a). That broad distribution, in turn, affords each neuron flexibility to encode different features for different stimulus-pairs. The fraction of mixed neurons shown in Fig. 4d corresponds to two *specific* stimulus-pairs. When we considered instead the entire learned stimulus-set, *most* of the neurons are mixed with respect to at least two pairs (Fig. S4b). Thus, contrary to the previous hypothesis [2], neurons with mixed representations of stimuli and predictions are common in the network model, especially in high-dimensional scenarios.

### Experimental evidence for desegregated predictive representations

We then turned to testing this key prediction of our network model, by looking for signatures of mixed representations of predictions and stimuli in experimental data. In our recent work, we recorded primary auditory cortex responses in mice that were trained to associate a simple behavior, pressing a lever, with a simple outcome, a predictable tone [13]. Following extensive training, we made extracellular recordings from auditory cortex while animals were presented with *probe* auditory stimuli that differed from the *expected* stimulus along a variety of different dimensions, and while animals either pressed the lever or heard the tone passively (Fig. 4e).

Here we analyzed this data as follows. For each neuron, we computed the difference (Δ) between the mismatch (passive: sound only) and match (active: lever press + sound) neural responses (Fig. 4e, bottom), similar to our analysis of the neural activity in the model (Fig. 4c). Note that for each of the four probe sounds, ‘match’ corresponds to a lever press paired with the probe sound, while ‘mismatch’ corresponds to responses following the probe sound without lever press. We expected Δ values of mixed neurons to lie in the upper left or lower right corners of the plot (similarly to Fig. 4c, blue rectangles). This would correspond to neurons with match and mismatch responses that are similar for the expected sound but differ for the probe sound, or vice versa.

We quantified the degree of mixing, or desegregation of the predictive representation, by computing the Pearson correlation coefficient of the Δ values corresponding to the expected sound and each probe sound separately (Fig. 4e). We defined this coefficient as the *segregation index*, which is close to 1 if the Δ’s are strongly correlated between the two stimulus-pairs (expected, probe). A segregation index close to 0 means that the representations of stimuli and predictions are ‘maximally mixed’. We additionally computed representation similarity between the expected and probe sounds, as the correlation between neural responses to those stimuli. Crucially, representation similarity was based on neural responses in a separate experi-mental window during which sounds were presented passively, not following a lever press [13]. If neurons are segregated into two functional classes, the segregation index should be close to 1 irrespective of the representation similarity. By contrast, we found that the segregation index depends strongly on the representation similarity (Fig. 4f). Specifically, when the expected and probe sounds are similar (Fig. 4e,f, green shades), the segregation index is close to 1, though a random subsampling analysis indicates a statistically significant effect of the representation similarity on the segregation index. When the probe differs from the expected sound more substantially (Fig. 4e,f, orange), the segregation index drops to ∼ 0.5. This relation between representation similarity and degree of segregation is consistent with the prediction of our model, with an appropriate level of coding sparsity (Fig. 4f). The significant dependence of the segregation index on the representation similarity, and the fact that the segregation index is substantially smaller than 1, suggest that predictive processing is mixed in the mouse auditory cortex. A similar relationship was found when we used the ‘complementary’ mismatch response to compute the Δ’s, i.e., based on the neural response to a lever press with no sound, rather than a sound with no lever press (Fig. S5).

We note that the analysis presented here is an indirect test of the model prediction that predictive representations are mixed. Indeed, the desegregation in the model involves two learned stimulus-pairs (Fig. 4c), while in the experiment the animal was only trained on the expected sound. Nevertheless, the decreased segregation index we found for probe sounds markedly different from the expected sound provides strong evidence against the notion that predictive processing circuit is functionally segregated into separate neural populations. Our model provides a framework to generate hypothesis that could be tested more directly in future experiments.

### Predictive processing in excitatory–inhibitory networks

Thus far we have focused on relating neural responses in the model to measurements of excitatory neurons’ activity [12, 13, 16]. Each neuron’s projections in our network could be both excitatory (E) and inhibitory (I), so it does not obey Dale’s law. Given the growing literature on the role of inhibitory neurons in computing predictions [35, 36], we sought to link our model to experiments more tightly by extending it to a network with separate E and I neurons. We did so by requiring that the activity of E neurons in the E/I network matched exactly that of neurons in the original model. This guarantees that the E neurons possess the predictive coding properties we studied so far, and opens the door to study the functional role of I neurons. The connectivity in the E/I network has four components, corresponding to synapses to and from E and I neurons (Fig. 5). We used non-negative matrix factorization to ‘solve’ for those components (Methods, [53, 54]). The balance level *B* defined previously based on feedforward and recurrent inputs (Fig. 2), is equal to the stimulus-specific component of the E/I balance in the E/I networks (SI §4).

**Fig. 5.**
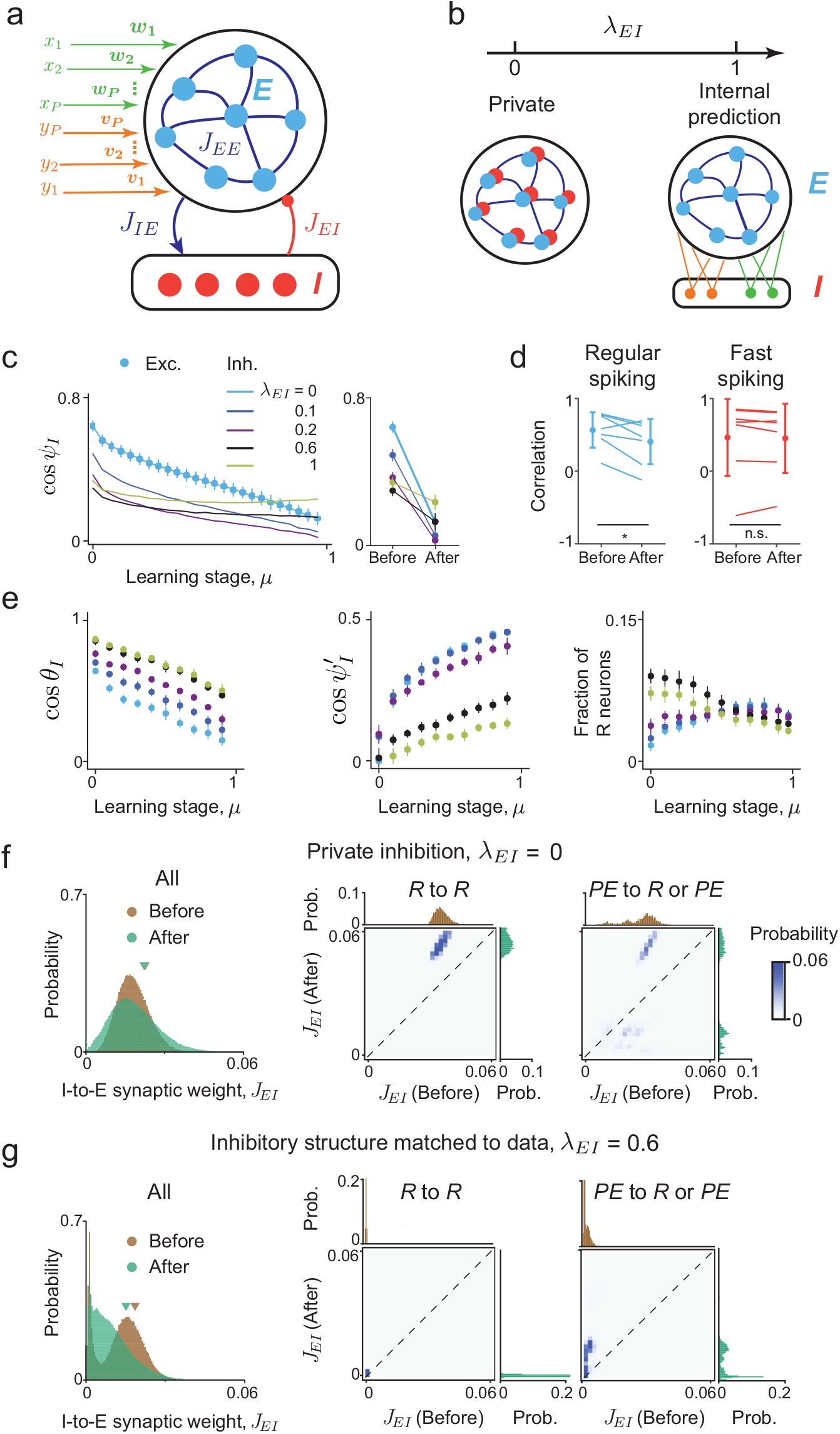
A data-constrained excitatory/inhibitory model suggests that internally-generated predictions are distributed across the network. (a) Schematic of the E/I network with separate connectivity components. Excitatory neurons receive external inputs, and their activity is constrained to equal that of neurons in our original model. (b) A family of E/I networks that satisfy the desired constraints, identified based on non-negative matrix factorization. Solutions are parameterized by *λ*_*EI*_, which interpolates between ‘private’ inhibition and inhibition that signals ‘internal predictions’. Varying *λ*_*EI*_ gives different patterns of inhibitory responses and connectivity structures. (c) The cosine similarity (cos *ψ*_*I*_) between the match and mismatch inhibitory responses to stimulus *x* (***r***_*xy*_, ***r***_*x*_), for different values of *µ* and *λ*_*EI*_ (left). Comparing cos *ψ*_*I*_ before and after learning (right) allowed us to link inhibitory connectivity structure to inhibitory representations. (d) Analogous correlation between population responses, computed separately for regular-spiking (RS) and fast-spiking (FS) neurons from Ref. [12]. Each point represents data from one animal. The mean and standard deviation of the correlations across animals are also shown. RS neurons significantly decorrelate during learning, while FS neurons’ correlation does not change. Correlations of RS and FS neurons after learning are similar. (e) The angle *θ*_*I*_ (left) between inhibitory population responses to the paired stimuli in the mismatch conditions (***r***_*x*_, −***r***_*y*_), and the angle 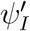 (center) between match and mismatch inhibitory population responses to stimulus *y* (***r***_*xy*_, ***r***_*y*_). Angles are shown as a function of *µ* and *λ*, leading to experimentally testable predictions pertaining to inhibitory representations. Fraction of inhibitory *R* neurons (right) as a function of *µ* and *λ*_*EI*_. For the experimentally constrained parameter *λ*_*EI*_ = 0.6, this fraction decreases for inhibitory neurons (black), while it does not change significantly for excitatory neurons (*λ*_*EI*_ = 0, blue, Fig. 4b). (f) Synaptic weight distribution of all I-to-E connections before and after learning, when *λ*_*EI*_ = 0 (left), and for pairs of E and I neurons belonging to specific functional classes (*R* to *R*, middle; *PE* to *R* or *PE*, right). Learning broadens the overall synaptic weight distribution and potentiates the inhibitory connections between inhibitory *R* neurons. (g) Same as (f), when inhibitory structure is matched to data (*λ*_*EI*_ = 0.6). Here learning sparsifies and depresses inhibitory connections. Connections between *R* neurons remain very small throughout learning. Surprisingly, connections from inhibitory *PE* neurons are strongly potentiated.

The aforementioned mathematical procedure did not yield a *unique* connectivity structure. Rather, we found a one-parameter family of connectivity structures that all meet those constraints. This parameter, denoted *λ*_*EI*_, interpolates between two extremes of structured E/I connectivity (Fig. 5b). In one extreme (*λ*_*EI*_ = 0), inhibition is *‘private’*: Each ‘parent’ E neuron projects to a single ‘daughter’ I neuron with equal activity. This has been an implicit assumption of previous predictive coding models with lateral inhibition [38, 55]. In the opposite extreme (*λ*_*EI*_ = 1), each I neuron receives a large number of excitatory inputs and signals an *‘internal prediction’* of one stimulus learned by the network, similar to previous models with segregated neural populations [35, 36]. We investigated the continuum of inhibitory representations between these extremes using the same approach applied to E neurons (Fig. 1b-e, Fig. 4b). We started with the alignment of inhibitory responses to stimulus *x* in the match (***r***_*xy*_) and mismatch (***r***_*x*_) conditions, at different learning stages (Fig. 5c). Before learning (*µ* = 0), increasing *λ*_*EI*_ leads to a marked decrease in the alignment of inhibitory responses. After learning (*µ* ≈ 1), increasing *λ*_*EI*_ leads to a non-monotonic effect on alignment. Intriguingly, for *λ*_*EI*_ = 1, after learning, the alignment of I responses in the two conditions is larger than that of E responses (Fig. 5c, compare green and black for *µ* = 1).

These properties allowed us to estimate the parameter *λ*_*EI*_ based on empirical measurements of regular-spiking (RS, putative excitatory) and fast-spiking (FS, putative inhibitory) neurons. To achieve that, we computed the correlation between auditory cortex match and mismatch responses, separately for RS and FS neurons recorded in Ref. [12], and then compared those correlations to the model before and after learning (Fig. 5d). Specifically, The pairing between movement and a probe sound (not presented during training) was regarded as *before*-learning and the pairing between movement and the expected sound as *after*-learning (Methods). This correlation decreased significantly during learning for RS neurons, consistent with the change in the model’s E population responses (Fig. 5c, blue circles). By contrast, correlation of FS population responses did not change significantly during learning, which rules out small values of *λ*_*EI*_. Moreover, the correlation value after learning was similar for RS and FS neurons, which rules out large values of *λ*_*EI*_. Taken together, our analysis suggests that an intermediate value of *λ*_*EI*_ ≈ 0.6 best captures the experimental observations, consistent with the suggestion of ‘promiscuous’ inhibitory connections mediating suppression of expected stimuli [11].

Given this experimentally-constrained value (*λ*_*EI*_ = 0.6), our theory generates testable predictions for inhibitory predictive representations. First, we expect that anti-alignment of mismatch I responses (*x*-only, *y*-only) is significantly weaker when compared to anti-alignment of E responses in the same conditions (Fig. 5e, left; Fig. 1d,e). Second, we predict large correlations between inhibitory responses in the match and *y*-only mismatch conditions (Fig. 5e, middle), when compared with E responses. The asymmetry of ***r***_*x*_ *·* ***r***_*xy*_ and ***r***_*y*_ *·* ***r***_*xy*_ overlaps in the model may in the future be related to distinct functional responses of inhibitory neuron subtypes [23, 56]. Third, the fraction of I neurons with *R* responses decreases moderately during learning, compared to E neurons. We note however that the fraction of E neurons with *R* responses shows moderate dependence on the threshold, particularly before learning (Fig. S6), which may make it challenging to detect differences in fractions of neurons with *R* responses between E and I neurons.

Previous work on predictive coding suggested that associative learning enhances top-down inhibitory projections from outside the local circuit [2, 16], which cancels bottom-up excitation and suppresses neural responses in the match condition. We therefore wondered what changes in inhibitory connectivity during learning lead to stimulus-specific suppression of neural activity in our E/I network model. One option is that inhibitory connections that predict the stimulus are strengthened [2]. Alternatively, inhibition could undergo more subtle reorganization such that inhibitory signals are distributed differently before and after learning.

We calculated the distribution of I-to-E synaptic weights before and after learning in the family of E/I network models. When inhibition is private (*λ*_*EI*_ = 0), this distribution broadens during learning (Fig. 5f). Examining the change in synaptic weights conditioned on the functional cell-type of pre- and post-synaptic neurons (*R* or *PE*), suggests that stimulus-specific suppression of E responses arises from potentiated I synapses from neurons ‘faithfully’ representing the stimulus. In other words, when inhibition is private, the predictive signal arises in part due to strengthened projections from inhibitory *R* neurons to excitatory neurons (Fig. S7). By contrast, when inhibitory structure was matched to experimental data (*λ*_*EI*_ = 0.6), learning leads to overall sparsification of I connections (Fig. 5g). Interestingly, here *R*-to-*R* connections can be either potentiated or depressed, unlike the *λ*_*EI*_ = 0 case (compare middle panel of Fig. 5f,g). Moreover, when *λ*_*EI*_ = 0.6, inhibitory connections originating from *PE* neurons that are initially very weak get strongly potentiated.

Together, our results suggest that (*i*) Predictive processing is learned without large increases of the average inhibitory connection strength. This was also seen for other values of *λ*_*EI*_ (Fig. S8). (*ii*) The ‘strategy’ for learning predictive processing can differ substantially, and depends on the underlying circuit structure (different values of *λ*_*EI*_ in the model). (*iii*) When inhibitory structure is matched to data, the ‘internal model’ is highly distributed and, surprisingly, arises in part from potentiated connections from inhibitory neurons signaling prediction-error. Another signature of this distributed strategy is the *decrease* of total inhibitory input to each excitatory neuron during learning (Fig. S8), which suggests that predictions are primarily computed by recurrent circuitry rather than directly from top-down inputs.

### Predictive representations in hierarchical neural networks

Sensory brain regions are known to have a laminar structure, and distinct layer-specific response characteristics in associative learning tasks [17, 20, 57]. In the context of the task involving sensorimotor predictions, it has been suggested that motor-related input originates from motor regions and first enters the primary sensory region via deep layers (L5/6) [2, 58–60]. On the other hand, the bottom-up sensory-related inputs first enter the primary sensory region via L4, which further projects to L2/3 where the bottom-up and top-down inputs are integrated and processed [61, 62]. To investigate the effects of the laminar structure on predictive processing, we extended the recurrent network model which has a single-*module* and no hierarchical structure, to a network model with three recurrently interconnected *modules* (Fig. 6). During associative learning, the network receives paired multimodal inputs. Crucially, the first module (M1) of the network receives inputs from one modality, and the last module (M3) receives inputs from the other modality (Fig. 6a). Differently from previous work [17, 39, 63], each module in our network computes bidirectional predictions, corresponding to inputs from the level above and below it in the hierarchy. For example, M2 computes predictions of activity in M1 and M3. Our hierarchical model can also be applied to cross-modal processing performed by distinct brain regions that exchange predictive signals bidirectionally (e.g., auditory and visual cortices, [16]), beyond laminar organization within a single brain-region.

**Fig. 6.**
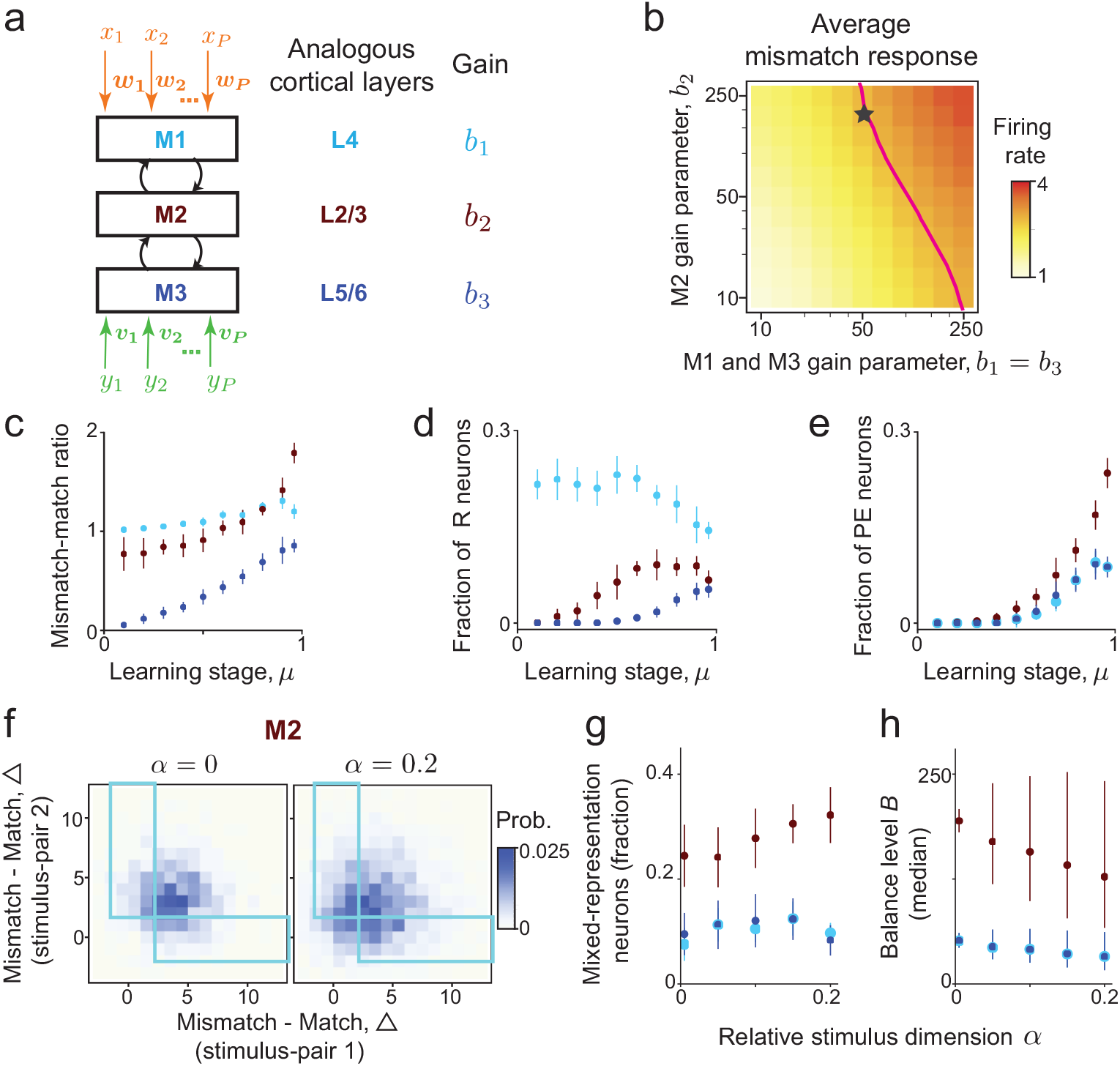
Representations of stimuli and prediction errors vary across a hierarchical network. (a) Hierarchical network for predictive processing with three modules. M1 and M3 receive stimulus *x* and *y* input, respectively. (b) The average *x*-only mismatch response increases with the module-specific gain parameters *b*_1,2,3_. Line: mismatch response amplitude used to constrain *b*_1,2,3_. Star: parameter values further constrained based on the fraction of prediction error neurons in M2, used in panels (c-h). (c) The ratio between the average firing-rates in the *x*-only mismatch and match conditions increases during learning. The increase is most prominent in M2. (d) The fraction of *x* representation (R) neurons at different learning stages. Differences between the modules diminish with *µ*. (e) The fraction of prediction error (PE) neurons at different learning stages. (f) Joint distribution of individual neurons’ Δ values, defined the difference between mismatch and match responses to two specific stimulus-pairs in M2. Mixed representation neurons are in the blue rectangular regions. The fraction mixed representation neurons increases with the stimulus dimension *α*. (g, h) Effects of increasing the stimulus dimension *α*. (g) The fraction of mixed representation neurons increases with *α* in M2, and remains constant in M1 and M3. (h) The median balance level decreases with *α* in M2 and remains approximately constant in M1 and M3. Error bars indicate inter-quartile range.

We first studied the effects of module-specific gain parameters. After learning, the average mismatch responses increase monotonically with *b*_1_ and *b*_2_ (Fig. 6b). We constrained the average mismatch response to be larger than certain threshold value and minimized the match responses for each module. Doing so gave a continuous set of parameter combinations for which the network satisfies those constraints (Fig. 6b, magenta line). We fixed *b*_2_ such that the fraction of prediction error neurons in M2 after learning is similar to the fraction in the single-module model (Fig. 3b), which also fixes *b*_1_ and *b*_3_ (Fig. 6b, star). With these constrained parameters, we assessed how associative learning shapes neural representations across different modules.

In the *x*-only mismatch condition (*x* = 1, *y* = 0), the overall mismatch responses increase during learning, with notable module-specific differences (Fig. 6c): neurons in M1 that directly receives the *x*-stimulus input have remarkably similar responses in the match and mismatch conditions throughout learning. In contrast, neurons in M3 respond predominately to stimulus *y* but gradually become tuned to stimulus *x* as learning progresses. Neurons in M2 exhibit the largest mismatch-match response ratio and develop the most significant prediction error responses after learning.

Next we categorized neurons along the hierarchy into functional cell-types. Before learning, neurons activated by the stimulus *x* independently of *y* (i.e., *x* representation neurons) are concentrated in M1–the module receiving the stimulus *x* input directly. During learning, *x* representation neurons arise also in M2 and M3, though the overall fraction of these neurons decreases from M1 to M3 (Fig. 6d). PE neurons are initially very rare, and emerge in all modules during learning, with the largest fraction concentrated in M2 (Fig. 6e). These results are consistent with activity of layer-specific primary sensory cortex neurons [12, 20, 58].

We finally evaluated the network responses for two stimulus-pairs. Similar to the single-module network model, mixed representation neurons arise in all modules after learning (Fig. 6f, g). The fraction of mixed representation neurons is maximal in M2, and it increases with the number of learned stimulus-pairs (Fig. 6f,g). We found that the more pronounced desegregation of neural representations is accompanied with a significant decrease in the median balance level in that module (Fig. 6h), suggesting that loose balance is the underlying circuit mechanism supporting the mixed predictive responses at the cellular level. Unlike our findings in M2, the fraction of mixed representation neurons and the median balance level in M1 and M3 do not show strong dependence on the stimulus dimensionality. These results highlight the impact of anatomical structure on shaping network function. Specifically, we found that different modules have different fractions of representation and prediction error neurons, reminiscent of recent experimental findings [18]. However, despite this heterogeneity, representations of stimuli and prediction error are desegregated in all modules after learning.

## Discussion

We investigated the neural representations formed in a class of recurrent neural networks that learn to generate high-dimensional predictions in natural conditions. Our mathematical analysis reveals key neural mechanisms supporting high-dimensional predictive coding; generates novel testable hypotheses for functional properties of the corresponding neural circuits; and provides a framework within which experimental data of large-scale neural recordings can be quantitatively analyzed. Additionally, our framework allows incorporating information on cell-types and anatomical structure into the model, which can elucidate their role in predictive computations.

We focused on a *recurrent* network model (Fig. 1) for two reasons. First, cortical circuitry that performs predictive processing is known to be highly recurrent. Plasticity of recurrent connections forms functional neuronal assemblies [64], which were suggested to underlie behaviorally-relevant sensory discrimination [65]. Second, predictions for sensory stimuli typically unfold over time, which can be naturally implemented by intrinsic dynamics of recurrent networks [32, 66]. While we focused on steady-state neural responses for mathematical tractability, our model could be extended in the future to study the temporal properties of high-dimensional predictive coding. Other interesting directions to extend our study are: networks with asymmetric connectivity, which could be done by imposing sparse connectivity [67]; and networks that learn predictions online [68, 69].

Our model suggests that balance between feedforward and recurrent input, or indeed between excitation and inhibition, can lead to robust internal predictions within local circuits. While this has been suggested previously [32–34, 70, 71], an important novel prediction revealed by our analysis is that in realistic conditions there is an optimal, finite balance level, which decreases with stimulus dimension (Fig. 2). Our theory further suggests that a network with infinitely high balance [33] could be especially vulnerable to noise in high-dimensional scenarios.

Based on our results, we hypothesize that the large degree of heterogeneity of empirical E/I balance levels in different experiments [51] may be a signature of the differences in the stimulus ensembles animals were exposed to. Our results in Fig. 2 and Fig. 3 suggest that this hypothesis could be tested systematically by exposing animals to increasingly rich sensory environments. Here too the temporal dynamics of the network may be important, as synaptic delays may affect the optimal degree of balance in circuits performing low-dimensional predictions [34, 72].

The role that balance plays in computing predictions has important implications for the source of predictive signals and the timescale of learning them. (*i*) Previous work has shown that cross-modal predictions are often *stimulus-specific* [12, 16, 48]: signals from one brain region can suppress responses to a particular predictable stimulus in another region (e.g., motor cortex activity suppressing visual cortical responses). It is notable that within our model those computations are performed without fine-tuning long-range projections [2]. Rather, local recurrent connections in the ‘receiving region’ can extract the predictions from long-range inputs with ‘promiscuous’ connectivity [11], relying on E/I balance and activity-dependent synaptic plasticity. (*ii*) Prediction-error responses in the same cortical region can arise at very different timescales, from as little as minutes [26] to days of training [12, 16]. We believe that the diversity of the identified E/I balance mechanisms (e.g., firing-rate adaptation, synaptic-scaling, Hebbian plasticity; see review in Ref. [43]), may explain this wide temporal range of predictive processing learning dynamics. Future work may reveal that our model has explanatory power also for the emergence of predictions over faster timescales than the experiments considered here and thus could be applied to predictive processing circuits in subcortical regions and in invertebrates.

An important finding of our work is that predictive representations are *desegregated*: neurons that signal prediction-errors for one stimulus-pair may faithfully represent the presence of stimulus for a second pair. Based on experiments where animals were probed with multiple types of unexpected sounds, we found signatures of this desegregation at the cellular-level in mouse auditory cortex (Fig. 4). Another recent study in mice performing multiple stereotyped motor actions reported mixed representations of the motor variables and reward prediction-errors across the neocortex [73], as suggested by our model for high-dimensional scenarios. Our model differs from previous work (e.g., [17,39,49,63]) by not explicitly assuming that separate neural populations encode prediction and prediction errors. Rather, the network develops mixed neural representations as a direct consequence of minimizing the multimodal prediction errors under energy constraints.

Our findings are related to the expanding literature on *mixed-selectivity* [74–76], where neurons exhibit complex tuning to multiple stimulus features. While even a random network can exhibit mixed-selectivity [75], the neurons’ tuning curves there are unstructured, which requires finely-tuned decoders to readout task-relevant variables. Here we report neurons that have mixed-selectivity to internally generated predictions of sensory and motor variables (Figs. 4, 5, 6). Crucially, the learned neural representations in our model are highly structured, and enable the reading out different stimulus features without ‘re-learning’ the decoder (Fig. 2).

Although neurons in our model network and in electrophysiological recordings from auditory cortex have mixed selectivity for stimuli and prediction-errors, the auditory cortex also contains neurons that more specifically encode prediction-errors [13]. Notably, the abundance of neurons with pure or mixed selectivity to stimulus and error could be also layer-specific [12]. This is recapitulated by our hierarchical network model (Fig. 6). Recent work in the mouse visual cortex identified specific genetic markers that are over-expressed in neurons that encode positive versus negative prediction-errors [18]. The differences in methodologies and time courses of analysis make direct comparisons across these studies difficult, and it remains possible that sensory cortex contains a large population of neurons that have shared roles in encoding stimuli and prediction-errors, as well as neurons that more strictly encode one or the other. Indeed, our analysis reveals that those classes of neurons may exist in different modules within a single network.

In summary, predictive processing is a ubiquitous and fundamental computation supporting diverse behaviors across animal species. Here we take a first step towards bridging the gap between theory of predictive processing and circuit-level neural recordings in predictive processing paradigms. Our results reveal the functional roles of specific circuit motifs and mechanisms in performing multimodal high-dimensional predictive processing. In a broader context, our work will advance the understanding of how the brain constructs complex internal-models by shedding light on commonalities and differences between biological predictive coding circuits and artificial systems, particularly those trained using self-supervised algorithms [39, 77].

## Methods

### Recurrent network model

Our model network consists of *N* neurons whose firing-rates are described by the time-dependent vector ***r***(*t*) = (*r*_1_(*t*), …, *r*_*N*_ (*t*)). The network is driven by high-dimensional stimulus input, denoted ***x***(*t*) = (*x*^1^(*t*), …, *x*^*P*^ (*t*)) and ***y***(*t*) = (*y*^1^(*t*), …, *y*^*P*^ (*t*)). The vectors ***x*** and ***y*** correspond to stimuli from two modalities that are paired during training.

The dynamics of the recurrent network are given by

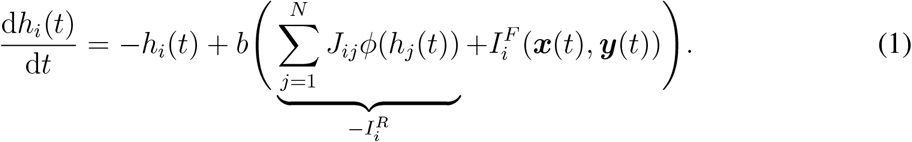

Here *h*_*i*_(*t*) is the voltage level of each neuron and is related to its firing-rate via a nonlinear activation function, *r*_*i*_(*t*) = *ϕ*(*h*_*i*_(*t*)). Note that the input each neuron receives in Eq. (1) is decomposed into the recurrent 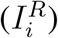 and feedforward 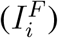 components. We rescaled the connectivity matrix *J*_*ij*_ and the feedforward input 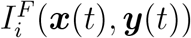 by a constant *b*, which can be interpreted as a gain parameter.

The explicit forms of *J*_*ij*_ and 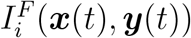 were determined based on a normative approach as follows (derivation details appear in SI §1). We assume that the neurons’ dynamics jointly minimize the following objective

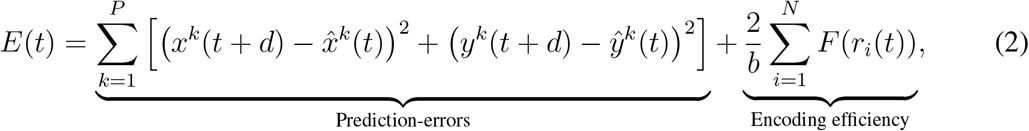

where 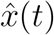 and 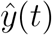 are the internal predictions generated by the network at time *t* and *F* (*r*) is a monotonically increasing function whose explicit form depends on *ϕ*, the nonlinear activation function (SI §1.1). For ReLU nonlinearity [*ϕ*(*z*) = max(*z* − *θ*, 0)], *F* (*r*) = (*r* + *θ*)^2^*/*2. Minimizing Eq. (2) is equivalent to performing Bayesian inference to extract the latent ‘cause’ of the sensory signals (SI §1.2). Furthermore, our network model generalizes previous models of predictive coding [1, 36, 38, 39, 63], by incorporating the effect of response nonlinearity into a regularization term that controls encoding efficiency. We note that the parameter *b* in Eq. (2) controls a trade-off between minimizing prediction-errors and maximizing encoding efficiency.

We further assume that the internal predictions are linear readouts of the network activity

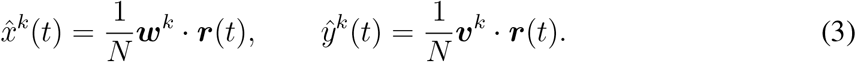

Here ***w***^*k*^, ***v***^*k*^ ∈ ℝ^*N*^ are the readout weight vectors. These internal predictions are, by definition, predictions of future input, as indicated by the delay *d* in Eq. (2). However, we will focus on the scenario where the input changes much more slowly than the neurons’ firing-rates. Therefore, on the timescale of firing-rate changes [Eq. (1)], we will regard the stimulus inputs to be approximately constant, i.e.,

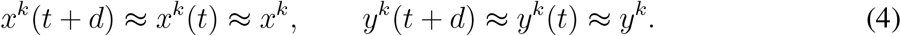

We assume that the weight vectors ***w***^*k*^ and ***v***^*k*^ change during learning so as to minimize the objective function *E*(*t*) [Eq. (2)]. This optimization process can be viewed as weight-changes governed by a combination of gradient descent on the squared prediction error in Eq. (2), and homeostatic plasticity (SI §1.1). If weights are initialized randomly, learning increases the correlation between the weight vectors (SI §1.1). Specifically, we show that in the large network size limit (*N* → ∞), the weight vectors have the following statistics,

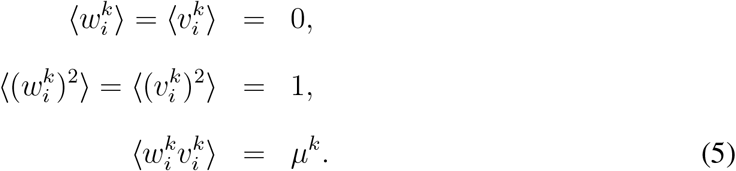

Here 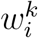 and 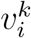 are the components of ***w***^*k*^ and ***v***^*k*^, which have zero mean and unit variance due to homeostatic plasticity. The correlation between them is *µ*^*k*^, which increases during learning (i.e., as the objective function *E* decreases). These weight changes can also arise from local plasticity rules applied to dendritic compartments (SI §1.3). For simplicity, unless noted otherwise, all stimulus-pairs have the same ‘age’, i.e., *µ*^*k*^ = *µ* does not depend on the index *k*. We further assume that the weight vectors have multivariate Gaussian distribution. Under these assumptions, we obtained analytical solutions for the dependence of steady-state firing-rate distribution on the stimulus input and the correlation *µ* in two limits (SI §2): the high-dimensional case where both *N* and *P* are large, and their ratio *α* = *P/N* is finite; and the low-dimensional case where only *N* is large, and *α* = 0.

The presence or absence of each stimulus was modeled by setting the corresponding components of ***x*** and ***y*** to 0 or 1. For example, the mismatch and match conditions for the *k*-th stimulus-pair correspond to,

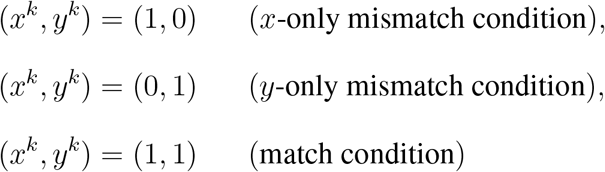

We extended our results to apply in scenarios with associations between more than two stimuli (SI §1.3).

### Geometry of representations of stimuli, predictions and prediction-errors

Under the above assumptions, the steady-state neural response vector [Eq. (1)] can be expressed as,

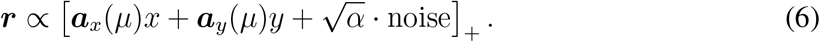

This form is revealing, since the stimulus-specific, *µ*-dependent vectors ***a***_*x*_(*µ*), ***a***_*y*_(*µ*) correspond to the directions along which the network encodes the stimuli in the *x*-only and *y*-only mismatch conditions. Eq. (6) also shows that, owing to the nonlinearity, the readout in the matched condition is not ***a***_*x*_(*µ*) + ***a***_*y*_(*µ*). The geometry of representing stimuli in the match and mismatch conditions is illustrated in Fig. 1d. Changes to these vectors during training (i.e., *µ* increases) correspond to the learned structure of neural representations of stimuli and prediction-errors. We further note that the magnitude of the noise in Eq. (6) depends on the stimulus dimensionality *α*, and thus it captures the interference between learned stimuli.

### Definition of balance level

The balance level for neuron *i* is defined as,

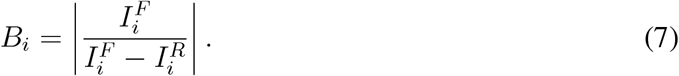

Here, 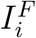 and 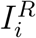 are the feedforward and recurrent input currents to neuron *i* at steady-state [Eq. (1)]. The balance level varies between neurons and between stimuli, because the weights 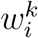 and 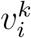 are different for different neurons and stimuli (indexed by *i* and *k*, respectively). The balance level distribution and its median shown in Fig. 2 were computed analytically (SI §2.3).

### Extracting the optimal balance level from experimental data

#### V-M experiment, Ref. [20]

We calculated the trial-averaged voltage of all the recorded L2/3 neurons as a function of time (Fig. 3a). Voltage level of each neuron was measured with respect to its baseline. We sampled 50 voltage levels from all recorded neurons and all time points in the match and mismatch time windows (Fig. 3a), which were −0.1 − 0s (match) and 0 − 0.1s (mismatch). The time *t* = 0 corresponds to point at which the treadmill was decoupled from visual flow in virtual reality. We then computed the standard deviation over those 50 samples of the voltage level in the match and mismatch conditions. By taking the ratio of these standard deviations, we obtained a dimensionless quantity that has a direct analog in the model: the standard deviation of *h*_*i*_ over neurons in the network in Eq. (1). Specifically, for *P* = 1, *θ* = 0, we computed this ratio explicitly (SI §2.1),

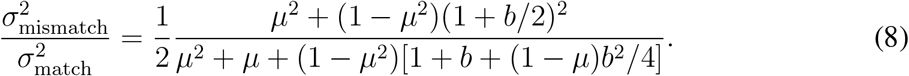

We use *µ* = 0.97 as the correlation value after training and fit this formula to the ratio obtained from data by adjusting the value of *b*. Using the best-fit value *b*^⋆^, we computed the median of balance level *B*^⋆^ in the network model (Fig. 3c).

#### A-M experiment, Ref. [12]

We calculated the trial-averaged firing-rates for all regular spiking neurons (*n* = 815) in the passive (mismatch) and movement (match) condition in two time windows: from *t* = −0.1s to stimulus onset (*t* = 0), and from stimulus onset to *t* = 0.06s (Fig. 3b). For every neuron, we calculated the change in its firing-rate between the two time windows in both conditions. We sampled 400 firing-rate change values from 815 neurons with replacement, and calculated the average firing-rate change in the passive and movement conditions. We computed the equivalent quantity in the model, i.e., average of *ϕ*(*h*_*i*_) over neurons in the network [Eq. (1)] in the match and mismatch conditions. For ReLU activation function, the ratio is also given by Eq. (8) and can be fit to the ratio obtained from the data by adjusting the parameter *b*. Again we calculated the median of balance level *B*^⋆^ based on the best-fit value of *b*^⋆^. The fitting procedure for both experiments was repeated 100 times, giving the scatter plot of estimated *B*^⋆^ values (Fig. 3c).

### Definition of functional cell types

We denote the steady-state voltage of neuron *i* in the mismatch conditions as 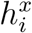 (*x*-only) and 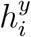 (*y*-only), and in the match condition as 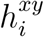. To classify neurons into functional types, deviations of individual neurons’ voltage response relative to the mean were compared to the standard deviation (denoted *σ*) of the steady-state voltage distribution. We evaluated *σ* using the voltage distribution in the *x*-only mismatch condition after learning (*µ* = 0.97).

A neuron *i* is a representation (*R*) neuron for the *x*-stimulus if it is depolarized upon presentation of the stimulus *x*, i.e., its voltage response in *x*-only mismatch condition is large, and its voltage responses in the match and mismatch conditions are similar. Mathematically,

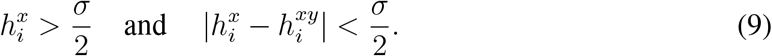

A similar criterion was used to identify *R* neurons for the *y*-stimulus. A neuron *i* is a prediction-error (*PE*) neuron if it signals the ‘mismatch’ between *x* and *y*, i.e., its voltage response in the *x*-only mismatch condition is large, and its voltage response in the match condition is small. Mathematically,

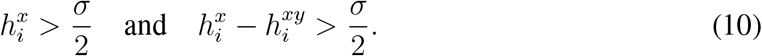

Neurons meeting these criteria are referred to as *positive* PE neurons, because their activity increases when *x* is presented but not expected (based on *y*). The activity of *negative* PE neurons increases when *x* is not presented but is expected. In our model, E neurons have a centered (zero mean) distribution of voltages for *α* = 0, therefore the threshold is applied to the voltage itself. For excitatory neurons in the high-dimensional regime (*α* > 0) and inhibitory neurons, since their voltage distribution has a non-zero mean, we used the centered voltage levels 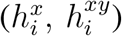 in the above criteria.

Note that neurons in the network may not belong to any of the those three classes (Fig. S3a). We computed the firing-rate statistics of neurons in the network analytically (SI §2, §3), which allowed use to obtain the fraction of *R* and *PE* neurons for different values of *µ* and *α*, shown in Fig. 4b,d. We further explored the effects of threshold level on the fraction of different functional types in Fig. S3b.

### Estimating functional segregation from responses to multiple stimuli from experimental data

We calculated the trial-averaged firing-rate change of each neuron in the match (active) and mismatch (passive) conditions, separately for each sound stimulus from our experimental data [13]. To calculate the segregation index for each type of probe sound, we restrict the analysis to neurons responsive in the passive condition to that probe sound and the learned (expected) sound. Responsive neurons were defined as those having firing-rate that was one half of the standard deviation above the mean firing-rate for the expected sound in the passive condition. Changing the threshold does not affect the results in Fig. 4e,f. For these neurons, we computed pairs of Δ values, defined as the difference between mismatch and match responses, for the probe and expected stimulus. The Pearson correlation coefficient between those Δ values was defined as the segregation index.

To estimate the similarity of the expected and probe stimuli, we computed individual neurons’ trial-averaged firing-rate change following presentation of those stimuli in the passive condition from our experimental data [13] (the same time windows used in the A-M experiment, Fig. 3). For each animal, we considered population firing-rate vectors consisting of all its recorded neurons. Representation similarity was defined as the Pearson correlation of those vectors for pairs of auditory stimuli (expected and probe, Fig. 4f). We note that this similarity in the model is calculated from the activity of all neurons that are active in either the expected or probe stimuli in passive condition.

### E/I network model

In the network with separate E and I neurons, the time-dependent voltages of E and I neurons are given by the following set of differential equations,

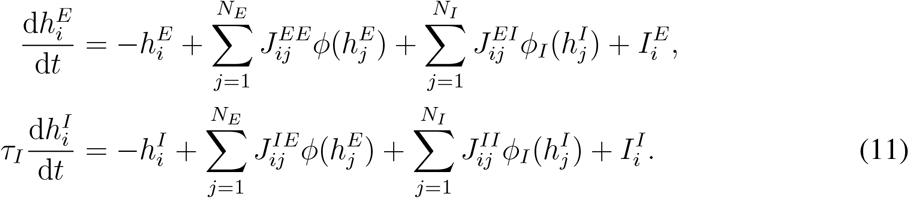

We assume that the activation function for inhibitory neurons is ReLU with zero threshold, *ϕ*_*I*_(*x*) = max{*x*, 0}. Matching the E neurons’ activity at steady state to the activity of neurons in our original network [Eq. (1)] gives constraints on the connectivity components and the feedforward input (SI §4),

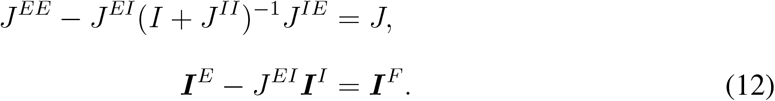

Here *J* and ***I***^*F*^ are the connectivity matrix and feedforward input used in Eq. (1). We further assume that the matrix *I* + *J*^*II*^ is invertible. In general, there are many possible solutions {*J*^*EE*^, *J*^*EI*^, *J*^*IE*^, *J*^*II*^, ***I***^*E*^, ***I***^*I*^} satisfying Eq. (12). We therefore identify a family of solutions. This continuum interpolates between the solution with private inhibition, where *J*^*IE*^ is equal to the identity matrix; and solutions with an inhibitory internal prediction, where rows of *J*^*IE*^ are given by the stimulus weight vectors (SI §4). Moreover, we show that up to a constant, the balance level defined earlier [Eq. (7)] is the same as the stimulus-specific, local component of the E/I balance level in the E/I network (SI §4).

We extended the definition of functional cell-types [Eqs. (9,10)] to I neurons. We note that here the average input to inhibitory neurons is not 0, so we subtracted the mean from the voltage level [*h*’s in Eqs. (9,10)] before applying the criteria on the deviations from the mean.

### Analyzing responses of regular spiking and fast spiking neurons

We estimated the connectivity structure parameter *λ*_*EI*_ based on recordings of regular spiking and fast spiking neurons [12]. Using the same time windows as Fig. 3b and Fig. 4e,f, we calculated individual neurons’ trial-averaged firing-rate change in the passive and movement conditions for the expected sound and the probe sound. Those firing-rate changes recorded in each animal form eight population vectors (regular/fast spiking, expected/probe sound, movement/passive). We calculated the Pearson correlation between population vectors under movement and passive conditions, giving four values for each animal, shown in Fig. 5d. The correlation values for presentation of the expected sound were regarded as ‘after learning’, while correlation values for presentation of the probe sound that was not associated with the lever press were regarded as ‘before learning’.

### Hierarchical recurrent network model

In the hierarchical network model, each neuron belongs to one of three modules, indicated by superscripts in the equations governing neural activity,

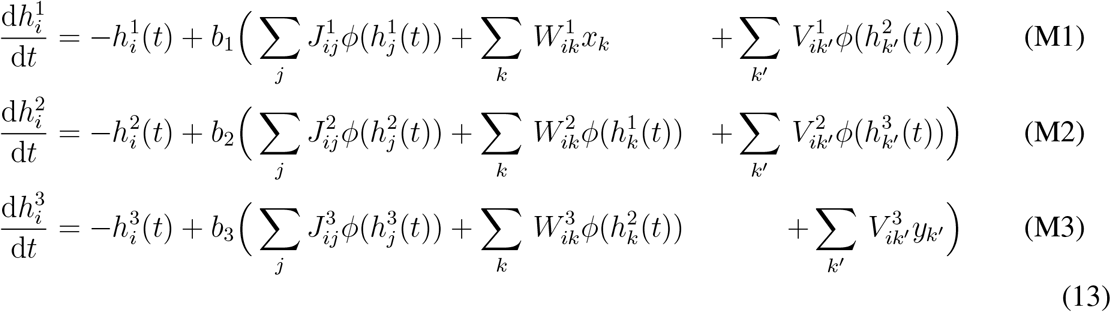

The definitions of feedforward and recurrent connectivity are generalizations of the single module network. Moreover, this model can be extended to a hierarchical network with a arbitrary number of layers. Details are provided in SI §1.3.

### Statistical tests

In Figs. 3c, 4f and 5d, we used two-sided, unpaired *t*-tests. ^⋆^ = *p* < 0.05 and ^⋆⋆⋆^ = *p* < 0.0005.

## Acknowledgments

The authors thank S. Azizpour Lindi, D. Bambah-Mukku, E. Mukamel, and I. Nelken for useful conversations, and I. Nelken for comments on a previous version of the manuscript. This work was supported by DARPA grant D21AP10162-00 (J.A.), DOE grant DE-SC0022042 (J.A.), NIH grants 1R01-NS135853 (J.A.), K99-DC020770 (N.J.A.), 1R01-DC018802 (D.M.S). B.W. thanks the UCSD Friends of the International Center for support. D.M.S. is a New York Stem Cell Foundation - Robertson Neuroscience Investigator. J.A. and B.W. thank the Kavli Institute for Theoretical Physics (KITP), Tel Aviv University, and ICERM at Brown University for hospitality during summer and fall 2023. KITP is supported by NSF grant PHY-1748958 and the Gordon and Betty Moore Foundation Grant No. 2919.02. ICERM is supported by NSF grant DMS-1929284.

## Author contributions

Developed the project and modeling approach, B.W., J.A. Solved, analyzed and simulated model, B.W. with inputs from J.A. Designed and performed experiments, N.J.A., D.M.S. Designed and performed data analysis, B.W. with inputs from J.A., N.J.A., D.M.S. Wrote paper, B.W., J.A. with inputs from N.J.A., D.M.S. Supervised the project, J.A.

**Fig. S1.**
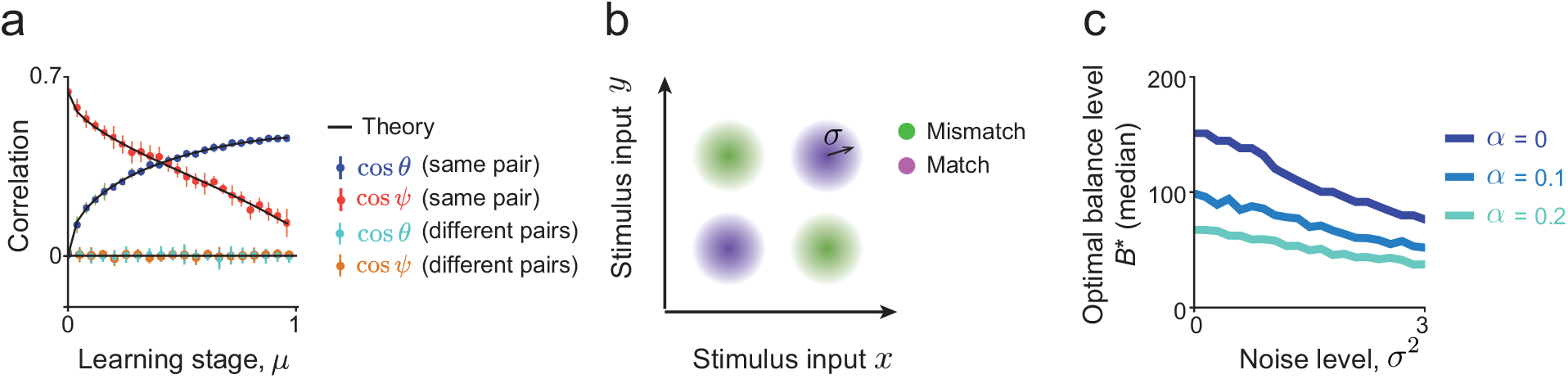
The geometry of predictive representations in the model. (a) Pearson correlation coefficient between neural responses in different stimulus conditions. As in Fig. 1, the angle *θ* is measured between the network’s responses to the two stimuli in mismatch conditions (i.e., ***r***_*x*_ and − ***r***_*y*_); while *ψ* is the angle between responses to the same stimulus in the match and mismatch conditions (i.e., ***r***_*xy*_ and ***r***_*x*_). Neural responses to stimuli from different stimulus-pairs remain uncorrelated, suggesting that the predictive signal learned by the network is stimulus-specific. Here *α* = 0. (b) Schematic of noisy stimulus inputs. Independent isotropic Gaussian noise (with S.D. denoted by *σ*) is added to the inputs in the match and mismatch conditions, relative to the noiseless stimulus presentation considered in Figs. 1,2. (c) The optimal balance level decreases as stimulus presentation becomes more noisy for all values of *α*.

**Fig. S2.**
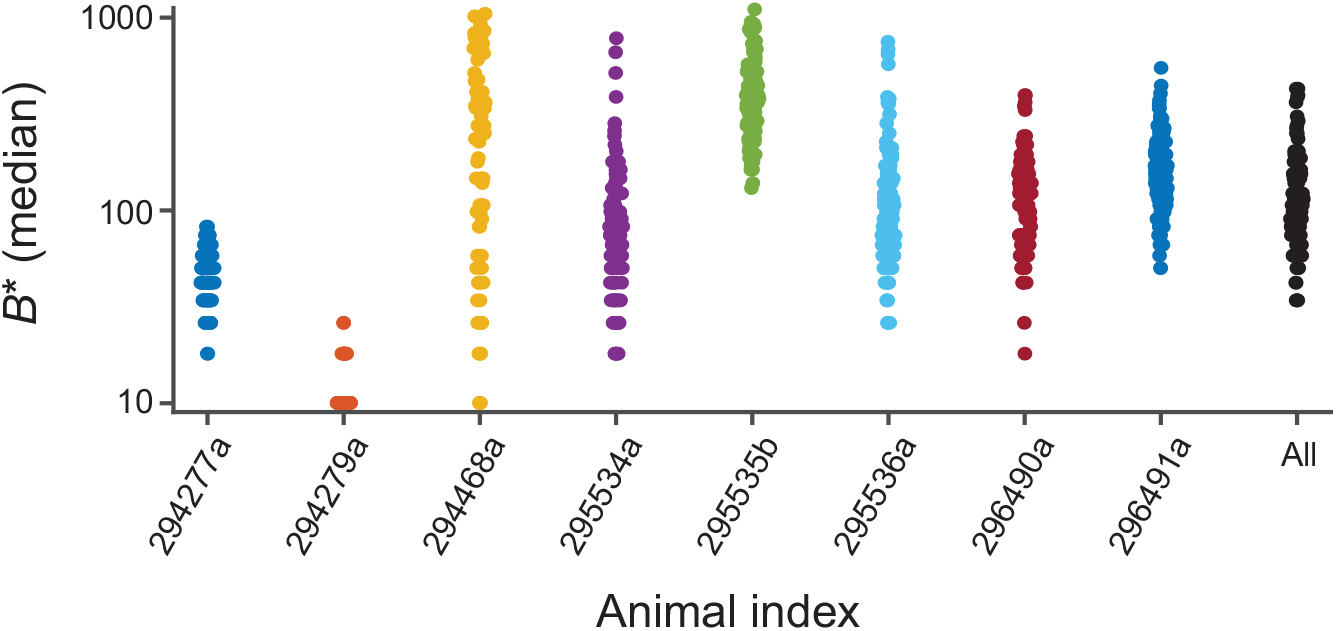
Estimated balance levels from individual animals. For each animal recorded in [12] (*n* = 8), the balance level was estimated as described in the Methods, sampling the firing-rates separately from each animal. There is marked variability across animals, suggesting that effects of learning multiple stimuli in the future are best studied *within animal* during learning.

**Fig. S3.**
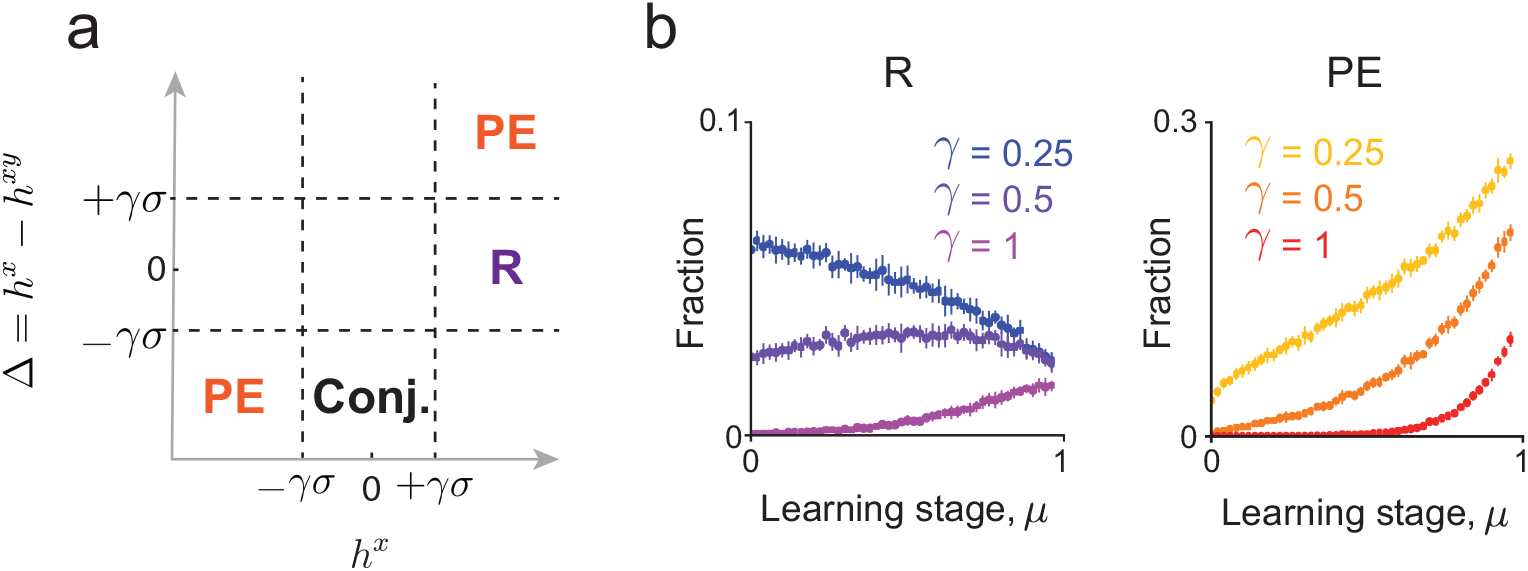
Abundance of functional cell types as a function of learning stage and classification threshold. (a) Criteria for classifying different functional cell types. The classification is based on setting two thresholds (*±γσ*) on the voltage response in the *x*-only mismatch condition 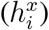, and its difference from the voltage response to match condition (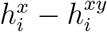, see Methods). The regions corresponding to prediction-error (*PE*) and representation (*R*) neurons for stimulus *x* are shown in the plot. Here we do not distinguish positive or negative *PE* neurons. Similar criteria are applied when replacing *x* with *y*. Also shown is the region corresponding to the conjunctive (Conj.) neurons, which have a small response in *x*-only mismatch condition but a large response in the match condition. (b) Fraction of *R* and *PE* neurons for different threshold values, as a function of the learning stage *µ*. The fraction of *PE* neurons increases during learning independently of the threshold. Here *α* = 0.

**Fig. S4.**
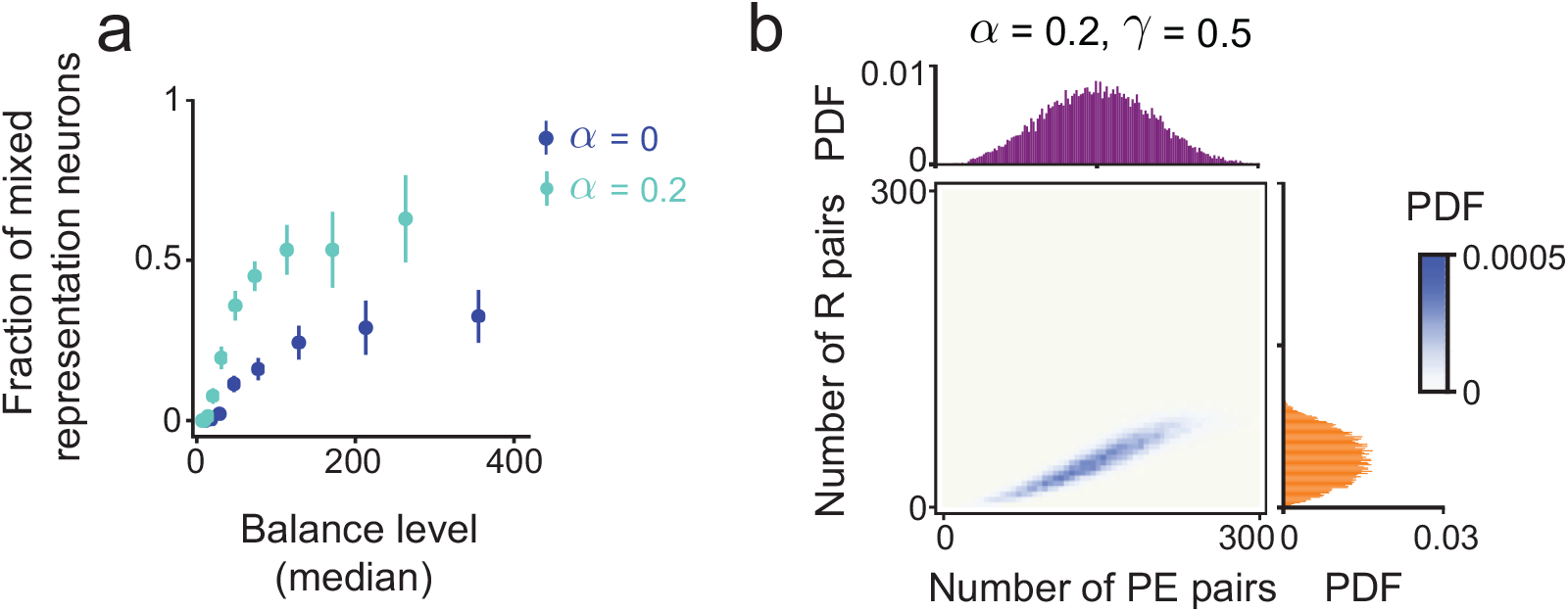
Fraction of mixed-representation neurons as a function of balance level and stimulus dimensionality *α*. (a) Here we vary the gain parameter *b* to generate a range of balance levels (median). As the stimulus dimensionality *α* increases, the fraction of mixed representation neurons for a fixed balance level also increases. (b) Each neuron in the network is a representation neuron for a certain number of stimulus-pairs (‘Number of R pairs’) and a prediction-error neuron for other stimulus-pairs (‘Number of PE pairs’). Plotted is the joint distribution of these two numbers for neurons in a network when it is trained to associate *P* = 400 stimulus-pairs. The corresponding marginal distributions are also shown. The joint distribution has a positive correlation. This indicates that when all *P* stimulus-pairs are considered, more neurons have a mixed representation than would be expected if the representation of stimulus and prediction-error was independent across pairs.

**Fig. S5.**
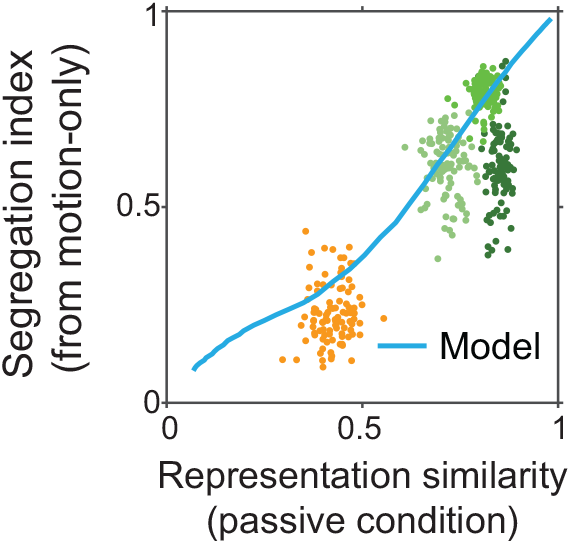
Segregation index as a function of representation similarity for different pairs of expected and probe sounds. Plotted are the segregation indices as a function of the representation similarity for different probe types (similar to Fig. 4f). Here the segregation indices are computed based on the differences Δ between the motion-only mismatch (passive: movement-only) and match (active: lever press + sound) neural responses. Colored points correspond to subsamples of the data. The results exhibit a similar trend as in Fig. 4f. The model curve shown in this plot is computed using a different sparsity level (by varying the firing threshold *θ*) compared to the values used in Fig. 4f. Under our main modeling assumptions: connectivity that is symmetric and puts the stimuli *x* and *y* on ‘equal footing’ during learning, synaptic weights with Gaussian statistics, and ReLU nonlinearity, we were not able to find a single value of *θ* to fit the data with two definitions of mismatch responses. Future work with more realistic network connectivity may give a choice of parameters that is consistent across both ways of comparing neural responses in expected and unexpected stimulus conditions.

**Fig. S6.**
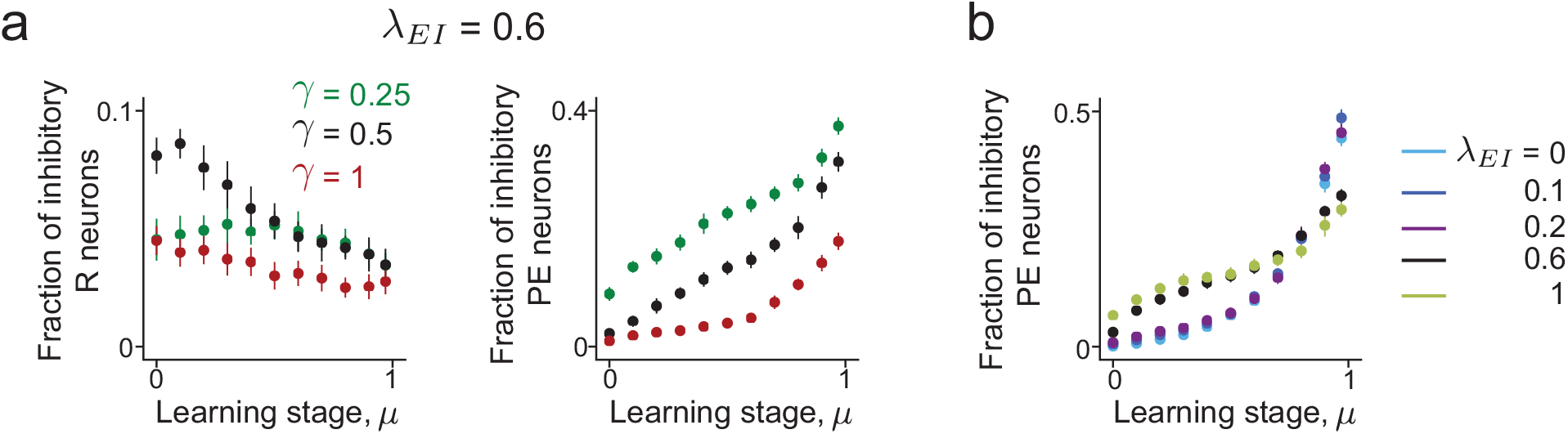
Abundance of functional cell types among inhibitory neurons. (a) Fraction of inhibitory representation (*R*) and prediction-error (*PE*) neurons at different learning stages (different values of *µ*) when using different voltage thresholds (±*γσ*). For the connectivity parameter that best matches our data (*λ*_*EI*_ = 0.6), the effect of learning is consistent across different thresholds. (b) Fraction of inhibitory prediction-error neurons at different learning stages for different values of *λ*. Unlike other network properties that do depend on the architecture of inhibitory connectivity (shown in Fig. 5), this quantity depends weakly on the parameter *λ*_*EI*_. In this plot we set *α* = 0.

**Fig. S7.**
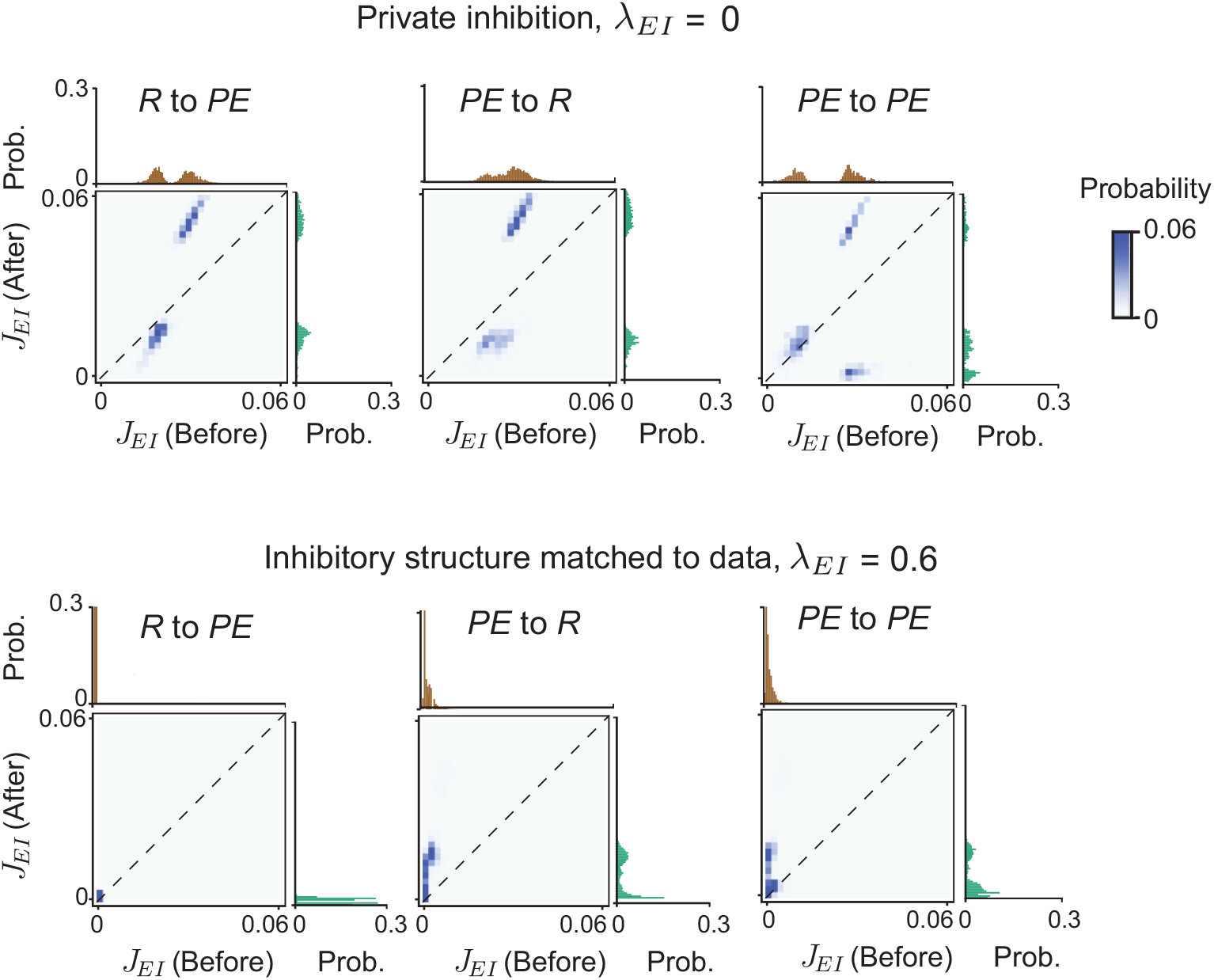
Changes to inhibitory to excitatory connections during learning do not depend strongly on the functional cell type of the target. Synaptic weight distribution of I-to-E connections before and after learning, when *λ*_*EI*_ = 0 (top) and *λ*_*EI*_ = 0.6 (bottom), for pairs of E and I neurons belonging to different functional classes: (*R* to *PE*, left; *PE* to *R*, middle; *PE* to *PE*, right). These fine-scale distribution show similar trends as in Fig. 5f,g.

**Fig. S8.**
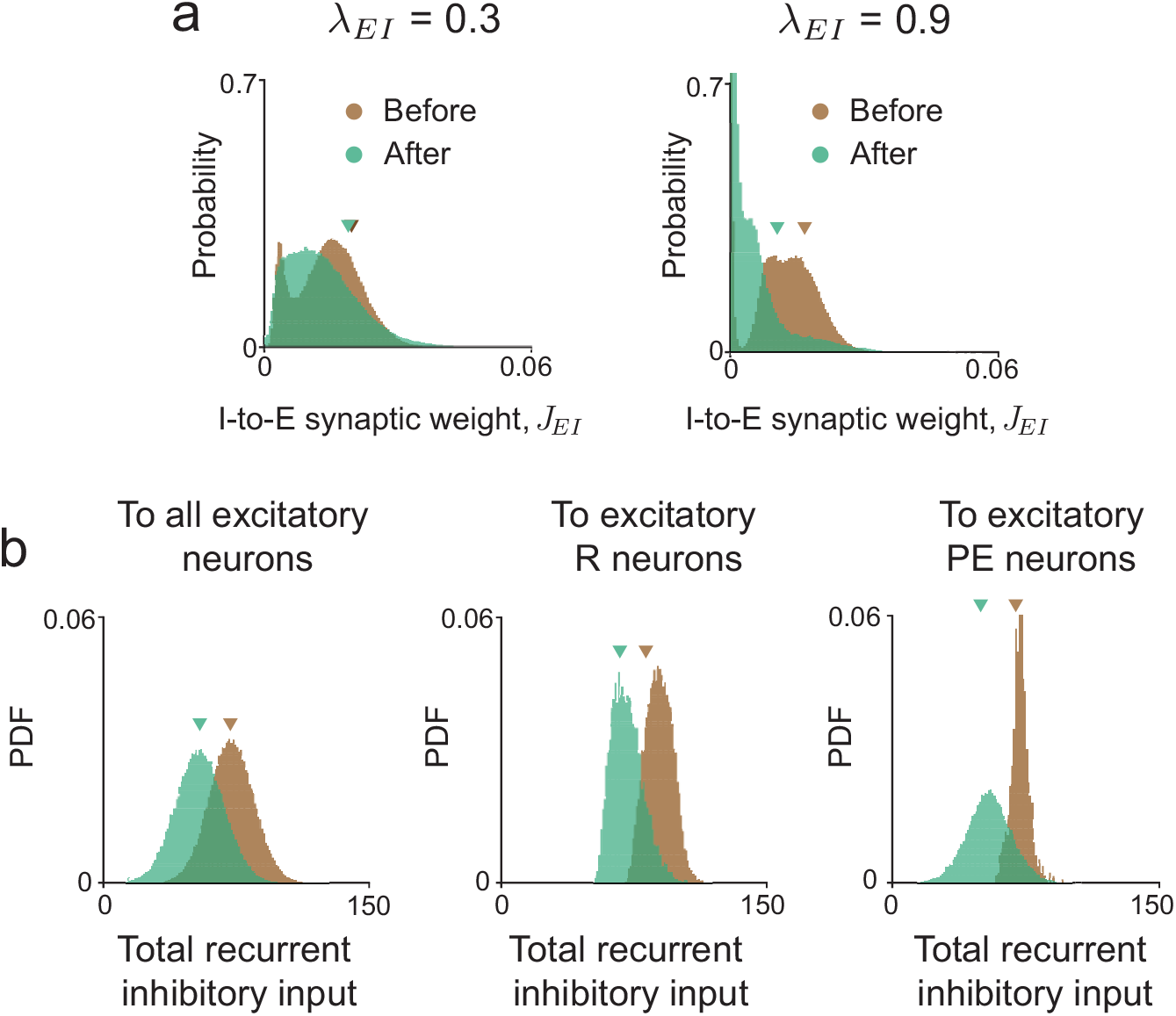
Learning predictive representations does not rely on overall potentiation of inhibitory connections, across different network architectures. (a) Synaptic weight distribution of all I-to-E connections before and after learning for values of *λ*_*EI*_ not shown in Fig. 5. There is no overall increase in the strength of inhibitory synapses after learning, suggesting that across different network architectures, predictive computations that lead to suppressed responses to expected stimuli are distributed. (b) Distribution of the total recurrent inhibitory input received by different populations of excitatory neurons, in the match condition. The overall inhibition received by excitatory neurons in the network decreases after learning.

## Supplementary Information

### 1. A NORMATIVE FRAMEWORK FOR HIGH-DIMENSIONAL PREDICTIVE PROCESSING

#### 1.1 The recurrent network model

We consider a network of *N* recurrently connected neurons, where the firing-rates of the neurons are denoted by the vector ***r***(*t*) = (*r*_1_(*t*), …, *r*_*N*_ (*t*)). The firing-rate of each neuron is related to its voltage level *h*_*i*_ via a nonlinear activation function, *r*_*i*_(*t*) = *ϕ*(*h*_*i*_(*t*)). We denote the learned paired inputs to the network as ***x***(*t*) = (*x*^1^(*t*), …, *x*^*P*^ (*t*)) and 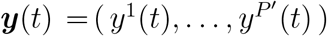. Notice that the dimensions of the paired inputs are not necessarily the same in this section.

In the predictive coding framework, the network continuously generates an internal prediction of the inputs. We assume that internal predictions (denoted 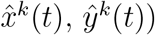 are linear read-outs from the network activity, i.e.,

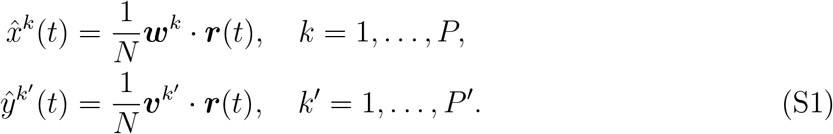

Here ***w***^*k*^, 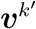 are the *N* -dimensional readout weight vectors.

Our aim is to derive a network model where the prediction-errors are minimized subject to some regularization term on encoding efficiency. Mathematically, we define the following objective function,

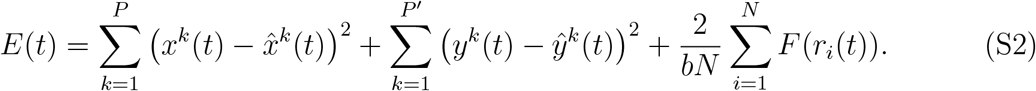

The first two terms of *E*(*t*) correspond to the prediction-errors. The regularization term, and the function *F* (*z*) in particular, depend on the nonlinear activation function *ϕ*. We consider those nonlinear activation functions where the firing-rate is *ϕ*_+_(*h* − *θ*) above a threshold *θ*, and 0 below the threshold. Mathematically,

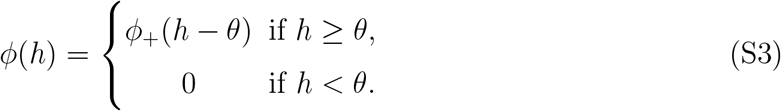

Here *ϕ*_+_ is a monotonically increasing smooth function which vanishes at 0, such that *ϕ* is continuous. This class of functions includes a number of activation functions used in previous work, e.g., rectified linear activation (ReLU, *ϕ*_+_(*h*) = *h*) and rectified *nonlinear* units, *ϕ*_+_(*h*) = *h*^*p*^ (*p* > 0), that coincide with ReLU for *p* = 1.

For this choice of *ϕ*, we show below that the recurrent network dynamics

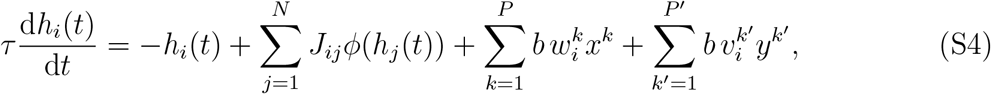

with the connectivity matrix and choice of regularization,

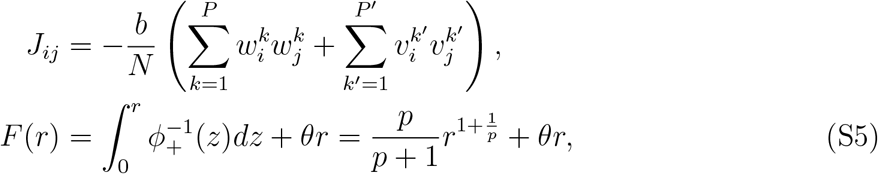

minimizes the objective [Eq. (S2)]. Note that adding a nonzero firing threshold (*θ* > 0) in the regularization function enforces sparse neural responses, penalizing large firing-rates. For the ReLU nonlinearity (*p* = 1), we have *F* (*r*) = *r*^2^*/*2 + *θr* = (*r* + *θ*)^2^*/*2 − *θ*^2^*/*2.

We assume that the timescale of changes to the inputs is much slower than the timescale of changes to neuronal activity, such that we can ignore potential time-dependencies of ***x*** and ***y***. Under this assumption, the objective [Eq. (S2)] can be written as a function of the neural activity and readout weights,

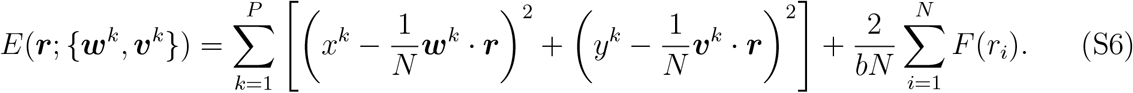

The neural activity ***r***(*t*) governed by the dynamical equations [Eq. (S4)] with the connectivity matrix [Eq. (S5)] minimizes the objective function [Eq. (S2)]. This can be shown by directly evaluating the time derivative of *E*(*t*):

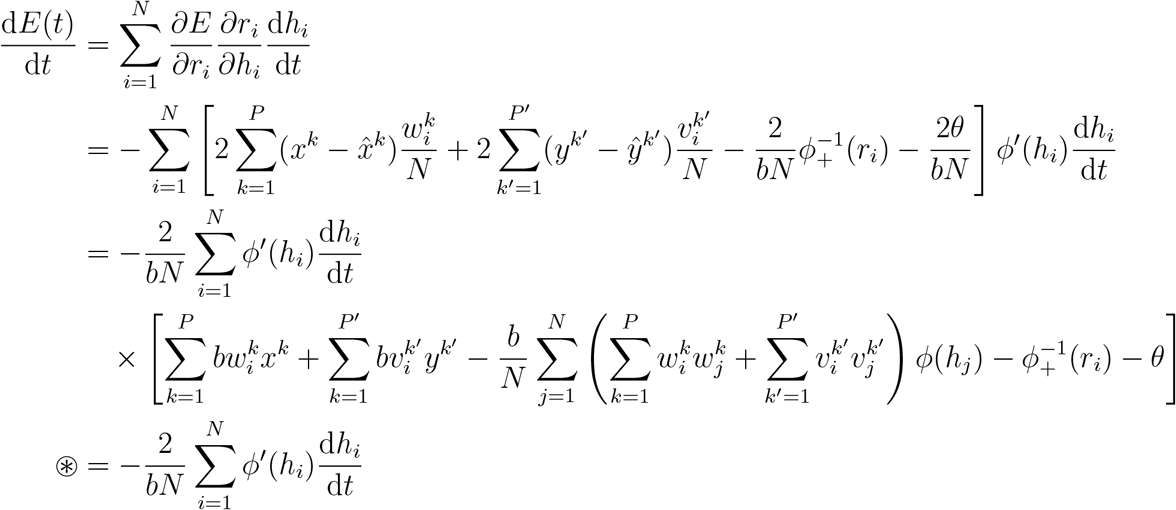

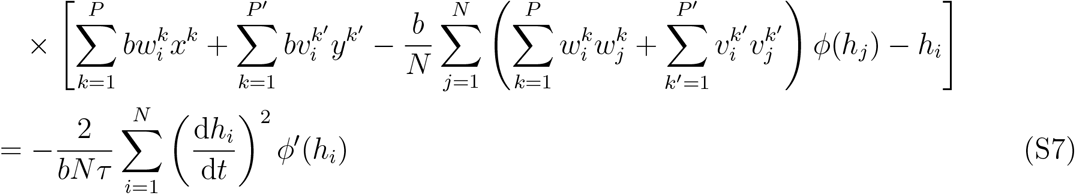

In the line indicated by ® we used the identity 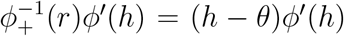. Each term in the sum that appears in the last line of Eq. (S7) is positive, so the time derivative of *E*(*t*) is negative. The existence of Lyapunov function for Eq. (S4) indicates that the network will reach a (stable) fixed point which satisfies for each neuron *i*,

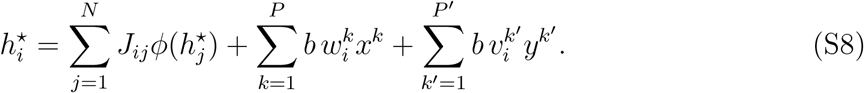

Moreover, since *E*(***r***) is a strictly convex function of the firing-rate vector ***r***, the optimal fixed-point solution ***r***^⋆^ is unique. From Eq. (S8), ***h***^⋆^ is also unique. Furthermore, that fixed point is a global minimum of *E*, which can be shown by evaluating the first-order derivatives of Eq. (S2) at the fixed point. Taken together, our results show that the network is guaranteed to reach a stable fixed-point for any input combination (indicated by *x*^*k*^ and *y*^*k*^), which is the minimum of Eq. (S2).

In the following sections, we will assume that there are *P* distinct pairs of stimuli indexed by *k*, (*x*^*k*^, *y*^*k*^). The corresponding feedforward weight vectors ***w***^*k*^, ***v***^*k*^ are assumed to be random, with mean 0. Associative training induces correlations between each component of the feedforward weights, via, for example, Hebbian-type plasticity. More precisely, for *i, j* = 1, …, *N* and *k, k*′ = 1, …, *P*,

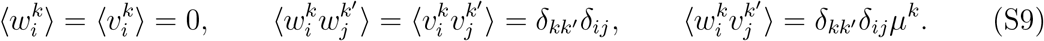

Here ⟨⋯⟩ denotes the expectation over the probability distribution of synaptic weights. To study how neural representations change during learning we vary *µ*^*k*^ systematically. Note that we have rescaled *µ*^*k*^ by *N* ^−1^ relative to the notation used in the main text.

Our choice of synaptic weight statistics [Eq. (S9)] arises from an optimization procedure that minimizes the objective function [Eq. (S2)]. Indeed, performing gradient descent on *E* within a short time window Δ*t* induces the following weight changes,

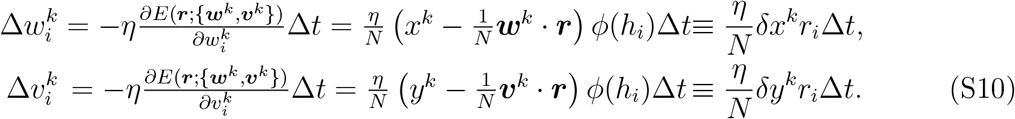

We assume that the learning rate is small *η* ≪ 1, such that the neural dynamics [Eq. (S4)] remain at the steady state ***r***^⋆^. We will show below (SI §2.1) that during associative learning (*x*^*k*^ = *y*^*k*^ = 1), the variables representing prediction errors are non-negative (*δx*^*k*^, *δy*^*k*^ ≥ 0), which implies that the weights could grow unbounded during learning.

To prevent this potential blow-up, we introduce a normalization mechanism that regularizes the weights. After each ‘learning-step’ [Eq. S10], the weights change according to a ‘homeostatic-step’,

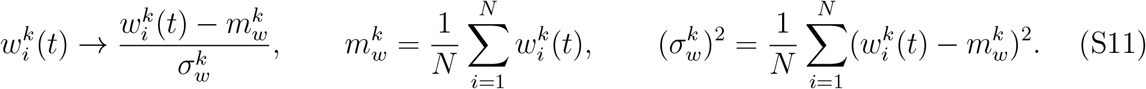

Here 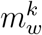 and 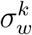 are the means and the standard deviations of the weight vector ***w*** computed over the *N* neurons. Similar updates are applied to the weights ***v***. We show that under these update rules, *µ*^*k*^(*t*), the correlation between ***w***^*k*^ and ***v***^*k*^ at time *t* during the learning process, increases monotonically.

We first note that the homeostatic step [Eq. (S11)] ensures that weight vectors have zero mean and unit variance. Upon presentation of the stimulus-pair *k*, the steady-state input to neuron *i* is independent of inputs to other neurons. Additionally, in the *N* → ∞ limit, 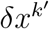 and 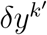 are nonzero only if *k*′ = *k*. These properties are shown explicitly using a replica calculation below (SI §2.1). It is therefore sufficient to verify that applying the learning-step [Eq. (S10)] does not lead to a decrease in the correlation. This can be done by a direct calculation of the correlation in Eq. (S11). Notice that in the *N* → ∞ limit,

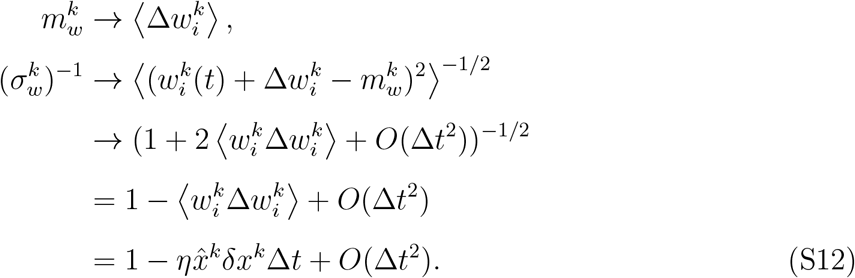

Therefore the weight 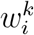 after the learning and homeostatic steps is,

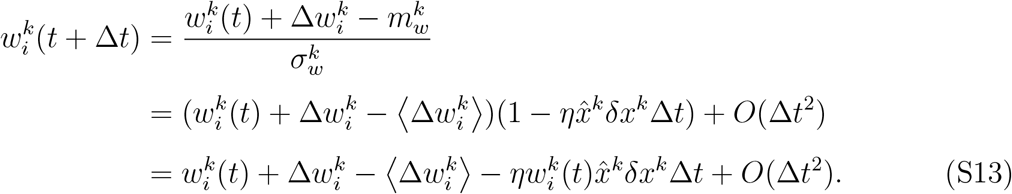

Using this approximation and a similar expression for 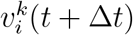, the correlation is now,

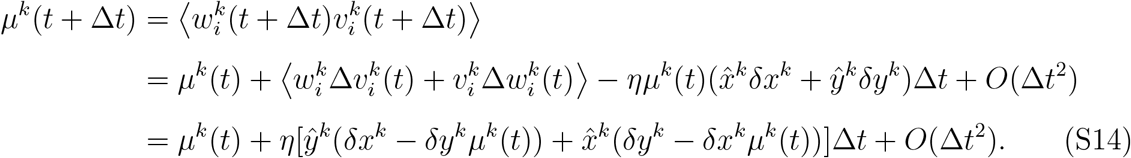

In the match condition (*x*^*k*^ = *y*^*k*^ = 1) we have from symmetry that 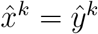 and *δx*^*k*^ = *δy*^*k*^. We will show in SI §2.1 using a replica calculation that 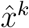, *δx*^*k*^ ≥ 0, which together imply that the bracket is positive when *µ*^*k*^(*t*) ≤ 1. Thus the correlation between the weight vectors increases during associative learning. This justifies our choice of weight statistics [Eq. (S9)] as a description for the network during associative learning.

#### 1.2 A Bayesian inference perspective of the network model

The predictive coding framework is often used to account for inference of latent causes of sensorimotor inputs to the brain, based on prediction and prediction-error signals [1, 17, 49, 55]. In this section we show that our model can similarly be viewed as a network performing Bayesian inference. Specifically, the network’s neural dynamics [Eq. (S4)] implement the inference (or state estimation) of latent variables driving inputs. Moreover, the slow synaptic weight changes during learning [Eq. (S9)] can be viewed as a mechanism for improving the accuracy of the inference performed by the network.

We consider a scenario where sensory inputs in the environment are generated by a probabilistic generative model, *p*(***x, y***|***r***), where ***x, y*** are the (possibly time-dependent) sensory inputs and ***r*** represents the latent variables that determine the statistics of the sensory inputs. We denote the prior distribution over the latent variables as *p*_0_(***r***). Then given the sensory inputs ***x, y***, the latent variables ***r*** can be inferred by maximizing the posterior distribution via Bayes’ rule,

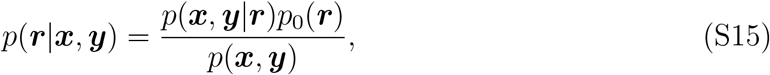

where *p*(***x, y***) = ∫*p*(***x, y***|***r***)*p*_0_(***r***)*d****r*** is the marginal distribution of the sensory inputs, independent of the latent variables.

Suppose that the generative distribution is a multivariate Gaussian and that its mean is a linear readout of the latent variables,

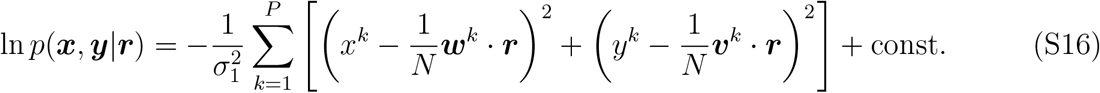

Further suppose that the prior distribution has the form,

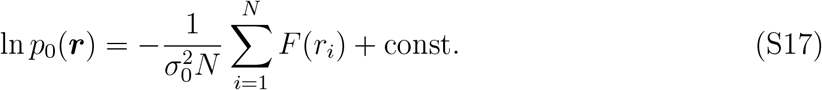

Then, recalling Eq. (S7), we see that the neural dynamics [Eq. (S4)] maximize the log posterior distribution,

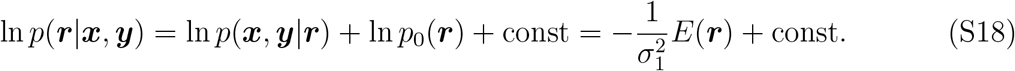

Here *E*(***r***) is the objective function in the previous section with 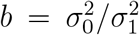. Thus, our model’s gain parameter *b* is related to the prediction accuracy *σ*_1_. The latent variables ***r*** here correspond to the firing rates of the neurons in the network.

In the more general case where sensory inputs are not generated exactly according to Eq. (S16), prediction accuracy can be improved by adjusting the readout weights ***w***^*k*^, ***v***^*k*^ to maximize the log posterior distribution [Eq. (S18)] based on the learning rule [Eq. (S11)]. This weight optimization procedure is equivalent to using a variational approach for maximizing the Bayesian model evidence, as introduced in previous predictive coding literature [17, 49, 78]. We also note that the nonlinear response function *ϕ* appears in the regularization *F* (***r***) [Eq. (S5)] is linked to the ‘encoding’ of prior information on the latent variables, *p*_0_(***r***).

#### 1.3. Extensions of the network model

##### 1.3.1. Associations between more than two modalities

Our network model can be generalized to apply to scenarios in which the animal is trained to associate multiple (*M* ≥ 3) sensorimotor inputs. Here the network generates internal predictions for each input, that can be linearly read-out,

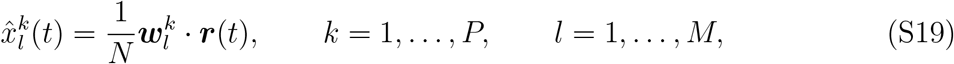

where 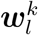 are the readout weights for each input in each stimulus modality. The objective function [Eq. (S1)] is now,

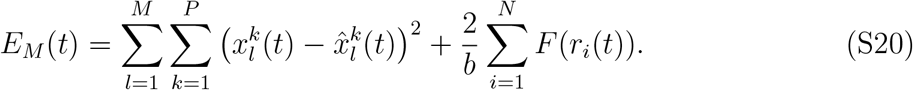

The network dynamics and recurrent connectivity matrix are,

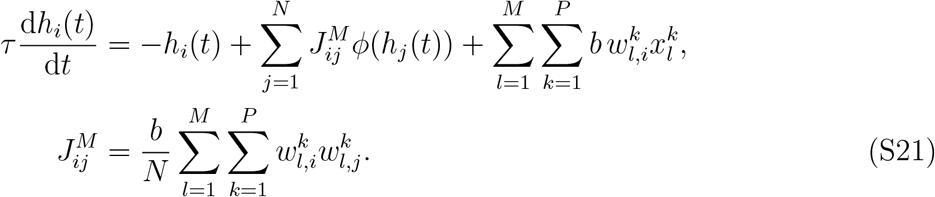

Using similar derivations as above, one can show that (*i*) *E*_*M*_ (*t*) is a Lyapunov function for the network dynamics, and (*ii*) the network will reach a unique stable fixed point for any combination of the inputs 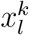. Assuming that the feedforward weights corresponding to associated stimuli become increasingly correlated during learning (similarly to the *M* = 2 case), will make this model useful for studying predictive representations when training animals on more complex stimulus combinations.

##### 1.3.2. Neurons with dendritic compartments

The network model with point neurons [Eq. (S4)] and the associated learning rules [Eq. (S11)] can be extended to a model with dendritic compartments. Crucially, this extension allows the learning rule to be realized by local plasticity rules.

Following the approach introduced in Refs. [37, 38], we first notice that Eqs. (S4-S5) can be rewritten by decomposing the connectivity to synaptic weights onto specific dendrites, giving,

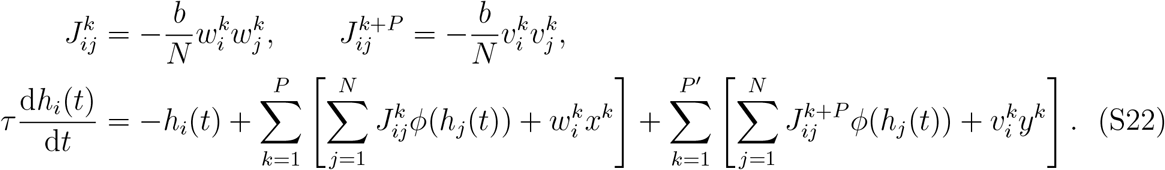

Here we think of *h*_*i*_(*t*) as the *somatic* membrane potential of neuron *i*. Next we introduce *P* + *P*′ dendritic compartments corresponding to neuron *i*. The voltages 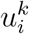 for *k* = 1, …, *P* and for *k* = *P* + 1, …, *P*′ + *P* are respectively governed by the equations,

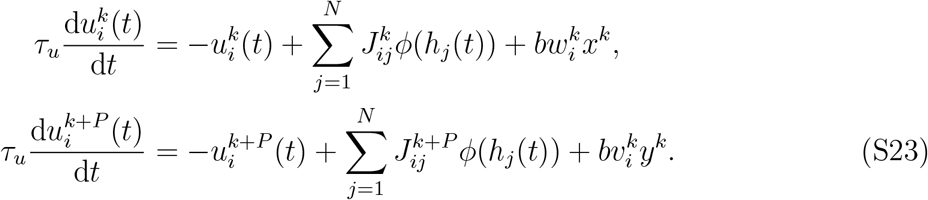

The somatic voltage level is then driven by the dendrites,

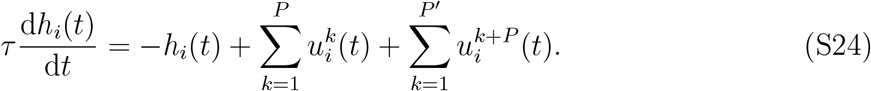

Under the assumption that dendrite voltage changes faster than somatic voltage, *τ*_*u*_ ≪ *τ*, this recovers our original model with point neurons [Eq. (S4)].

The learning rule of the dendrite-specific feedforward weights is given by,

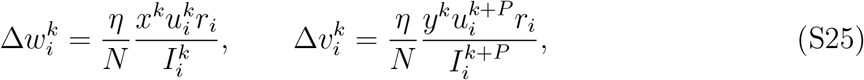

where we have denoted 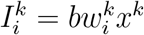 and 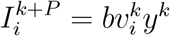. Note that the quantities on the right hand side are ‘local’ to the feedforward synapses 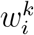 and 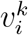. At steady-state, 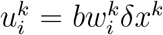 and 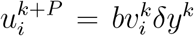. Together with the definition of 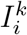, this learning rule is the same as Eq. (S10). To avoid unbounded growth of the weights in this setting, we assume a similar homeostatic mechanism which recovers the previous learning rule for the feedforward weights [Eq. (S11)].

The recurrent weights are subject to the learning rules,

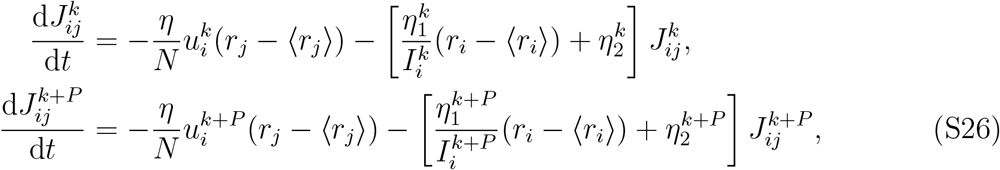

where 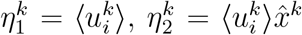 and 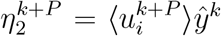 are activity-dependent learning rates. The dendrite-specific synaptic weights [Eq. (S22)] are solutions to these learning dynamics.

We note that the increase in correlation between ***w*** and ***v*** during learning is reflected in this plasticity rule by the dependence of both *J*^*k*^ and *J*^*k*+*P*^ on the firing rates ***r***. Since those rates depend on inputs from both modalities, both sets of dendrite-specific synaptic weights change based on the interplay between the multimodal input.

##### 1.3.3. Hierarchical network architecture

In the recurrent network model studied thus far, a single module integrates inputs from multiple sensorimotor modalities. Here we generalize this model to a network consisting of multiple (*L*) modules arranged in a layered structure. Each module has *N* neurons with firing rates denoted as ***r***^*l*^, *l* = 1, …, *L*. We assume that the paired stimulus inputs enter the network via the first and the last module respectively (Fig. 6a). For convenience, we denote the inputs as ***x*** ≡ ***r***^0^ and ***y*** ≡ ***r***^*L*+1^.

Each module generates predictions of the activity of ‘adjacent’ (earlier and later) modules, i.e., neurons in module *l* generate predictions for neural responses in modules *l* − 1 and *l* + 1. Those predictions are assumed to be linear readouts of the firing rates,

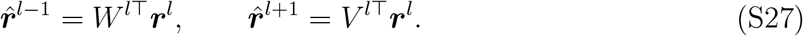

Here *W*^*l*^, *V* ^*l*^ are the readout matrices. The objective function for this hierarchical network is a sum of the objective function applied to each module-separately with the corresponding prediction errors and firing-rate regularization,

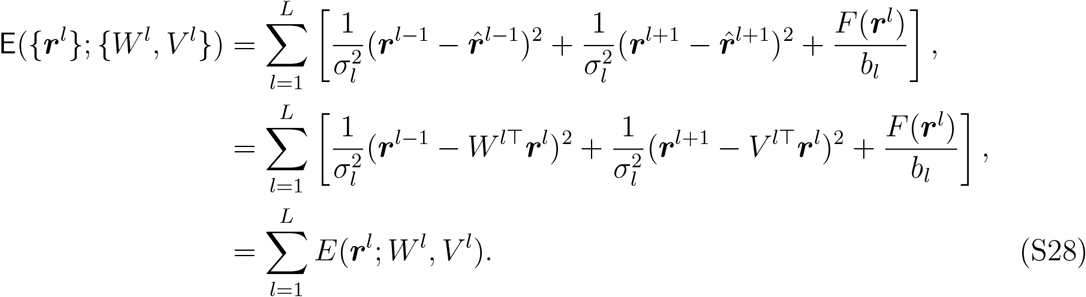

Here *σ*_*l*_ measures the module-specific precision of predictions and *b*_*l*_ is the module-specific regularization. The assumption that the neurons in module *l* minimize the module-specific loss *E*(***r***^*l*^; *W*^*l*^, *V* ^*l*^) implies that the neural dynamics within each module and the recurrent synaptic weights have identical form to those in the single-module case,

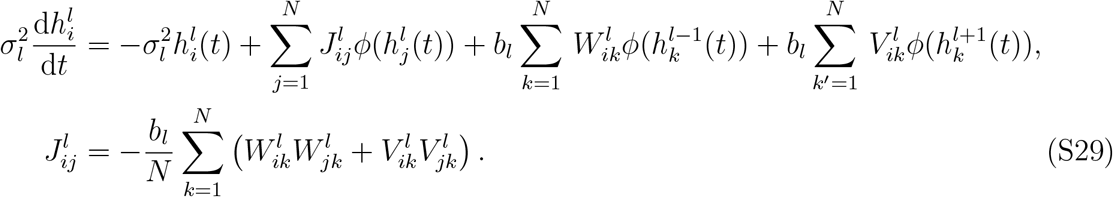

Similarly to the network with a single module, we assume that associative learning induces correlations between the corresponding weight vectors for each stimulus-pair. In the hierarchical network, the feedforward weight matrices in the first and last modules *W* ^1*T*^, *V* ^1^ have dimensions *N* × *P* rather than the *N* × *N* dimensions of matrices in intermediate modules. We assume that the intermediate feedforward weight matrices have rank *P* (the stimulus dimension). Furthermore, because the process of training the network to associate stimuli *x*^*k*^ with *y*^*k*^ is symmetric under the substitutions *x* ↔ *y, W* ↔ *V*, we assume that *W, V* are symmetric matrices. With these assumptions, we the weight matrices are,

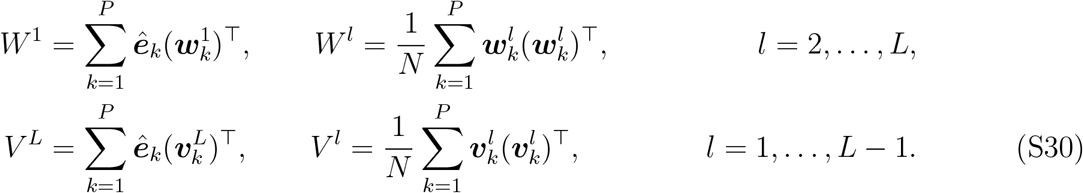

During associative learning, these weight vectors become correlated and their statistics are,

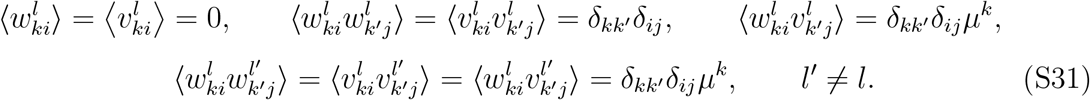

Here the first line specifies the weight statistics within module *l*, and the second line specifies the statistics across modules. The recurrent connectivity within each module simplifies to a form which is identical to that of the single module network,

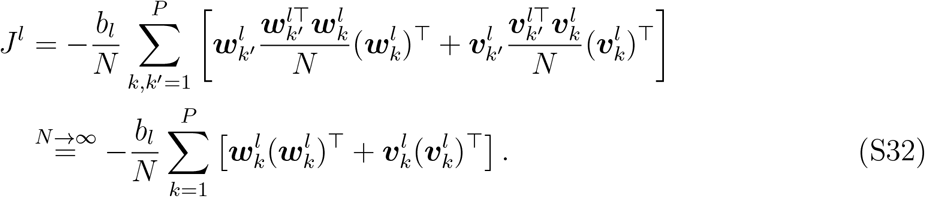

### 2. PREDICTIVE REPRESENTATIONS IN RECURRENT NETWORKS

When the stimulus inputs do not depend on time, the objective function *E* [Eq. (S2)] can be viewed as a function of the firing-rates and synaptic weights,

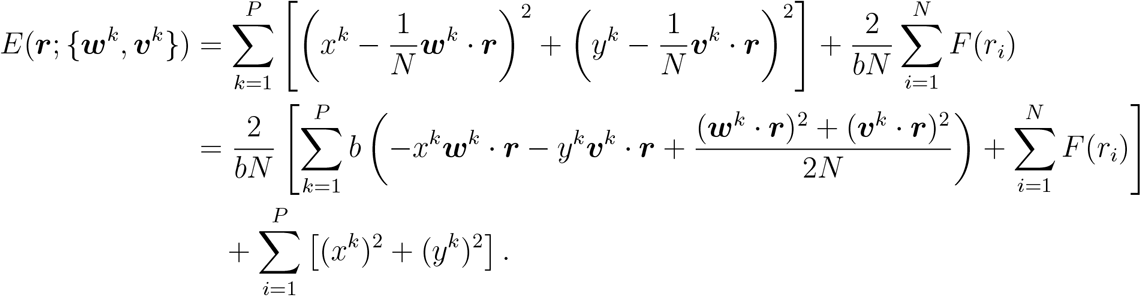

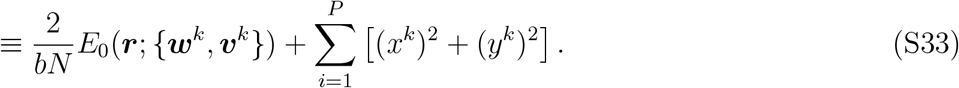

The steady state firing-rates can be expressed as minimization over *E*_0_, since the second term in Eq. (S33) does not depend on ***r***,

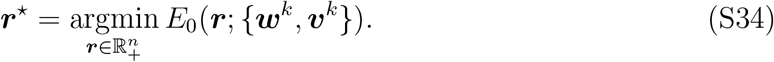

Next we will use the replica method [79, 80] to calculate the firing-rate distribution of neurons in the network,

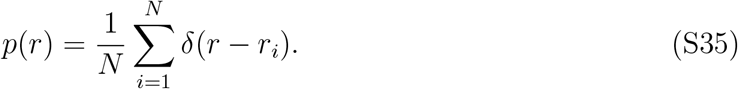

In general, firing-rates in the network depend on the specific realization of random weights ***w***^*k*^, 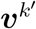. We find however that in the *N* → ∞ limit, the firing-rate distribution is self-averaging and depends only on the distribution of synaptic weights. By choosing which of the *x*^*k*^ and *y*^*k*^’s are nonzero, we can study the network response in different stimulus conditions. For convenience, we assume that at any given time, only a finite number of stimulus-pairs are presented, or equivalently, there are only *K* = *O*(1) pairs (*x*^*k*^, *y*^*k*^) for *k* = 1, …, *K*, where at least one stimulus is nonzero. We set the decay timescale to *τ* = 1.

#### 2.1. Replica calculation of the firing-rate statistics

We consider the partition function

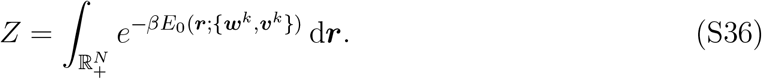

We suppress the domain of integration over firing-rates for readability in the following calculations. In the limit *β* → ∞, the dominant contribution to *Z* comes from the fixed point solution which minimizes *E*_0_(***r***; {***w***^*k*^, ***v***^*k*^}) in Eq. (S34). The logarithm of the partition function concentrates around its expectation, so we use the replica trick,

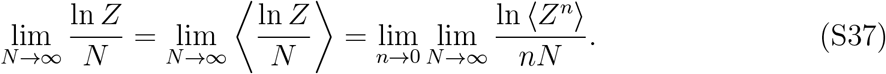

We make the standard assumption that the order of the limits can be exchanged in the last equality. We first calculate ⟨*Z*^*n*^⟩. For readability, we use *g* for the gain parameter (instead of *b*) in Subsection 2.1, and *a, b* = 1, …, *n* for the replica indices. Without loss of generality, we assume that the presented stimuli (i.e., indices *k* such that *x*_*k*_ or *y*_*k*_ is nonzero) are the first *K* pairs, *k* = 1, …, *K*.

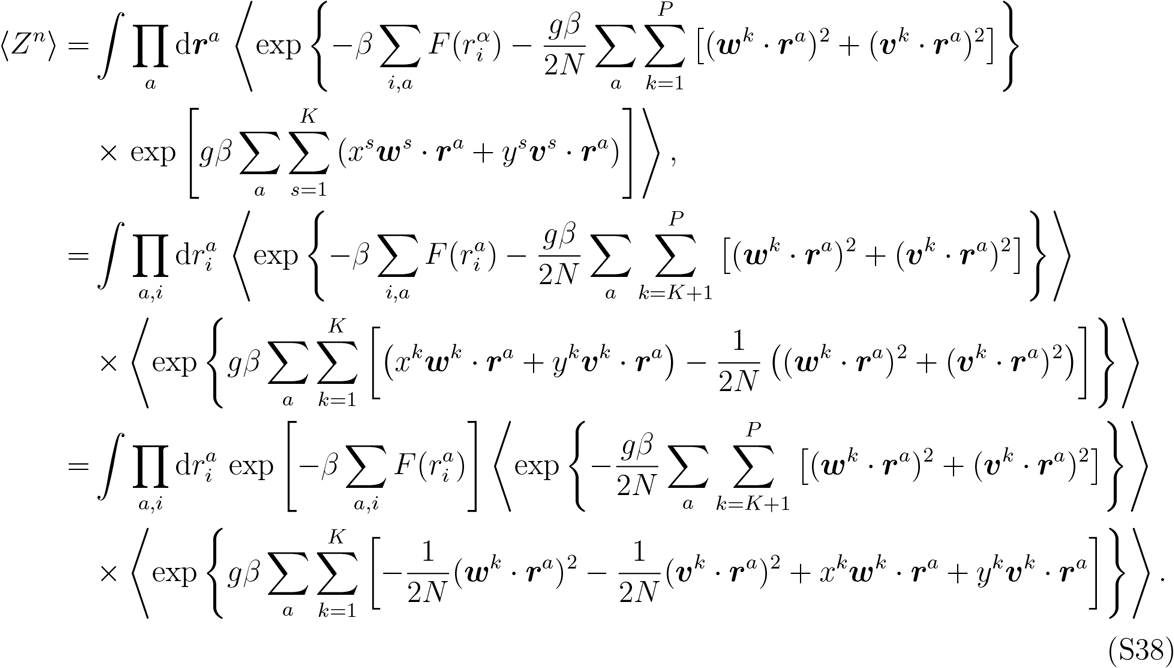

Notice that we have split the summation over all *P* stimulus-pairs and averaging over the corresponding synaptic weights into the presented pairs (*k* = 1, …, *K*) and the rest (*k* = *K* + 1, …, *P*). We first perform calculations for the *P* − *K* ‘absent’ stimulus-pairs. Using the integral representation of Gaussian function, we get,

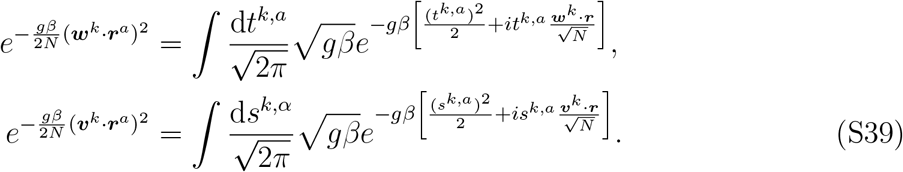

Using these, the term corresponding to the *P* − *K* absent stimulus-pairs becomes,

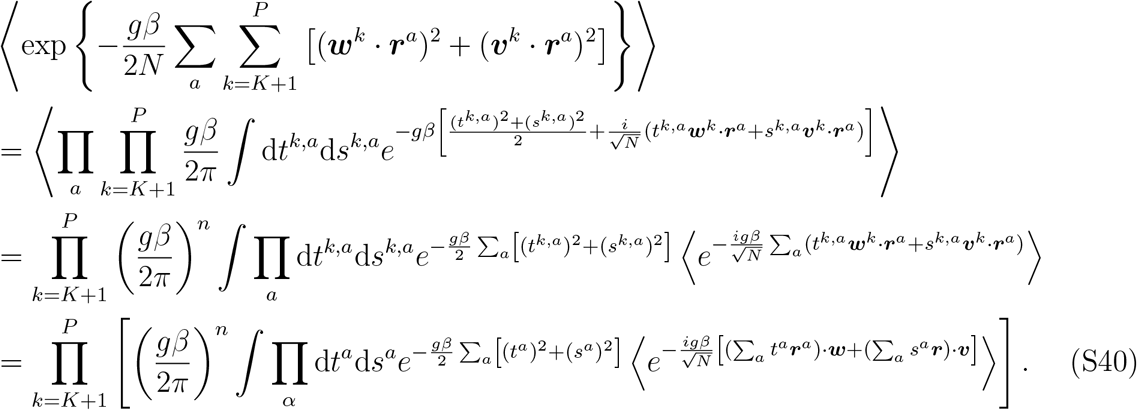

In the last line we have suppressed the superscript *k*. Recall that for each *k*, angle brackets denote the average over a pair of synaptic weight vectors, each of which has components sampled from the same distribution with mean 0 and correlation *µ*^*k*^ [Eq. (S9)]. We work out the last factor of the integrand,

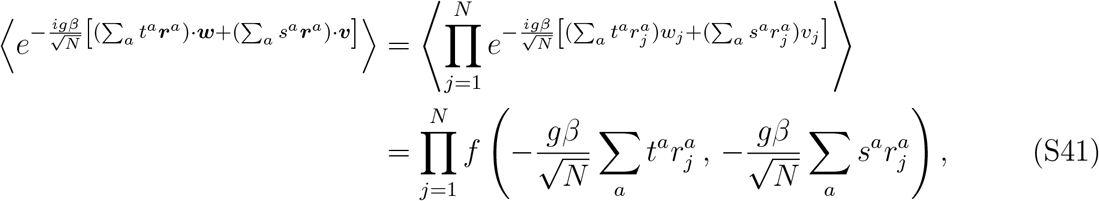

where *f* (*x, y*) is the joint characteristic function of the random vectors ***w***^*k*^, ***v***^*k*^ with correlation *µ*^*k*^. The Taylor expansion of *f* (*·, ·*) in the limit *N* → ∞ is,

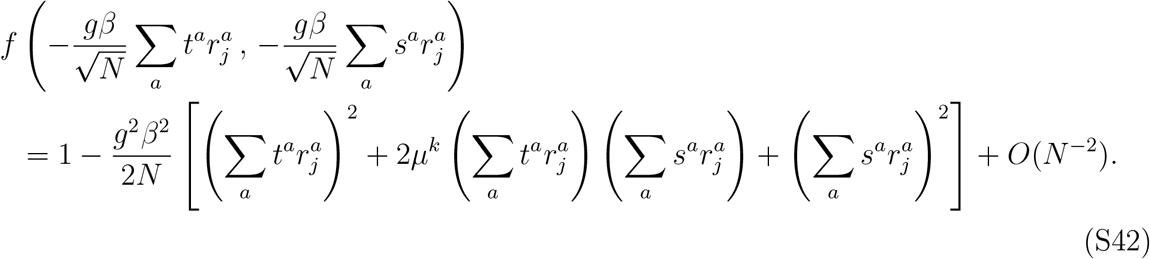

Using this we get,

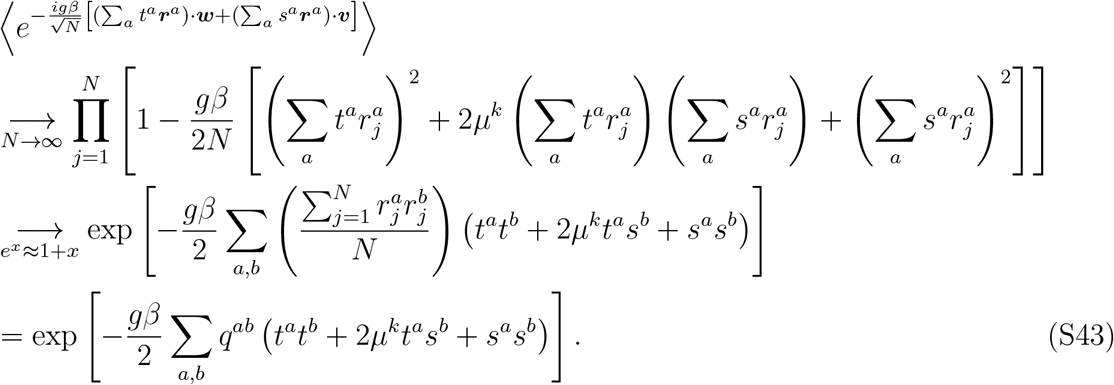

In the last line we have introduced the usual definition of the order parameter,

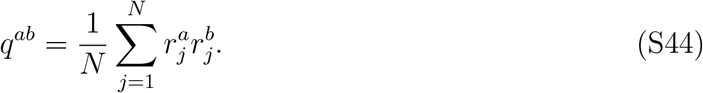

Collecting terms, we find that Eq. (S40) becomes,

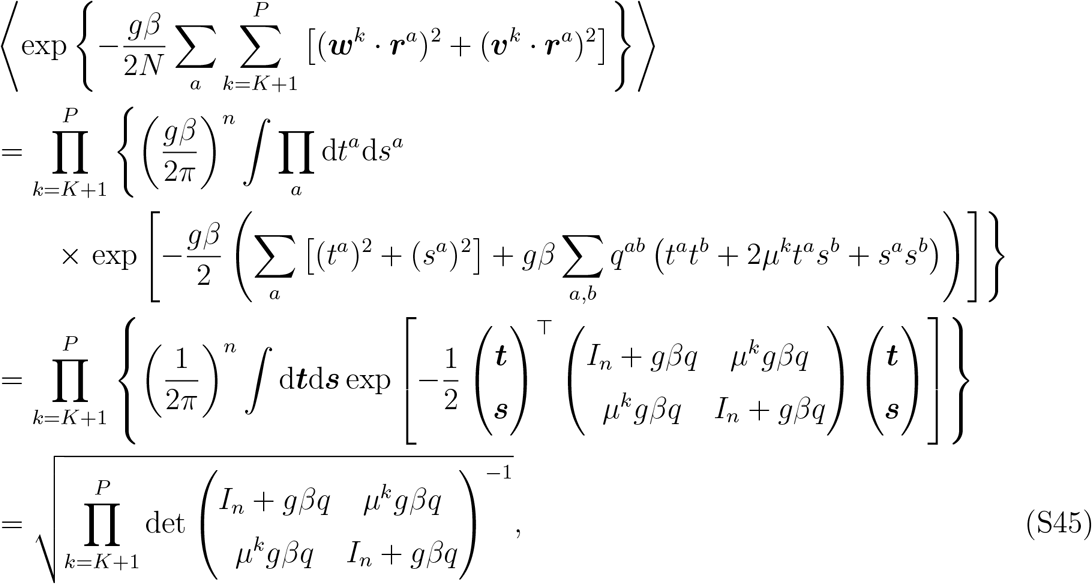

Here *q* is an *n* × *n* matrix [Eq. (S44)] and *I*_*n*_ is the *n* × *n* identity matrix. In the next to last step of Eq. (S45) we rescaled the integration variables *t, s* by 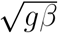.

The term in Eq. (S38) corresponding to the *K* presented pairs can be calculated in a similar fashion, which yields,

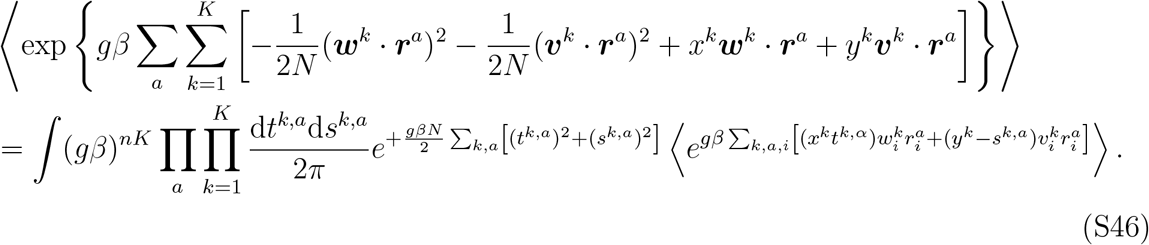

We introduce the delta function to enforce the definition of the order parameter *q*,

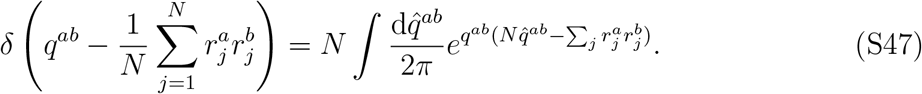

Putting all terms together Eq. (S38) gives,

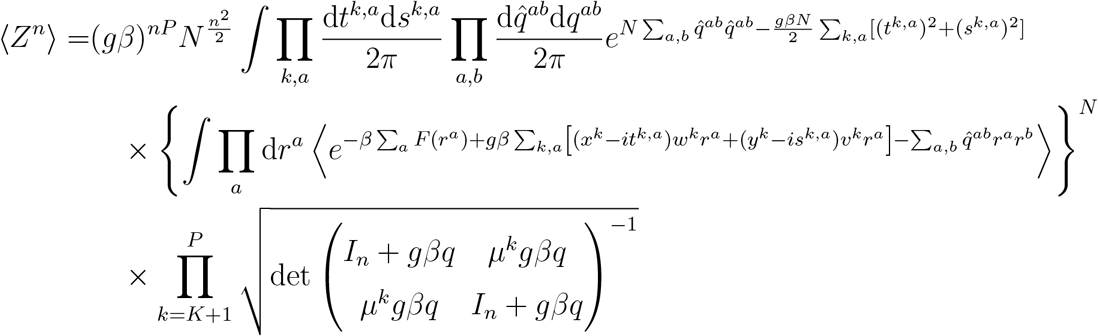

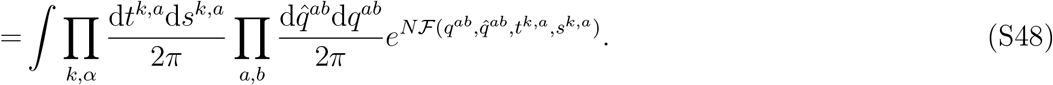

In the last line we have defined ℱ (*q*^*ab*^, 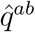, *t*^*k,a*^, *s*^*k,a*^) as,

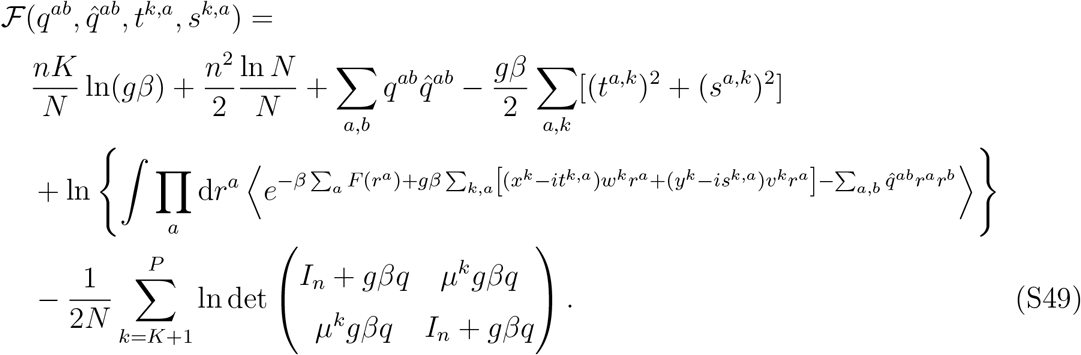

In the limit *N* → ∞, we use the saddle point approximation to compute the integral in the last line of Eq. (S48). Furthermore, because the Lyapunov function *E*_0_ is convex [Eq. (S33)], the saddle point solution is replica symmetric, i.e.,

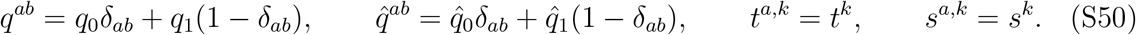

We then simplify the terms in ℱ,

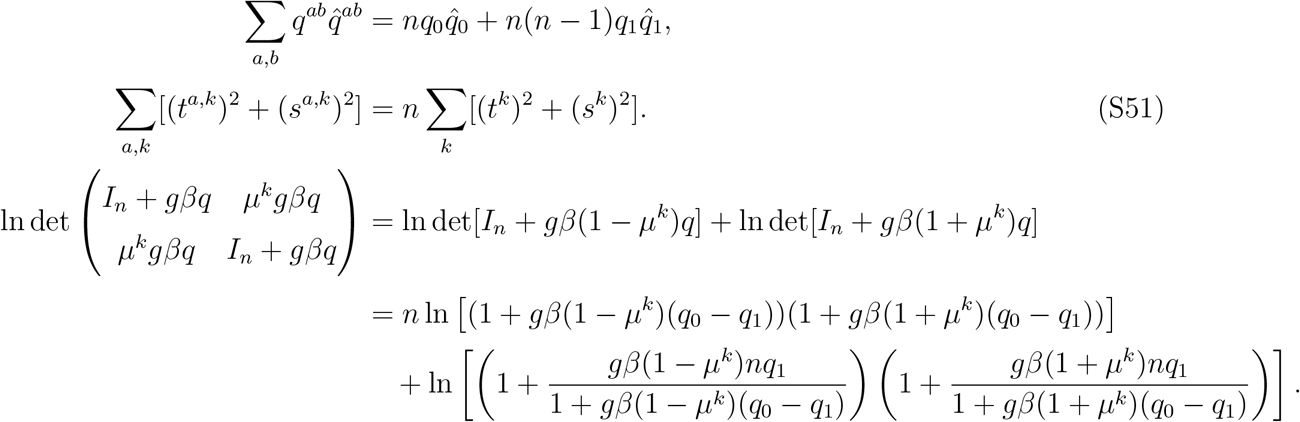

Simplifying the term in the third line of Eq. (S49) requires a number of additional steps. Using the integral representation of Gaussian function we write,

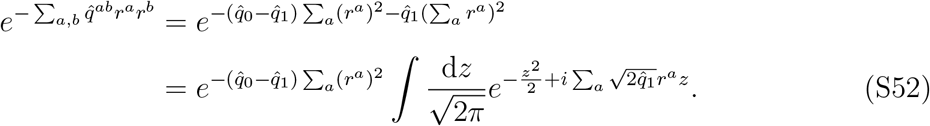

Substituting this into the integral in Eq. (S49) gives,

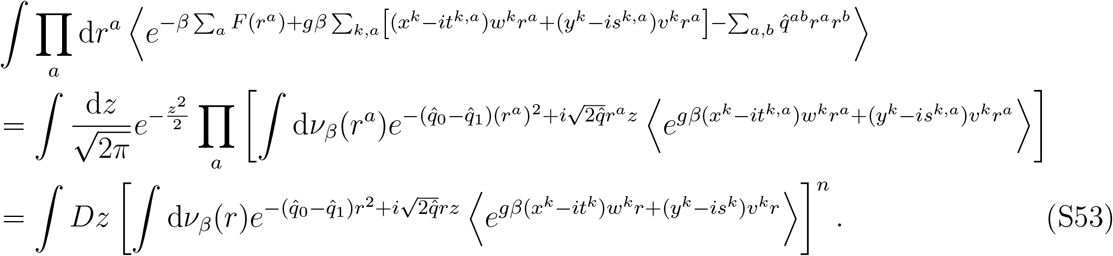

Here we have introduced the notation,

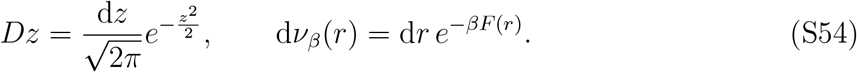

Therefore, under the replica symmetric ansatz, Eq. (S49) becomes

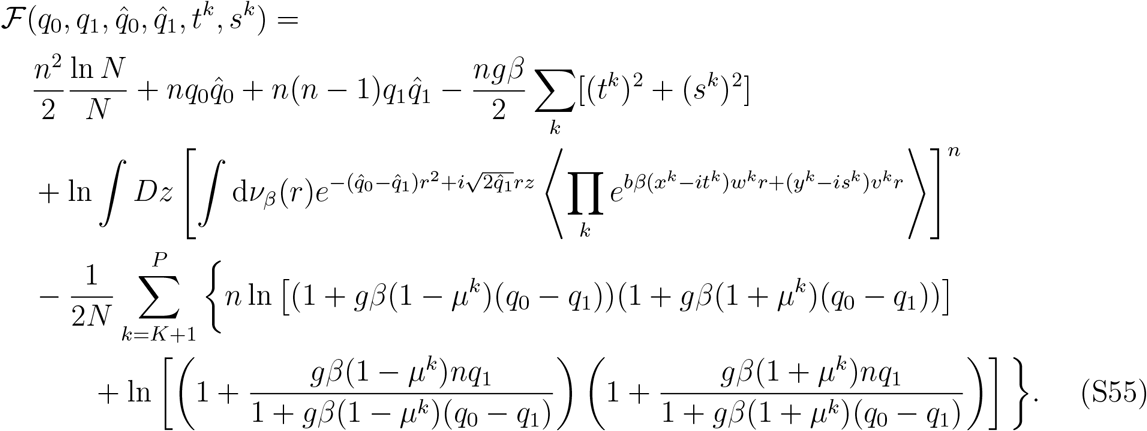

Now we take the limits *P, N* → ∞ and *n* → 0, and identify *α* = *P/N*, which gives,

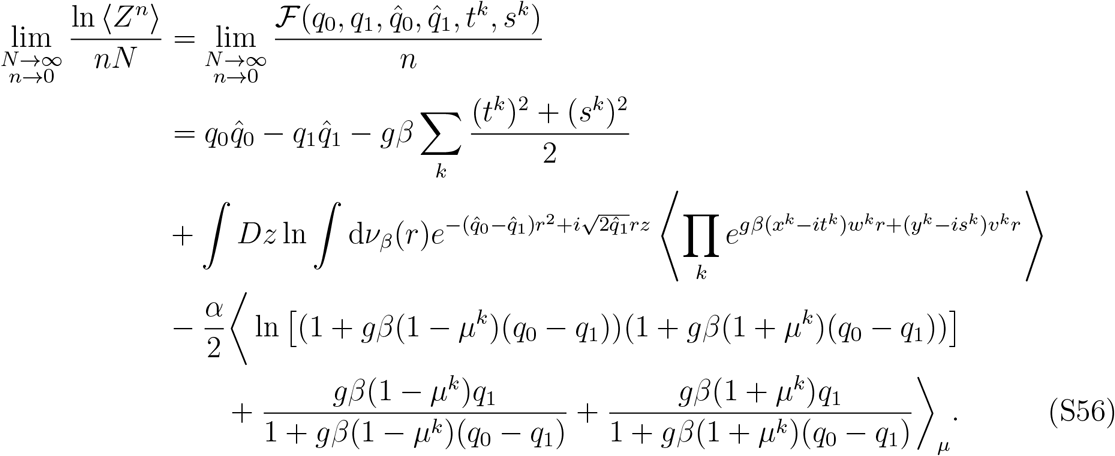

In the third line of Eq. (S56) we used the fact that for a well behaved function *A*(*z*),

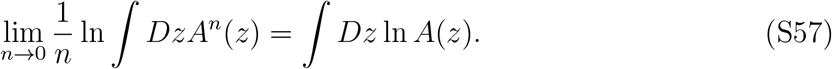

The last term in Eq. (S56) (proportional to *α/*2) was obtained by taking the limit over *P, N* → ∞ and introducing *α*, and assuming that the number of presented stimulus-pairs *K* is finite (necessary for the neural activity to remain finite, justified below). The average (*· · ·*)_*µ*_ is over the distribution of correlation values *µ*^*k*^, *k* = 1, …, *P* (i.e., over all learned stimulus-pairs).

To simplify the calculations, we define new variables

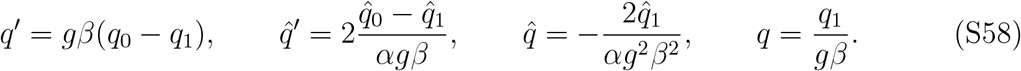

and further make the change of variables, *t*^*k*^ → *it*^*k*^ and *s*^*k*^ → *is*^*k*^. With these simplifications, we rewrite Eq. (S56) as,

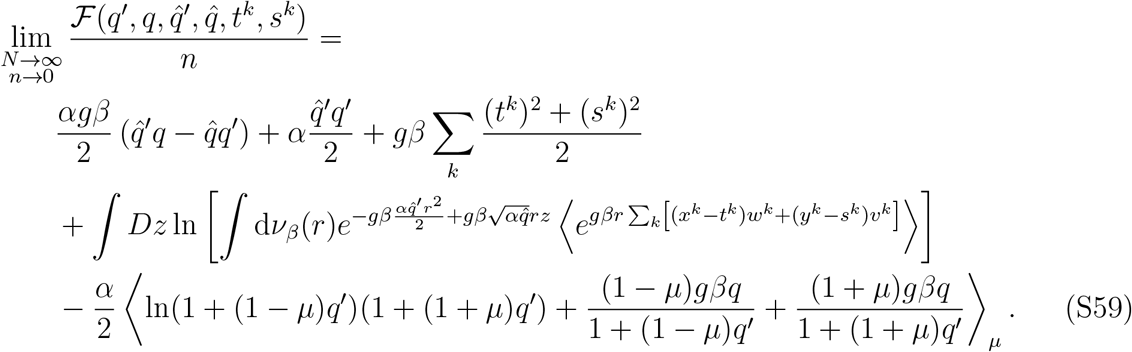

To extract information about the network’s response properties as *N* → ∞, we evaluated these expressions at the saddle point of 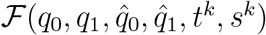. The saddle point satisfies,

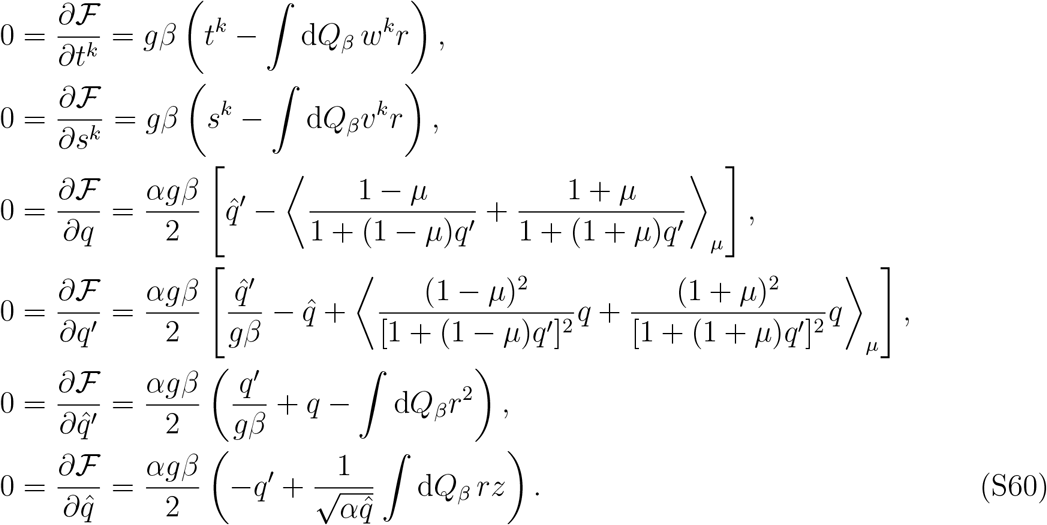

Here we have defined the probability measure d*Q*_*β*_ as,

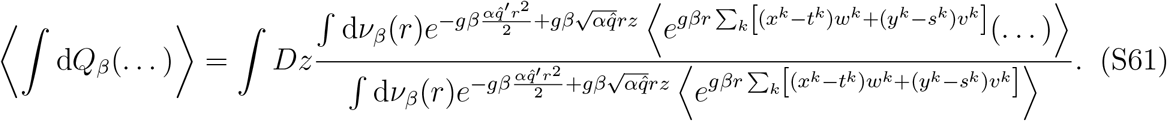

Indeed, the probability measure d*Q*_*β*_ contains a Boltzmann distribution with the corresponding Hamiltonian,

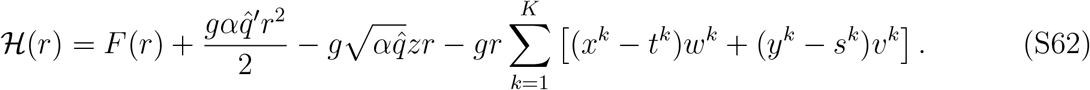

In the limit *β* → ∞ (‘zero temperature’), the Boltzmann distribution is dominated by the minimum of ℋ (*r*) (i.e., the ‘ground-state’). Since ℋ(*r*) is strictly convex, there is a unique minimum *r*^⋆^ ≥ 0.

If *r*^⋆^ > 0, the ground state satisfies ‘_*t*_(*r*^⋆^) = 0, or equivalently,

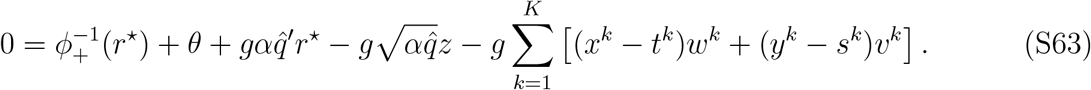

Otherwise, *r*^⋆^ = 0. Indeed, the two cases can be written in a compact way,

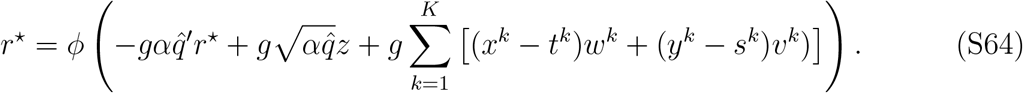

The solution of above equation defines a function *r*^⋆^(*w*^*k*^, *v*^*k*^, *z*). We recognize the argument of *ϕ* as the total input to each neuron and *r*^⋆^ as its nonlinear firing-rate response. It is important to note that the solution *r*^⋆^ depends on the Gaussian integration variable (*z*), the random synaptic weights (*w, v*), and the variables indicating the stimuli being presented (*x, y*), so overall the saddle point equations are expected to give a *distribution* of firing-rates, not a single value.

Substituting the ground-state solution *r*^⋆^ into the saddle point equation, we get at *β* → ∞,

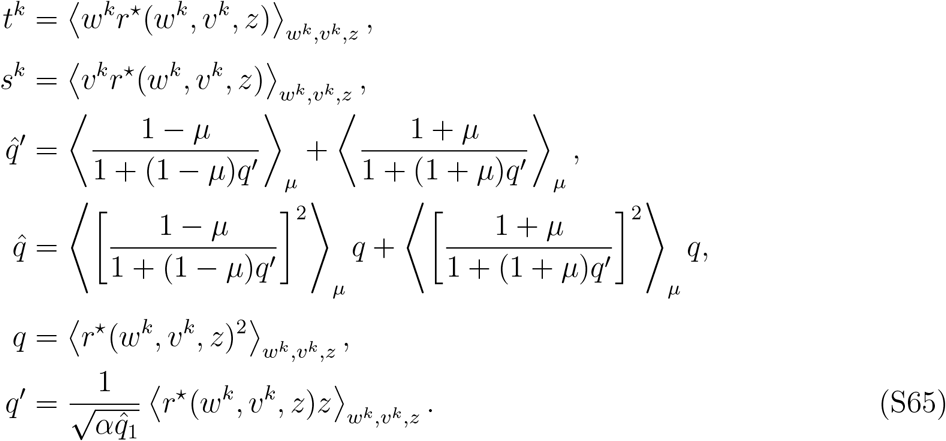

Notice that the order parameters *t*^*k*^ and *s*^*k*^ coincide with the internal predictions 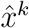 and 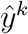 [Eq. (S1)], and the order parameter *q* is the second moment of the firing-rate distribution.

#### 2.2 Single-neuron and population statistics

We summarize the main results obtained from the above calculations: Given the distribution of synaptic weights {*w*^*k*^, *v*^*k*^} and a standard normal random variable *z*, the firing-rate distribution *p*(*r*) is the same as the distribution of the ground-state firing-rate *r*^⋆^(*w*^*k*^, *v*^*k*^, *z*) [Eq. (S34)]. The order parameters *q, q*^*t*^, 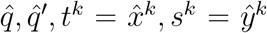 which appear in *r*^⋆^(*w*^*k*^, *v*^*k*^, *z*) need to be solved from the saddle point equations [Eq. (S60)]. Moreover, the voltage distribution of the neurons in the network is simply the distribution of the argument of the firing-rate transfer function *ϕ* in Eq. (S64), i.e.,

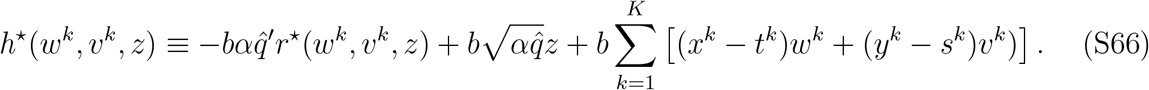

Below we restrict our analysis to the special case where {*w*^*k*^, *v*^*k*^} follow a multivariate Gaussian distribution; all the stimulus-pairs are learned equally well *µ*^*k*^ = *µ*; and the activation function is ReLU, *ϕ* = [*x* − *θ*]_+_.

##### 2.2.1. The high-dimensional case, P/N → α > 0

Under the above assumptions, Eq. (S34) can be solved exactly, giving neurons’ firing-rate and voltage distributions,

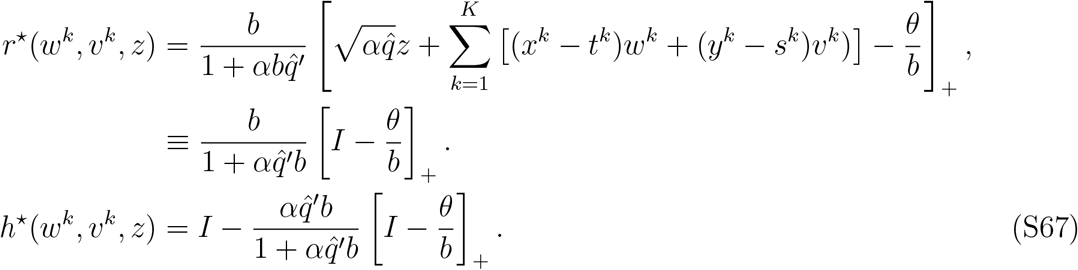

For convenience, we denote the Gaussian variable 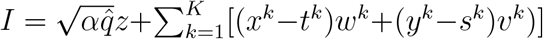. Each neuron receives input with mean 0, and variance (denoted *σ*^2^) that depends on the stimuli presented – how many, and whether they are matched or mismatched. From the above equation we see that neurons’ firing-rates follow a truncated Gaussian distribution. The saddle point equations [Eq. (S65)] can be simplified into,

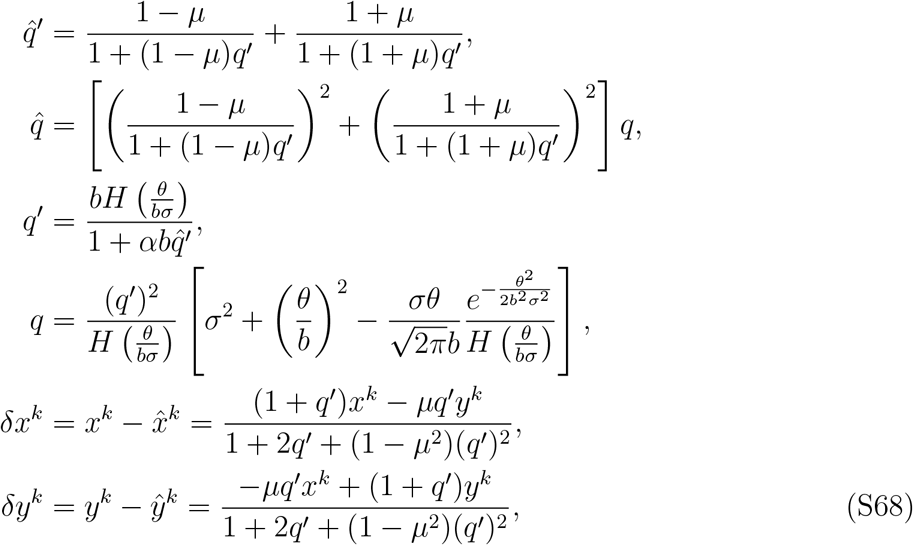

where 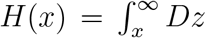 is related to the complementary error function. Since *q*′ ≥ 0 and *x*^*k*^ = *y*^*k*^ = 1 in the match condition, *δx*^*k*^ = *δy*^*k*^ ≥ 0. The variance *σ*^2^ in the above equations is given by,

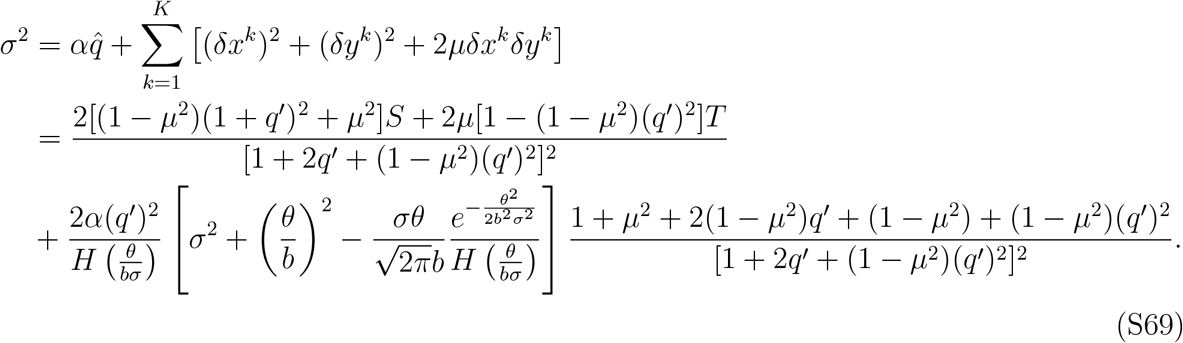

Here, we define variables that quantify the number of stimuli presented and whether their presentation is matched or mismatched: 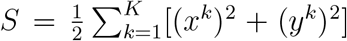 and 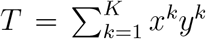. Combining Eq. (S69) with the first and third lines of Eq. (S68) gives a solution for *σ, q*′, 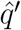. By substituting these into the other saddle point equations, we get all the order parameters.

In this case, the mean and variance of the firing-rate distribution are,

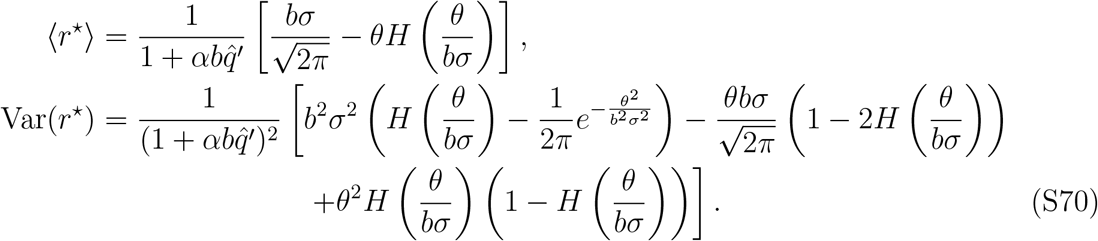

##### 2.2.2. The case *α* → 0

When *α* → 0, the saddle point equations reduce to,

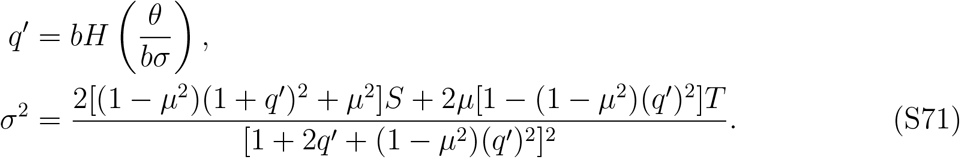

Once the values of *q*′ and *σ* are obtained from Eq. (S71), other order parameters in Eq. (S68) can be computed directly. Note that when *θ* = 0, then *q*′ = *b/*2. For a general threshold value *θ* ≥ 0, *q*′ is proportional to the gain parameter *b* and can thus be regarded as an order parameter quantifying the ‘effective gain parameter’ in the network. We see from Eq. (S71) that *q*′ depends on *σ*, which is scaled in turn by the quantities measuring the total stimulus strength, *S* and *T*. Thus, the changes of *q*′ in the match versus mismatch condition can be viewed as a global gain component in the predictive signal.

The single neuron firing-rate [Eq. (S34)] is now,

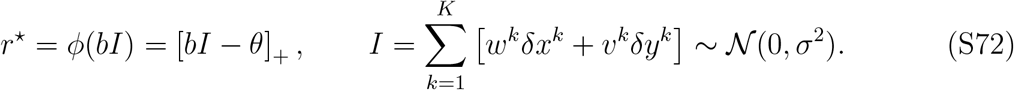

Notice that the mean and variance of the firing-rate [Eq. (S72)] can be obtained from Eq. (S70) by setting *α* = 0, and that the variable *I* in this case coincides the voltage level of neurons in the network [Eq. (S67)]. These results are used to generated the firing-rate statistics in Fig. 1.

In the case where only one stimulus-pair is presented (*K* = 1), the Pearson correlation between firing-rate vectors in the mismatch and match conditions can be calculated as follows. We denote by *I*_*x*_, *I*_*y*_, *I*_*xy*_ the voltage levels in the *x*-only, *y*-only mismatch and match conditions, respectively. The *I*’s are multivariate Gaussian variables with mean 0. We computed the correlations between inputs to neurons in the different mismatch conditions 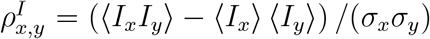 and between the mismatch and match conditions 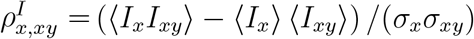. Here 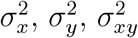, are the variances of *I*_*x*_, *I*_*y*_, *I*_*xy*_, respectively. We found,

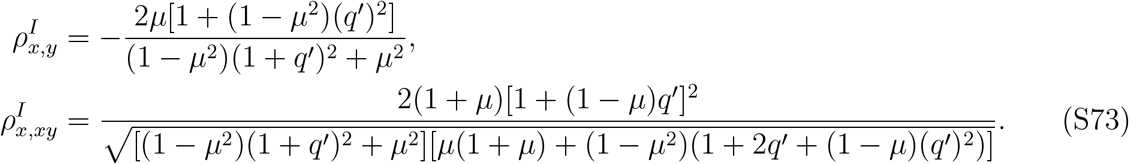

From the symmetry in the model we have 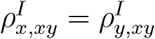.

In most cases, the experimentally accessible quantity is the firing-rate rather than the input current, so we also computed the Pearson correlation between firing-rates. We denote this correlation as 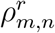, where *m, n* can refer to the conditions *x, y, xy*, and write its formal definition,

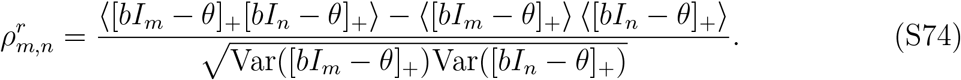

When *θ* = 0, the cross covariance between firing-rates can be worked out as,

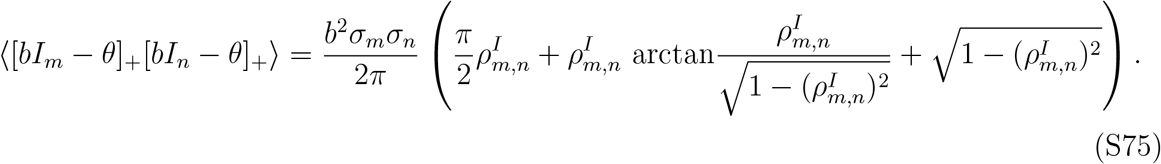

Together with the firing-rate mean and variance [Eq. (S70)], we obtained an explicit expression at for the firing-rate Pearson correlation, 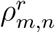. In the case of *θ* = 0, Eq. (S74) becomes,

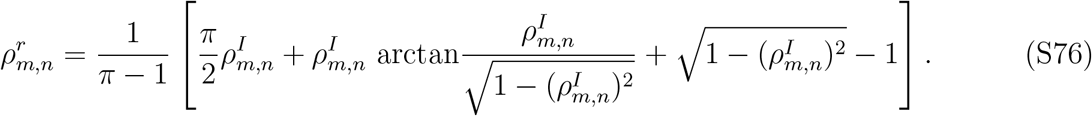

#### 2.3. Balance level distribution

The balance level for neuron *i* in the network is defined as,

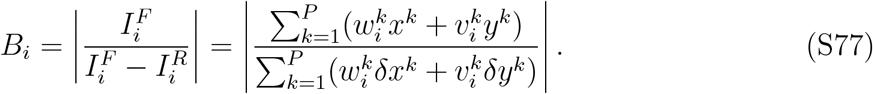

The *B*_*i*_’s are i.i.d. random variables for each *i*. Here the denominator is the net input 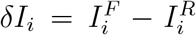 to neuron *i*, i.e., the difference between feedforward and recurrent input currents,

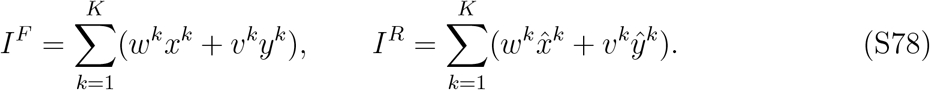

From Eq. (S66), *δI* can be expressed as

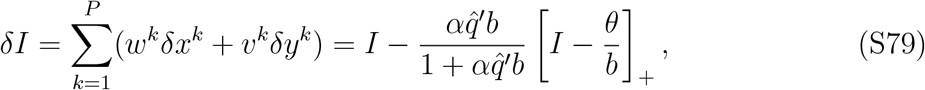

where 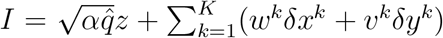 is defined in Eq. (S67). To simplify the notation we drop the subscript *i* from *δI*. Thus, to sample from the distribution of balance levels, one can first sample (*w*^*k*^, *v*^*k*^, *z*) from their corresponding distributions and then compute *I*^*F*^ and *δI*. The ratio between *I*^*F*^ and *δI* gives a sample of the balance level.

When the synaptic weights have Gaussian distribution and *α* = 0, the pair (*I*^*F*^, *δI*) is jointly Gaussian,

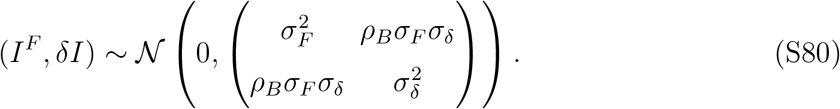

The coefficients of the covariance matrix of (*I*^*F*^, *δI*) are,

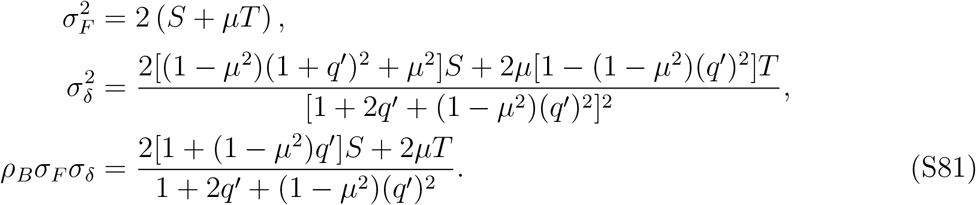

The balance level in this case can be expressed using a Cauchy random variable *ξ* as,

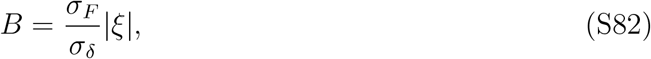

where the probability density function for *ξ* ∈ ℝ is,

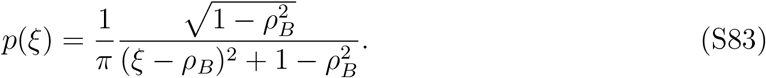

This result means that the average of the balance level distribution diverges. We use the quantiles to measure the magnitude of the balance level in the network (Fig. 2).

### 3. CHARACTERIZING DIFFERENT FUNCTIONAL NEURON TYPES

#### 3.1. Firing-rate correlations from two-body replica calculations

In this section we compute the probabilities of single neurons belonging to the different functional cell types for *two* stimulus-pairs. Since the stimulus-pairs and the neurons are statistically equivalent, we focus on the responses of neuron *i* to the first two stimulus-pairs, 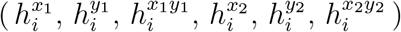. To mathematically characterize those voltage responses, we consider the joint distribution of the neurons’ firing-rates in two different stimulus conditions,

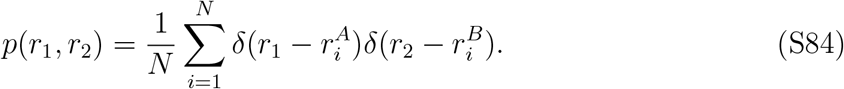

The superscripts *A, B* denote the stimulus conditions, i.e., *A* and *B* are chosen from {*x*_1_, *y*_1_, *x*_1_*y*_1_, *x*_2_, *y*_2_, *x*_2_*y*_2_}. We will show that at the limit *N* → ∞, the joint distribution for all different combinations of stimulus conditions can be obtained from the calculation of pairwise firing-rate correlations [Eq. (S84)].

To evaluate Eq. (S84), we consider two identical networks driven by different stimulus inputs. The energy function of the 1st system with firing-rates ***r***^*A*^ is,

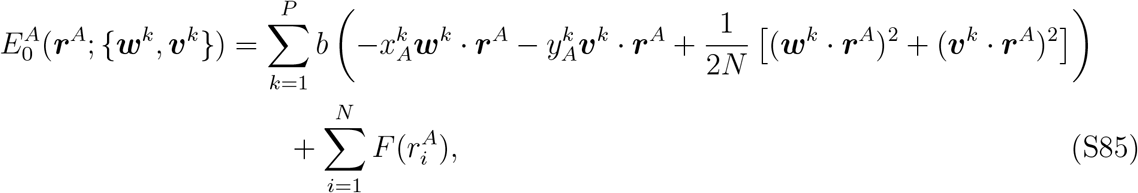

and similarly for the energy function of the 2nd system, 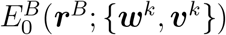. Note that in stimulus conditions *A* and *B*, only the first two stimulus-pair inputs are nonzero.

The partition function of the whole system is defined as

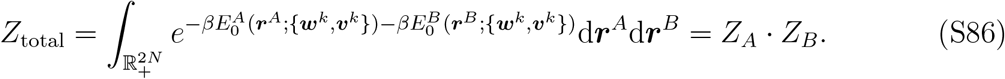

Again we use the replica trick,

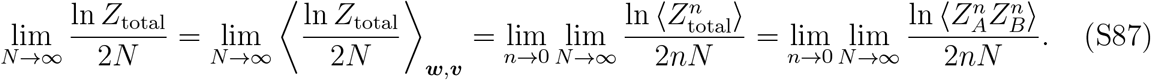

Note that the neural activities ***r***^*A*^ and ***r***^*B*^ of the two *separate but identical* networks are in fact statistically coupled due to the replica-average over (***w***^*k*^, ***v***^*k*^). The calculation for 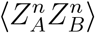 is similar to the one shown in §2. We denote the order parameters under replica symmetric ansatz as,

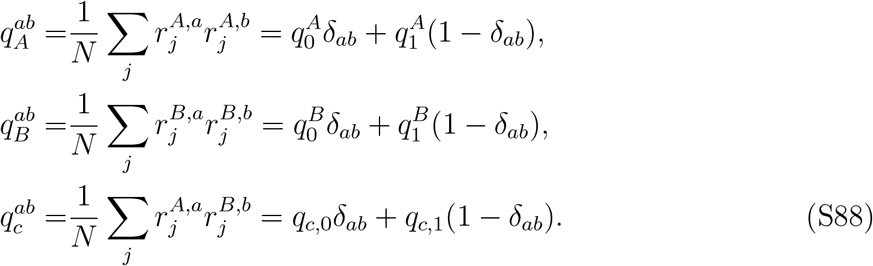

The last order parameter represents the overlap between replicas in system *A* and system *B*. Thus, the calculation of firing-rate correlations is very similar to the one-step replica symmetry-breaking calculation where the overlap between replicas within the same system is different from the overlap between the systems [79].

In the *N, P* → ∞, *P/N* → *α, n* → 0 limit, with similar changes of variables as before [Eq. (S58)], we write the result of the calculation as,

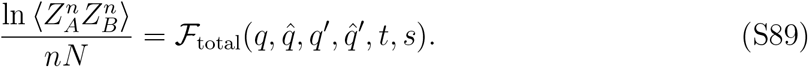

Each order parameter in the function ℱ_total_ has 3 components. For example, *q* has the components (*q*_*A*_, *q*_*B*_, *q*_*c*_). The calculation gives the function ℱ_total_,

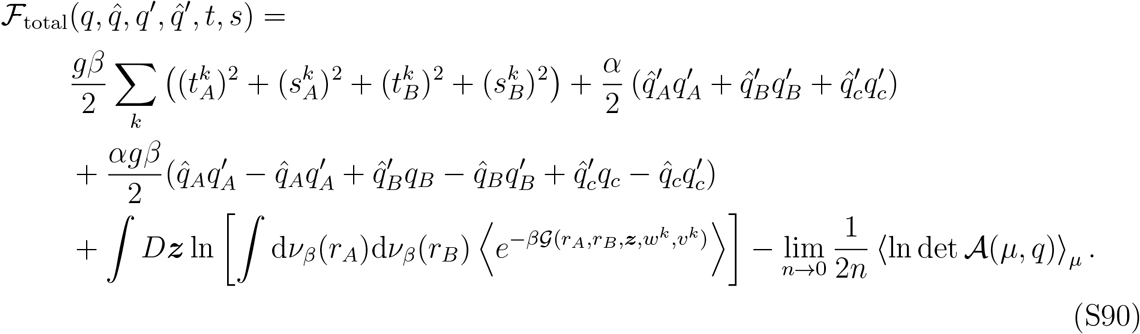

We introduced the functions,

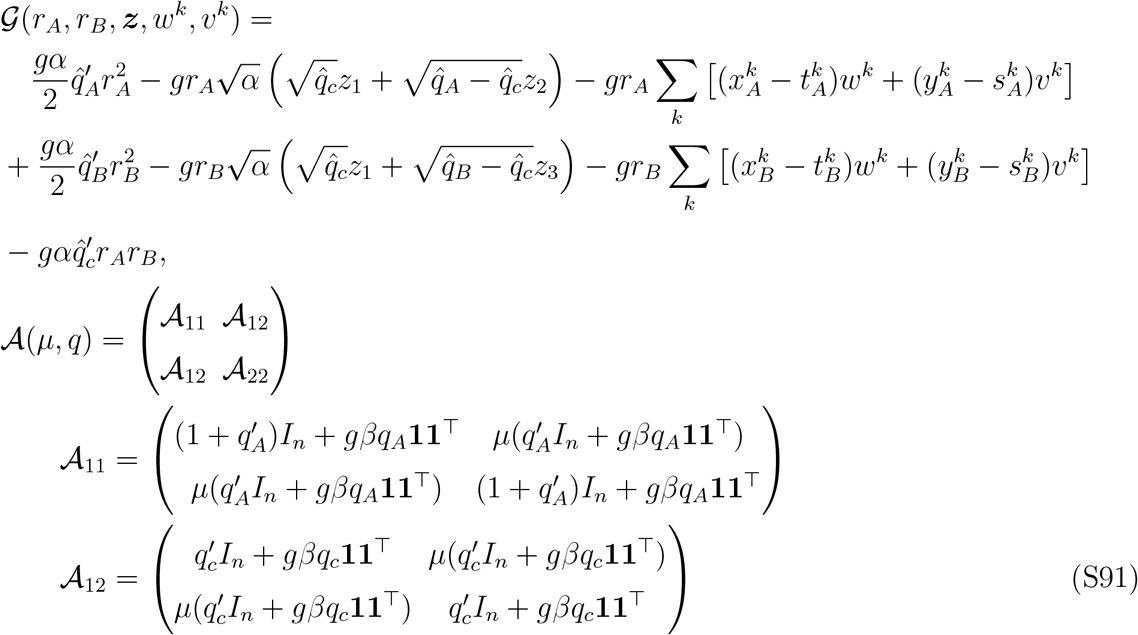

From symmetry, 𝒜_22_ is obtained by replacing *A* ↔ *B* in 𝒜_11_. Here, *I*_*n*_ is the *n* × *n* identity matrix and **1** is an *n*-dimensional vector of 1’s. All the order parameters in Eq. (S90) should be evaluated at the saddle point, in the limit *β* → ∞. We obtain these order parameters as follows.

First, we find the Hamiltonian corresponding to this system,

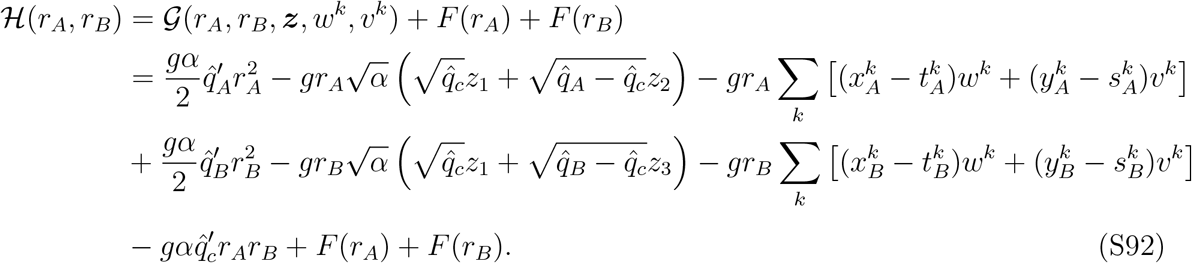

The extra terms *F* (*r*_*A*_) and *F* (*r*_*B*_) come from the probability measure d*ν*_*β*_. When *β* → ∞, the unique minimum 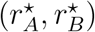 is given by,

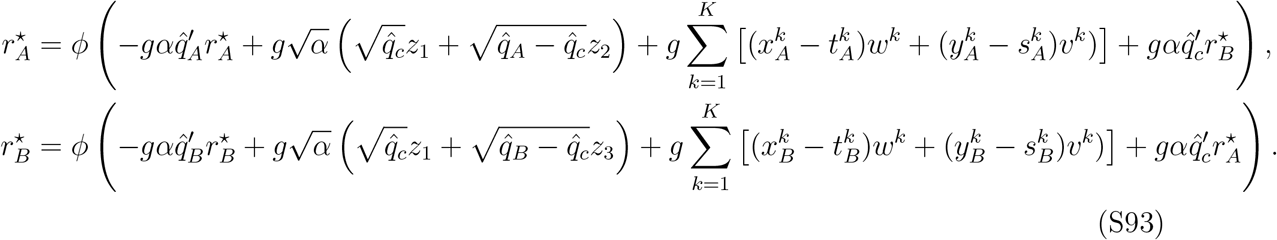

At the saddle point, the derivative of ℱ_total_ [Eq. (S90)] with respect to 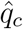 is set to 0, giving

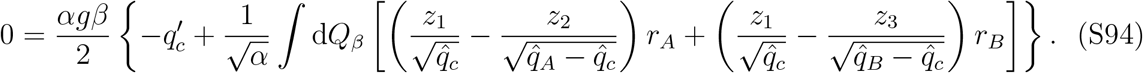

In the limit *β* → ∞ we find that,

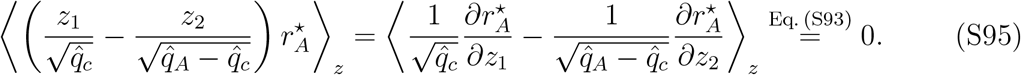

Similarly, the average over the term proportional to 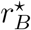 in Eq. (S94) is also 0. Substituting this into Eq. (S94), we get at the saddle point,

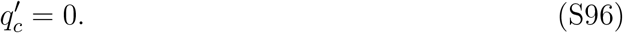

Next, we simplify the determinant of 𝒜 (*µ, q*). It is useful to write the submatrices as,

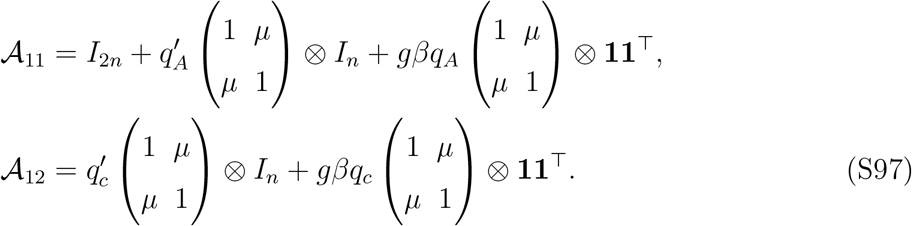

The symbol ⊗ denotes the Kronecker product between two matrices. For this product, we have the identity, (*A* ⊗ *B*)(*C* ⊗ *D*) = (*AC*) ⊗ (*BD*). Therefore the submatrices 𝒜_11_ and 𝒜_12_ commute and the determinant of 𝒜 (*µ, q*) becomes,

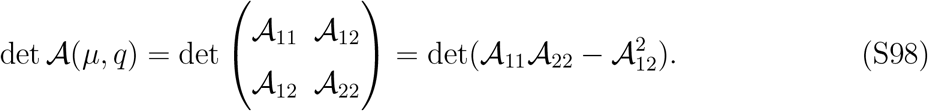

The two terms are equal to,

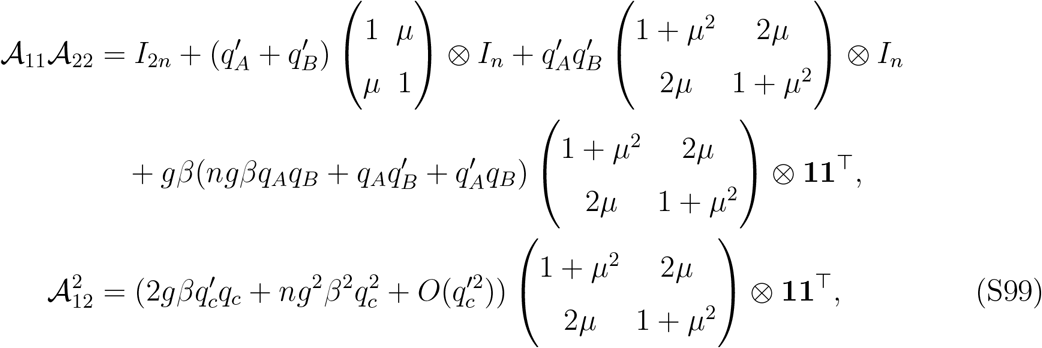

where we used 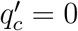. Note that we have kept the term linear in 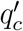 in 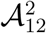, anticipating that we will need to evaluate derivatives with respect to 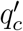 below. We find that the determinant has the following form,

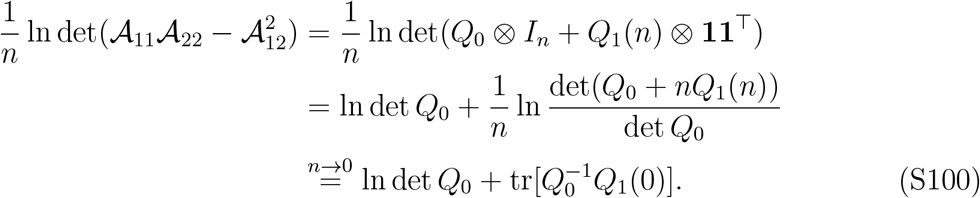

Here, *Q*_0_ and *Q*_1_(*n*) are 2 × 2 matrices that depend on the order parameters,

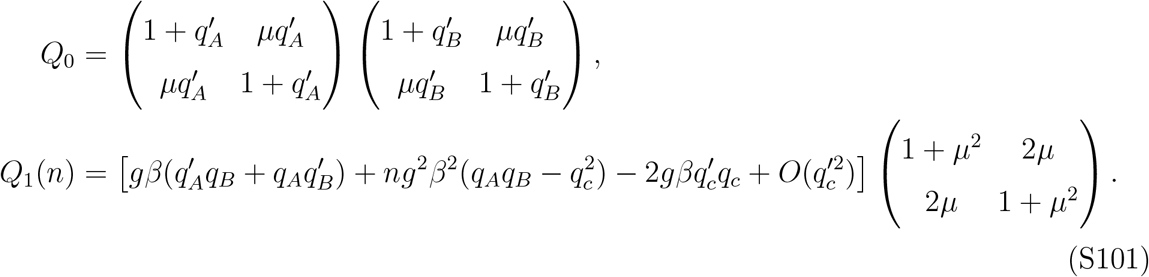

We use the above simplification to evaluate the derivative of ℱ_total_ [Eq. (S90)] with respect to *q*_*c*_ and set it to 0 at the saddle point,

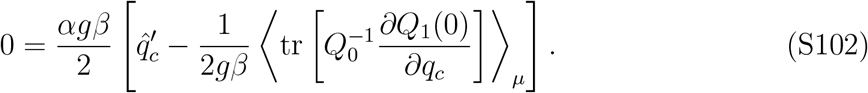

Using Eq. (S101), we find that 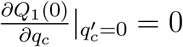. Therefore,

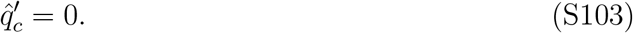

Substituting this result into Eq. (S93), we see this is consistent with the one-body replica results in Eq. (S34). Moreover, all the saddle point equations in the one-body scenario [Eq. (S60)] will hold in the two-body scenario. The two new equations when taking derivatives of ℱ_total_ with respect to 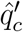 and 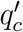 are,

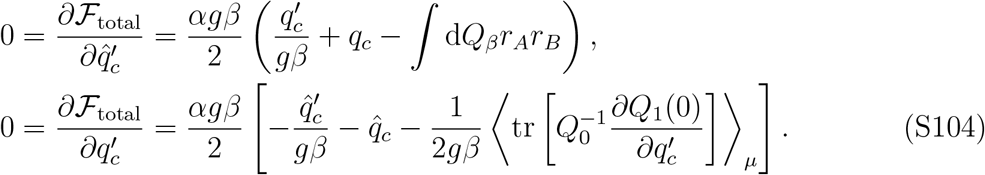

The second equation can be further simplified as follows. From Eq. (S101), we find

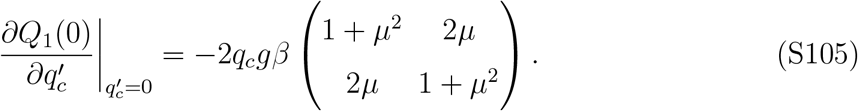

Combining with Eq. (S101), we get

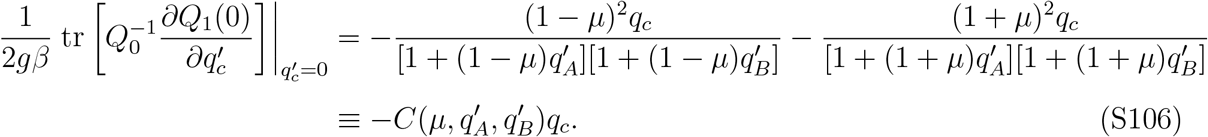

Therefore, using Eqs. (S104-S106), in the limit *β* → ∞, we get,

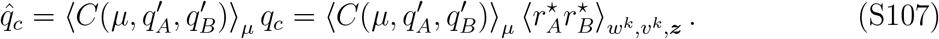

Below we consider the case where the activation function *ϕ* is ReLU and *µ*’s are the same for all learned stimulus-pairs. In this case 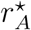 and 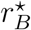 can be solved in closed form,

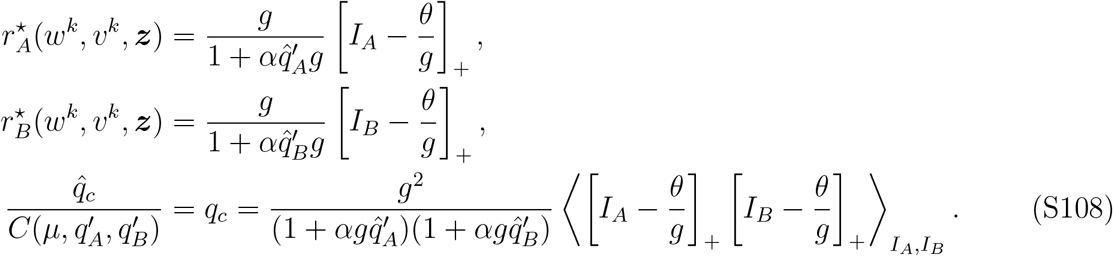

The random variables representing the currents can be read from Eq. (S93),

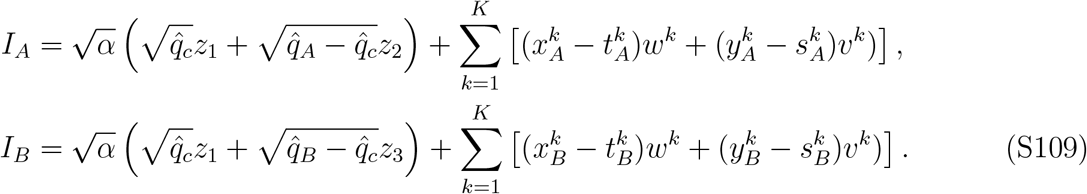

In summary, to obtain the joint distribution of neural activity under two stimulus conditions *A* and *B*, we first sample *w*^*k*^, *v*^*k*^, ***z*** from their corresponding distributions, and calcu-late *I*_*A*_ and *I*_*B*_ from Eq. (S109). The order parameters *q*_*A*_, 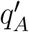, *q*_*B*_, 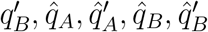 are then solved from the one-body replica equations [Eq. (S60)] and order parameters introduced in the two-body calculation 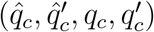 are obtained from Eq. (S108). Finally, 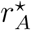 and 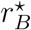 are calculated from Eq. (S93), which gives a random sample from the joint distribution.

The joint distribution of voltage levels can be computed similarly using the following formula for 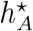,

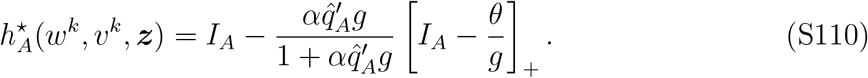

A similar formula holds for 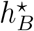 with the corresponding input current and order parameters. These results can be generalized to scenarios with more than two stimulus conditions. This joint distribution was be used to calculate the fraction of different functional neuronal types as shown in Fig. 3b,d.

#### 3.2. Explicit formulas in the Gaussian case

When the weights *w*^*k*^ and *v*^*k*^ are Gaussian, the input currents *I*_*A*_ and *I*_*B*_ are also Gaussian variables. Moreover, their variances 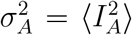 and 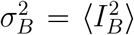 are given by the one-body calculation [Eq. (S69)]. We denote the input current covariance as *σ*_*AB*_ = ⟨*I*_*A*_*I*_*B*_⟩. This covariance is given by,

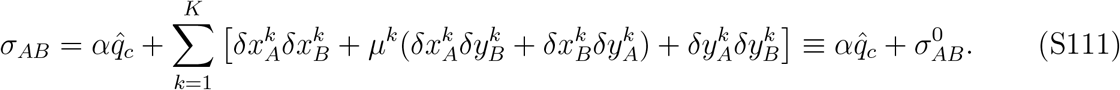

Here 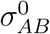 can be obtained from the one-body replica equations [Eq. (S60)].

Substituting *σ*_*AB*_ this into Eq. (S108) (i.e., averaging over the correlated Gaussians *I*_*A*_, *I*_*B*_), and defining *ρ*_*AB*_ = *σ*_*AB*_*/*(*σ*_*A*_*σ*_*B*_), we get a self-consistent equation for *ρ*_*AB*_,

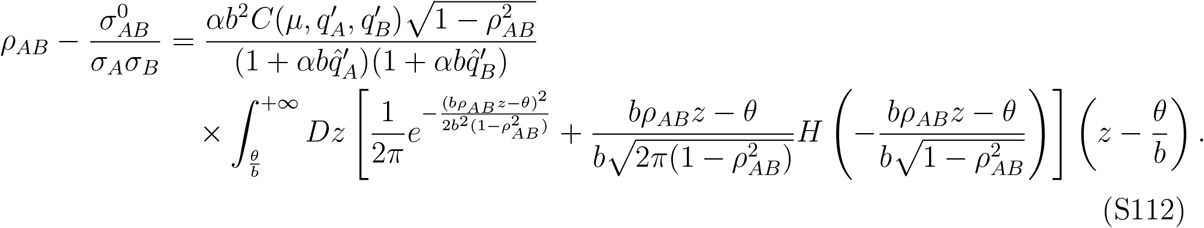

When *θ* = 0, the above equation for *ρ*_*AB*_ simplifies into

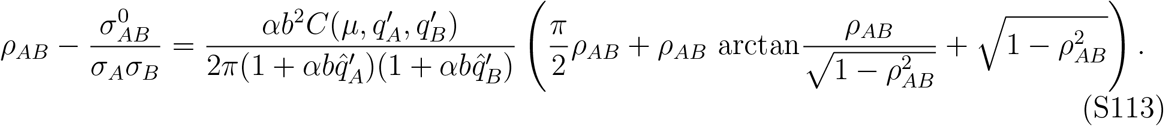

Note that the quantity *ρ*_*AB*_ calculated here is the high-dimensional counterpart of Eq. (S73), i.e., *ρ*_*AB*_ reduces to 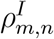 in the limit *α* → 0. We computed the firing rate correlations in the high-dimensional regime based on Eqs. (S74-S76) which give the correlation between the input current *ρ*_*AB*_.

The fraction of different functional neuronal types can also be obtained from the statistics 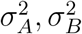 and *ρ*_*AB*_. Specifically, we set the stimulus conditions *A* =‘*x*-only mismatch condition’ and *B* = ‘match condition’. The fraction of *PE* and *R* neurons are defined as (Methods),

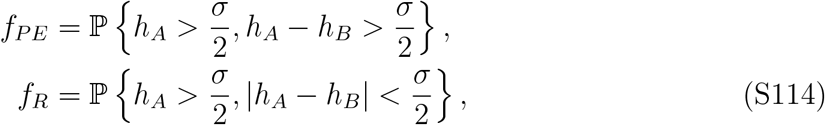

where *h*_*A*_, *h*_*B*_ are given by Eq. (S110). In the low-dimensional limit *α* → 0, *h*_*A*_ = *I*_*A*_, *h*_*B*_ = *I*_*B*_ and have multivariate Gaussian distribution. There the fractions of *PE* and *R* neurons have the explicit formulas,

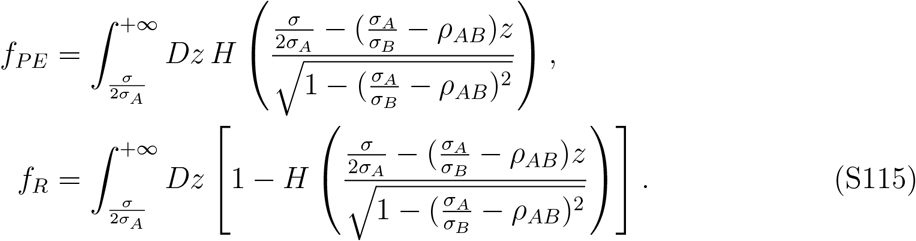

#### 3.3. Imperfect match of paired stimuli

We consider a network that learns a single stimulus association, and is presented with a ‘probe’ stimulus that is an imperfect match to the expected (learned) stimulus. This difference is modeled by letting the recurrent weight vector ***w*** be different from the feedforward weight vector ***w***′, giving the dynamics,

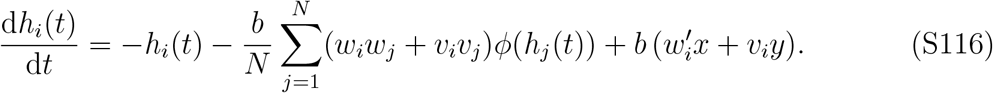

We used this model to understand recent experimental findings, where a motor-auditory association was learned, and animals were probed with sounds that differed from the learned tone [13]. We assume that the components of ***w***′ have mean 0 and unit variance [similarly to ***w*** and ***v***, Eq. (S9)], and the following cross terms,

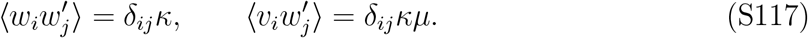

Here 0 ≤ *κ* ≤ 1 indicates the similarity between the learned stimulus input *x* and the one used as a probe. When *κ* = 1, the learned and probe stimuli are equal.

This network is very similar to the special case *α* → 0 of the network studied in §2.2.2. To understand its steady-state response, we use Eq. (S34) and define similarly,

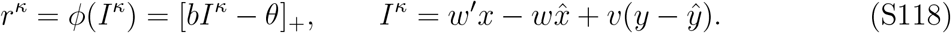

Here *ϕ* is assumed to be the ReLU function, 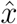 and 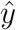 are the internal predictions [Eq. (S1)] and are given by the saddle point equations [Eq. (S68)],

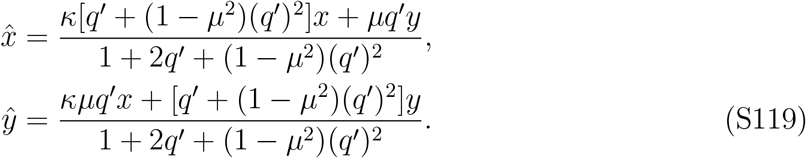

Note that we have modified them accordingly to account for fact that stimulus-pairing is ‘imperfect’. When all the weights have Gaussian distributions, the order parameters *q*′, *σ* satisfy [similarly to Eq. (S71)],

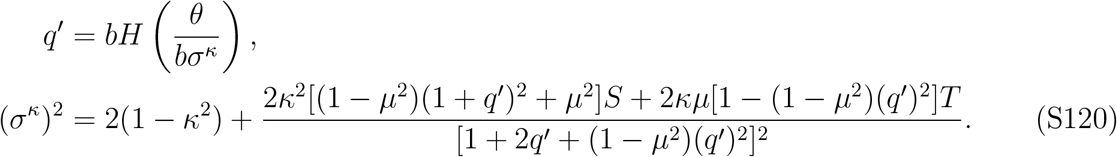

We computed the representation similarity between stimuli semi-analytically by first solving *q*′, *σ*^*κ*^, sampling *I*^*κ*^ from 𝒩 (0, (*σ*^*κ*^)^2^), and finally calculating the Pearson correlation coefficient [Eq. (S74)] between *r*^*κ*=1^ and *r*^*κ*^ for different values of *κ*.

To get the segregation index, we considered the difference between mismatch and match responses Δ for an arbitrary *κ* and *κ* = 1,

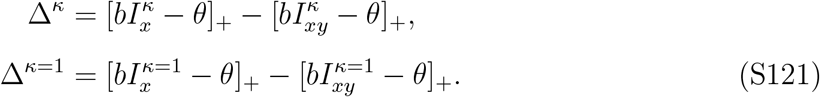

Note that 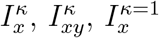 and 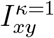 are random variables that depend on the random weights *w*′, *w* and *v*, order parameters 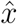 and 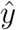, and the inputs *x* and *y*. The inputs were chosen according to the stimulus condition (match/mismatch). The segregation index (as a function of *κ*) is defined as the Pearson correlation between the two random variables Δ^*κ*^ and Δ^*κ*=1^, which is shown in Fig. 4f.

### 4. THE E/I NETWORK MODEL

#### 4.1. Derivation of the E/I connectivity in the model

We consider a network with two separate populations of excitatory and inhibitory neurons. The time-dependent voltages of *E* and *I* neurons are given by the following system of differential equations,

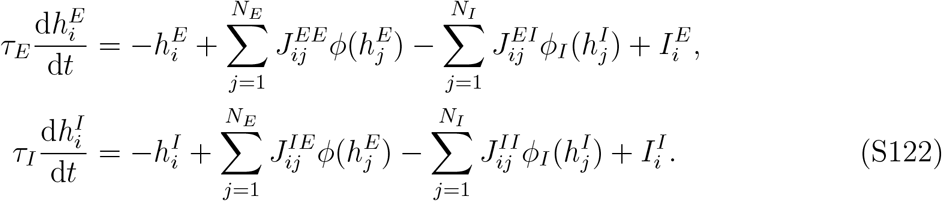

We assume that the activation function of inhibitory neurons is ReLU with threshold value equal to zero, *ϕ*_*I*_(*x*) = max{*x*, 0}. Notice the negative sign of the third term in both equations. This implies that the connectivity matrices *J*^*EE*^, *J*^*EI*^, *J*^*IE*^ and *J*^*II*^ are non-negative. We now derive these matrices, and the inputs *I*^*E*^ and *I*^*I*^, by matching the steady state activity of *E* neurons in the E/I network to the neural activity in the original network [Eq. (S4)]. At steady state, Eq. (S122) reads,

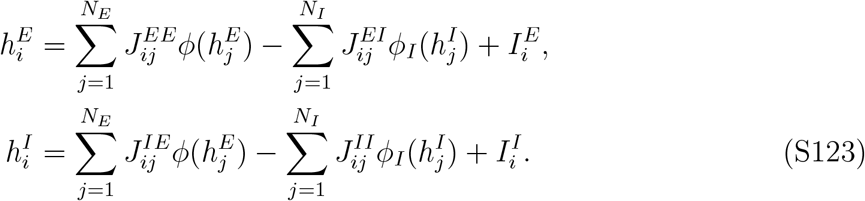

We restrict ourselves to choices of connectivity in which inhibitory neurons operate in the linear regime, i.e., 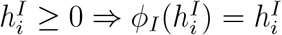. Substituting 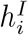 into 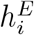 in Eq. (S123) we get,

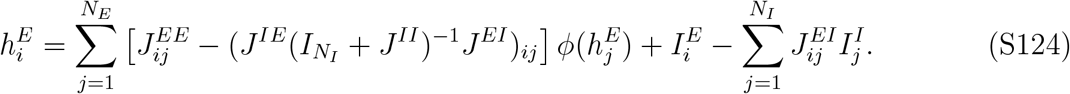

One can be check that the steady state solution is stable when *τ*_*I*_ *≪ τ*_*E*_. Here 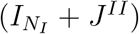 is assumed to be invertible. From now on we suppress the subscript *N*_*I*_ indicating the dimension of the identity matrix 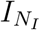. Equating this with the steady state in the original network [Eq. (S8)] gives the constraints on the connectivity and input,

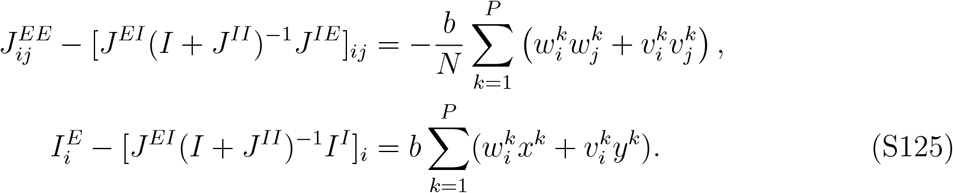

Following a scheme for separating E/I connectivity used in previous work [54], we define positive random variables 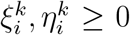 such that the variables 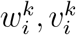 are retrieved when the mean is subtracted from the new variables. Mathematically,

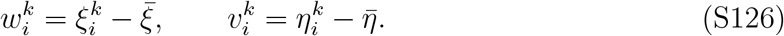

The means 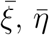 are chosen to be independent of the neuron and pattern indices *i, k*. Using the same trick as Ref. [54], the first equation in Eq. (S125) can be separated into two parts,

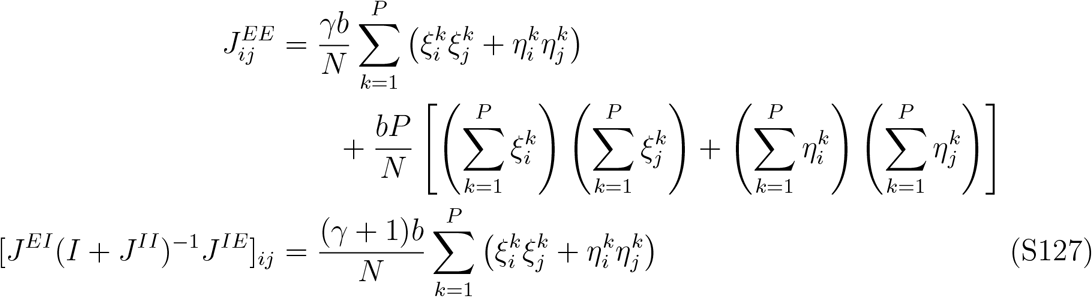

Here *γ* is an arbitrary positive number, which we set to 1 in all later results.

We make two additional assumptions: (*i*) ‘Feedforward’ stimulus input exclusively target excitatory neurons 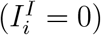; and (*ii*) *I*-to-*E* connectivity has the form 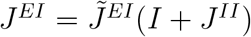, where 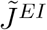 is a nonnegative matrix. Given these, Eqs. (S125, S127) become,

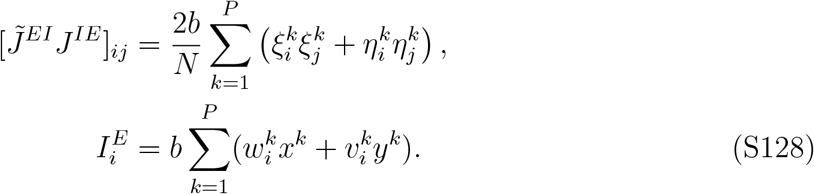

To obtain the E/I balance level for excitatory neurons in this network, we write the total excitatory input 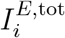 as the sum of different contributions,

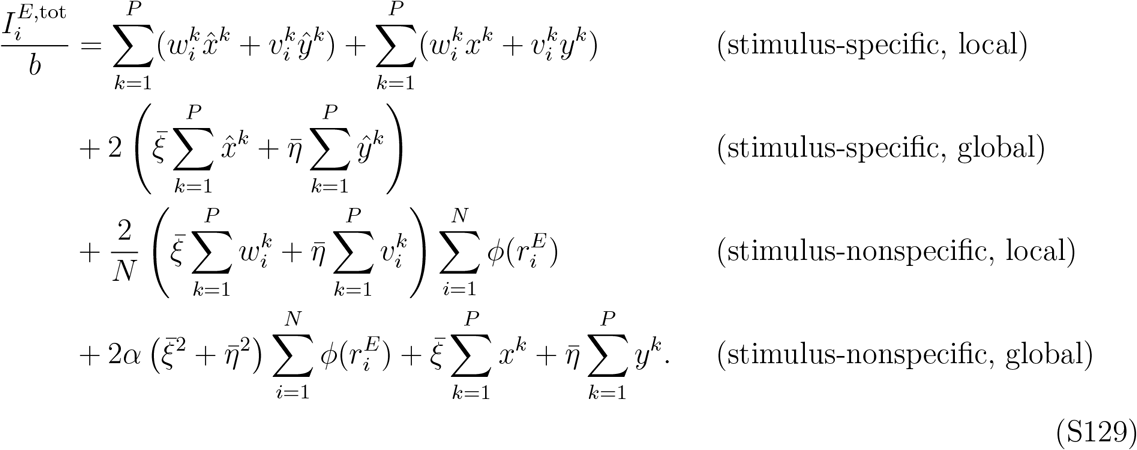

Taking the ratio between the stimulus-specific, local component and the net input to each excitatory neuron, we get,

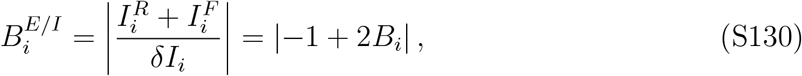

Where 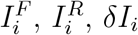, *δI*_*i*_ and *B*_*i*_ are those defined in the original network model [without separation of *E* and *I*; Eq. (S77)]. Therefore, for moderate values of *B*_*i*_ > 1*/*2, up to a scaling factor and shift, the stimulus-specific, local component of the E/I balance level is the same as the balance level we analyzed in Figs. 2, 3. Note that in the range of *α* values analyzed in Fig. 2, the fraction of neurons with *B*_*i*_ < 1*/*2 is negligible in both match and mismatch conditions.

#### 4.2. Interpolation via nonnegative matrix factorization

Solving for 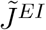 and *J*^*IE*^ in Eq. (S128) is equivalent to a nonnegative matrix factorization problem [53]. Using the shifted, nonnegative weight vectors, we define the matrices Ξ, *H, S*,

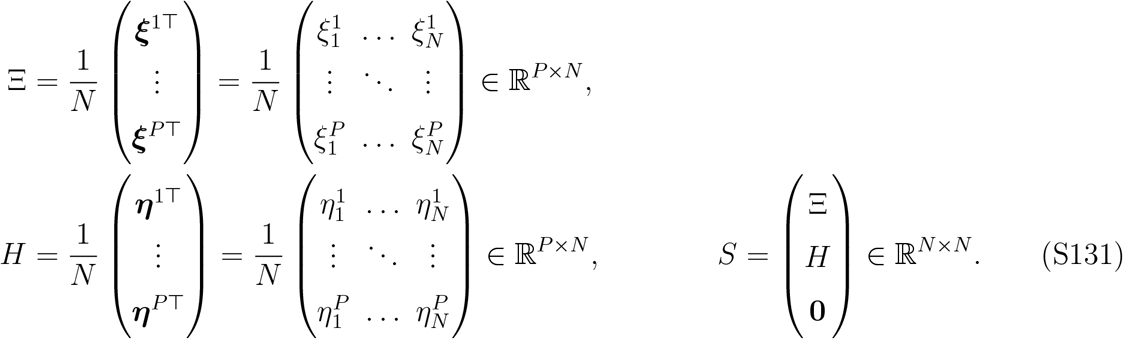

Throughout this section, we will assume 2*P* ≤ *N*, and ‘**0**’ pads with 0’s such that *S* is a square matrix. Thus, the connectivity equation [Eq. (S128)] can be rewritten as,

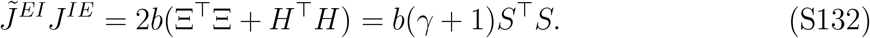

For each choice of a nonnegative matrix *J*^*IE*^, the above equation has a nonnegative solution *J*^*EI*^ if and only if the convex cone formed by the row vectors of *J*^*IE*^ contains the convex cone formed by the row vectors of *S* [formally denoted as cone(*J*^*IE*^) ⊇ cone(*S*)]. This condition can be derived from the definition of matrix multiplication [53]. Based on this condition, we identify a family of solutions {*J*^*EI*^(*λ*), *J*^*IE*^(*λ*)} parameterized by *λ* ∈ [0, 1] as follows. At one end, we choose *J*^*IE*^ equal to the identity (*J*^*IE*^(*λ* = 0) = *I*_*N*_). At the other end, *J*^*IE*^(*λ* = 1) = *S*^*t*^, where *S*^*t*^ is defined such that its first 2*P* rows are the same as the nonzero rows of *S* and the rest of its rows are randomly sampled from the vectors ***ξ***^*k*^*/N*, ***η***^*k*^*/N*. This ensures that cone(*S*^*t*^) ⊇ cone(*S*). This family of solutions assumes that the number of inhibitory neurons equal to the number of excitatory neurons.

The firing-rates of inhibitory neurons are given by,

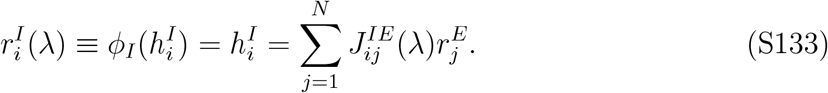

At the two ends, this reduces to,

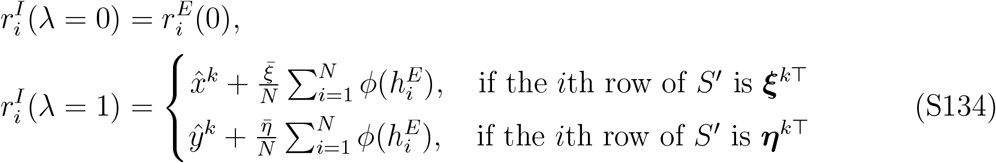

Based on these equations, we call *λ* = 0 the ‘private’ solution and *λ* = 1 the ‘internal prediction’ scenario. For *λ* = 1, the second term in 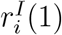 can be canceled by a global disinhibtory input. For intermediate *λ*’s, it may seem natural to choose a linear interpolation between the two solutions, *J*^*IE*^(*λ*) = *λJ*^*IE*^(1) + (1 − *λ*)*J*^*IE*^(0). We find however that this choice does not ensure that the solution for *J*^*EI*^ is nonnegative.

Instead, we choose *E*-to-*I* connectivity as follows. Two intermediate points within the segment [0, 1] are denoted as *λ* = 0^+^ and *λ* = 1^−^, thereby dividing the segment into three.

At those points we choose *J*^*IE*^ to be,

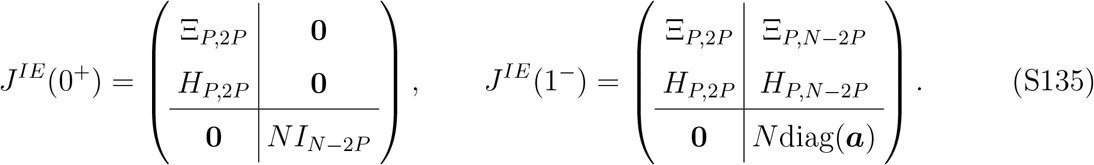

Here, Ξ_*P*,2*P*_, *H*_*P*,2*P*_ consist of the first *P* rows and first 2*P* columns of Ξ and *H*, respectively; Ξ_*P,N*−2*P*_, *H*_*P,N*−2*P*_ consist of the first *P* rows and last *N* − 2*P* columns of Ξ and *H*, respectively; diag(***a***) is a diagonal matrix, with diagonal elements given by the *N* −2*P* components of the vector ***a*** which is specified below. Again the **0**’s are used for padding.

The interpolation of *J*^*IE*^(*λ*) from *λ* = 0 to *λ* = 1 thus consists of three regions:

I. *λ* from 0 to 0^+^: The upper left block of *J*^*IE*^ changes from an identity matrix to a matrix of stimulus input vectors.
II. *λ* from 0^+^ to 1^−^: The upper and lower right blocks linearly interpolate the matrices shown in Eq. (S135). Results in the main text are taken from here.
III. *λ* from 1^−^ to 1: The lower part of the matrix changes to contain stimulus vectors.

We start with solutions in Region (II) which we found to be the most relevant to the empirical measurements in [12], since we estimated *λ* ≈ 0.6. Network properties for a range of *λ* values between 0 and 1 (Figs. 5, S6, S7, S8) are also based on the results in Region (II). The connectivity matrices *J*^*EI*^(*λ*) and *J*^*IE*^(*λ*) in Region (II) are given by,

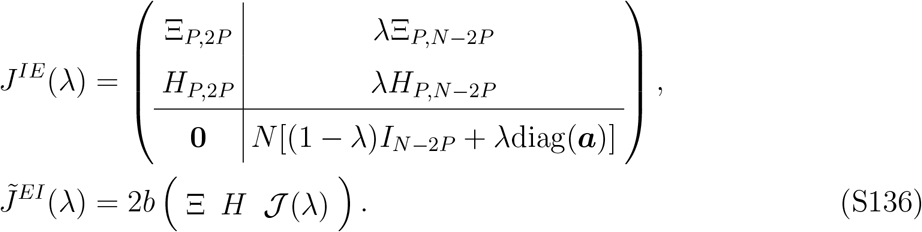

Here 𝒥 (*λ*) is a *N* × (*N* − 2*P*) matrix whose elements are given by

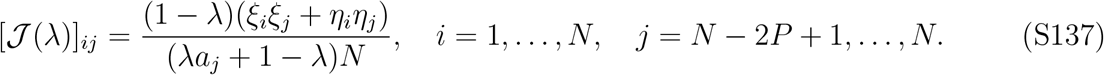

One can check that cone(*J*^*IE*^(*λ*)) ⊇ cone(*S*), and thus Eq. (S132) is satisfied and the elements of *J*^*EI*^ are nonnegative for every *λ*.

The interpolation in Region (I) requires smoothly ‘morphing’ the upper left block of the connectivity matrix involving Ξ and *H* to the identity matrix. This can be done by replacing the last row and last column with 0 and then setting the last diagonal element to be 1. Repeating this replacement *P* times yields the identity matrix. We note that in the low-dimensional case [*P* = *O*(1)], this procedure only changes the *E* connections to *P* out of *N* inhibitory neurons. Thus its effect on the overall statistics of inhibitory neurons’ activity is negligible. In the high-dimensional case [*P* = *O*(*N*)], the distributions of neural activity and synaptic weights themselves change smoothly along this interpolation path. Similarly, in Region (III), we replace every row in the lower part of the matrix with one of the randomly sampled vectors that appear in the matrix *S*′.

#### 4.3. Plasticity of inhibitory weights during learning

The interpolation solutions presented in the last section are valid for any set of positive real numbers *a*_*i*_, *i* = 2*P* + 1, …, *N*. In Fig. 5 we choose the *a*_*i*_’s as follows,

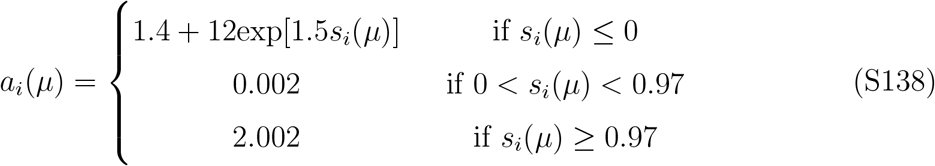

where

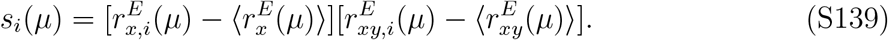

Here 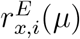 and 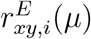 are the firing-rates of the *i*-th excitatory neuron in the *x*-only mismatch and match conditions for a given value of *µ*. 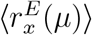 and 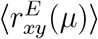 are the average firing-rates over all the *E* neurons in the two conditions. This mathematical form for *a*_*i*_ is chosen to match the experimental data on fast spiking neurons (Fig. 5c,d).

To track individual synapses during learning, we generate the *k*th stimulus input vectors ***ξ***^*k*^ and ***η***^*k*^ as follows: (1) We first generate two independent isotropic Gaussian vectors 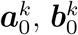, with mean equal to 3 and standard deviation equal to 1; (2) Then we form the a linear combination to generate two correlated Gaussian random variables,

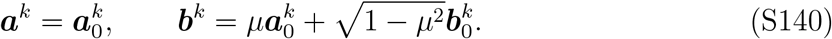

(3) Finally, we clip both variables to positive and define them as ***ξ***^*k*^ and ***η***^*k*^. In this case, the resulting vectors ***w***^*k*^ and ***v***^*k*^ [Eq. (S126)] will be approximately correlated Gaussian variables with mean 0. These procedures are used to produce the plots in Fig. 5c,e,f,g.

### 5. PARAMETER VALUES USED IN THE FIGURES

Unless specified, in all the main and supplementary figures, ***w***^*k*^ and ***v***^*k*^ have joint Gaussian distribution and satisfy Eq. (S9). The number of neurons in the network is *N* = 2000.

**Figure 1:** We set *α* = 0, *θ* = 0 and *b* = 150 throughout this figure.

Panel b: We use Eq. (S72) to generate *N* = 2000 samples of 2D random variables (*I*_*x*_, *I*_*xy*_) and compute the corresponding firing-rates.

Panel d: The theory lines for the Pearson correlation between different stimulus conditions are calculated from Eqs. (S71, S73, S76). The simulation points are calculated by sampling the neurons’ firing-rates as described in the Panel b caption. As each vector represents the mean-subtracted firing-rate vectors, the cosine of the angle is equivalent to the Pearson correlation coefficient between the original firing-rate vectors.

Panel f: The firing-rate distribution on the left (‘Our model’) is generated in the same way as in Panel b. As the neural responses to two stimulus-pairs are mutually independent at *α* = 0, the joint distribution is a product of the corresponding marginal distributions. The firing-rate distribution on the right (‘Segregated model’) is generated by using the same marginal distributions (as in the plot of ‘Our model’), but adding a nonzero correlation (which equals to 0.9) in the input variables (*I*_*x*_, *I*_*xy*_) that are used to calculate the firing-rates.

**Figure 2:** We set *θ* = 0 throughout this figure.

Panel b: *α* = 0, *b* = 150. The ‘Early’ and ‘Late’ plots for balance level distribution are calculated at *µ* = 0 and *µ* = 0.9 respectively.

Panel d: *α* = 0, *b* = 150. For SVM classification, stimulus inputs in the mismatch condition are generated from Gaussian mixtures centered at (0, 1) and (1, 0), both of which are isotropic and have variance 0.05. Similar Gaussian mixtures are used for stimulus input in the match condition, except that the centers are at (0, 0) and (1, 1). The SVM model is fitted using the Matlab function ‘fitcsvm’. The classification error is calculated via the matlab functions ‘crossval’ and ‘kfoldLoss’.

Panel e: This figure panel is an illustration and the parameters are *α* = 0, *µ* = 0.7.

Panel f: The threshold on the firing-rate for determining the optimal *b* is chosen such that at *α* = 0, the optimal balance level is the same as the one fitted to experimental data [20] in Figure 3 (*B*^⋆^ ≈ 162).

**Figure 3:** We fit both sets of experimental data [12, 20] using Eq. (8) (Methods).

**Figure 4:**

Panel b: *α* = 0, *b* = 150.

Panel c: The values of *b* in both plots are chosen to be at the optimal values.

Panel d: Plotted on the y axis is the fraction of mixed-representation neurons among all *PE* neurons for the stimulus pair 1.

Panel f: We set *b* = 189, which is the value extracted from the data [12]. The sparsity levels are defined as the fraction of active neurons in the network and changed by varying the firing-rate threshold *θ* in the network model. The threshold values corresponding to the three plotted curves are *θ* = 4.5, 6.5, 21.5.

**Figure 5:** We set *θ* = 0, *J*^*II*^ = 0 throughout this figure. Before and after learning correspond to *µ* = 0 and *µ* = 0.97. During learning, the functional cell types of a specific *E* or *I* neuron in the network might change. The cell-type-specific synaptic weight statistics shown in Fig. 5f,g only include synapses whose pre- and postsynaptic neurons maintain their identity throughout learning. Other parameter values can be found in §4.3.

**Figure 6:** *θ* = 0 throughout this figure. The number of neurons for each module is 400. All the error bars are computed based on 30 random samples of synaptic weight vectors. The steady state of the network is obtained by simulating the ODEs [Eq. (S29)] for total time *t* = 4.

Panel b : *α* = 0, *µ* = 0.97. The colormap indicates the firing rate averaged over all neurons in all modules in the *x*-only mismatch condition.

Panel c: *b*_1_ = *b*_3_ = 50 and *b*_2_ = 190 are the values at the star position in panel b. *µ* = 0.97.

Panel d-h: *b*_1_ = *b*_3_ = 50 and *b*_2_ = 190 are the values at the star position in panel b.

**Figure S1:**

Panel a: *α* = 0, *b* = 150.

Panel c: The threshold on firing-rate for determining optimal *b* is chosen such that at *α* = 0, the optimal balance level is the same as the one fitted to experimental data [20] in Fig. 3 (*B*^⋆^ ≈ 162). This threshold remains fixed for different values of *α*.

**Figure S3:** Throughout this figure, *α* = 0, *b* = 150.

**Figure S4:** Throughout this figure, *µ* = 0.97, *b* = 150.

**Figure S5:** The threshold value corresponding to the model curve is *θ* = −20. We set *b* = 189, which is the value extracted from the data [12].

**Figure S6:** Throughout this figure, we set *α* = 0, *b* = 150.

**Figure S7 and S8:** We set *θ* = 0, *J*^*II*^ = 0 throughout these figures. Before and after learning correspond to *µ* = 0 and *µ* = 0.97. Other parameter values are the same as in Fig. 5.

